# HOG pathway modulation is a determinant of echinocandin treatment failure in *Candidozyma auris*

**DOI:** 10.64898/2026.11.11.765954

**Authors:** Dimitrios Sofras, Stef Jacobs, Anitra Zīle, Austin Mottola, Alvis Zvirgzdins, Alicia Pageau, Giorgio Boccarella, Hugh Gifford, Vladislav Biriukov, Alexandre K. Dubé, Louise Walker, Dhara Malavia-Jones, Ian Leaves, Ziyue Wang, Bram Van den Bergh, Celia Lobo Romero, Pieter van den Berg, Jan Michiels, Rhys A. Farrer, Judith Berman, Neil A. R. Gow, Toni Gabaldón, Alons Lends, Markus Ralser, Carol A. Munro, Christian R. Landry, Patrick Van Dijck, Hans Carolus

**Affiliations:** Department of Biology, KU Leuven, Leuven, Belgium; National Institute of Research and Innovation, Riga, Latvia; Shmunis School of Biomedical and Cancer Research, Tel Aviv University, Tel Aviv, Israel; Department of Biochemistry, Charité – Universitätsmedizin Berlin, Berlin, Germany; Faculty of Medicine and Life Sciences, University of Latvia, Riga, Latvia; Département de Biochimie, de Microbiologie et de Bio-informatique, Faculté des Sciences et de Génie, Université Laval, Québec City, Canada; Institut de Biologie Intégrative et des Systèmes (IBIS), Université Laval, Québec City, Canada; PROTEO, Le regroupement québécois de recherche sur la fonction, l’ingénierie et les applications des protéines, Université Laval, Québec City, Canada; Centre de Recherche en Infectiologie (CRI), Université Laval, Québec City, Canada; Centre of Microbial and Plant Genetics (CMPG), KU Leuven, Leuven, Belgium; VIB Center for Microbiology, VIB, Leuven, Belgium; Medical Research Council Centre for Medical Mycology at the University of Exeter, Exeter, United Kingdom; Institute for Research in Biomedicine, Barcelona Institute of Science and Technology, Barcelona, Spain; Barcelona Supercomputing Center (BSC-CNS), Barcelona, Spain; School of Medicine, Medical Sciences and Nutrition, University of Aberdeen, Aberdeen, United Kingdom; CIBER de Enfermedades Infecciosas, Instituto de Salud Carlos III, Madrid, Spain; Catalan Institution for Research and Advanced Studies (ICREA), Barcelona, Spain; Max Planck Institute for Molecular Genetics, Berlin, Germany; KU Leuven One Health Institute, Leuven, Belgium

## Abstract

*Candidozyma auris* is a rapidly emerging fungal pathogen whose drug-resistant profile poses a formidable challenge to antifungal treatment. Echinocandins are the recommended first-line therapy for invasive *C. auris* infections, but resistance to this drug class is on the rise. Here we show that, beyond resistance, *C. auris* can readily acquire genetically encoded echinocandin tolerance, a phenotype of slow but sustained growth and increased survival at concentrations above the minimum inhibitory concentration (MIC) that reflects incomplete maximal inhibition and a reduced Hill coefficient without an increase in MIC. Tolerance was therefore undetectable by standard clinical susceptibility testing, yet it significantly reduced treatment efficacy in an immunocompromised murine model of systemic infection. Genetic dissection revealed that tolerance arises through disruption of the high-osmolarity glycerol (HOG) mitogen-activated protein kinase (MAPK) pathway through the loss of function of Ssk1, Ssk2, Pbs2 or Hog1, or specific substitutions in Nik1. The disruption of the HOG pathway did not confer fitness or virulence defects in *C. auris,* and in *Candida albicans* and *Nakaseomyces glabratus,* it did not confer tolerance, suggesting a unique non-essential rewiring of this pathway in *C. auris.* Proteomic and cell-wall analyses linked tolerance to altered cell-wall remodelling during echinocandin exposure. Evolutionary modelling and deep sequencing of *FKS1* hotspots in experimentally evolved populations showed that tolerance increases the rate at which resistance-conferring *FKS1* hotspot substitutions emerge. Putative high-impact mutations in HOG pathway effectors in clinical isolates suggest that this phenotype occurs in patients, where it might reduce treatment efficacy or facilitate resistance emergence. This work therefore identifies HOG-mediated acquired tolerance as a hidden clinically relevant direct and indirect determinant of echinocandin treatment failure.

## Introduction

*Candidozyma (Candida) auris* is a rapidly emerging fungal pathogen that has caused healthcare-associated outbreaks worldwide and presents a major challenge because of its frequent multidrug resistance^1,2^. Treatment of invasive fungal infections relies on four major antifungal classes: azoles, polyenes, echinocandins and the nucleoside analogue flucytosine^2,3^. Among clinical *C. auris* isolates, resistance is commonly reported at 77-91% for fluconazole, 12-24% for amphotericin B, and 1-12% for echinocandins^4,5^. Because of their favourable safety profile, high efficacy and relatively low resistance rates, echinocandins are recommended by the IDSA (Infectious Diseases Society of America) and CDC (US Centers for Disease Control and Prevention) as initial therapy for candidemia^6^.

Echinocandins inhibit Fks1, the catalytic subunit of the β-1,3-glucan synthase, an enzyme required for fungal cell-wall biosynthesis. Resistance is generally defined by a stable, heritable increase in minimum inhibitory concentration (MIC)^7^. Acquired echinocandin resistance primarily results from substitutions in three hotspot (HS) regions (HS1, HS2 and HS3) of *FKS1* that alter the drug target and increase the MIC^8,9^.

In 2025, the European Committee on Antimicrobial Susceptibility Testing (EUCAST) established species-specific clinical breakpoints for *C. auris* against anidulafungin (ANF) and micafungin (MCF) to define clinical resistance and guide treatment decisions, based on the MIC^10^. Routine antifungal susceptibility testing (AFST) determines the MIC only, and tolerance is generally not evaluated in susceptibility classifications or treatment decisions^7,14,15^. In this study, we define tolerance as slow growth at supra-MIC concentrations without an MIC increase^7^. Tolerance is distinct from heteroresistance, in which subpopulations have different MICs, and persistence, in which a small subpopulation survives drug exposure without growing, although inconsistent and overlapping definitions exist^7,11–13^.

Fungal tolerance has been studied predominantly for azoles, and azole tolerance has been described as slow growth of a subpopulation of otherwise susceptible cells^12,13,16^. Fluconazole tolerance has been associated with persistent candidemia, suggesting that this phenotype can influence treatment outcome, despite a below-breakpoint MIC^16^. Echinocandin tolerance has been studied far less extensively, and its genetic basis in

*C. auris* remains unclear^17,18^. Prior studies have suggested a link between the high-osmolarity glycerol (HOG) mitogen-activated protein kinase (MAPK) pathway (hereafter, HOG pathway) and echinocandin susceptibility in *C. auris,* with reports of both increased and decreased susceptibility in different assays^19,20^.

The HOG pathway is a conserved MAPK cascade that coordinates fungal adaptation primarily to osmotic (hence its name) and other environmental stresses^21,22^. In pathogenic *Candida* and other closely related species, signals from upstream histidine kinases, including Sln1 and Nik1, converge through Ypd1 and the response regulator Ssk1 on the Ssk2–Pbs2–Hog1 kinase module^23^. In the canonical phosphorelay model, phosphate transfer through Ypd1 keeps Ssk1 phosphorylated and inactive. When this transfer is interrupted, unphosphorylated Ssk1 activates the Ssk2–Pbs2–Hog1 kinase cascade. Beyond osmoadaptation, HOG pathway signaling contributes to oxidative and cell-wall stress responses and interacts with the cell-wall integrity (CWI) and Cek1 MAPK pathways^23,24^. In *C. auris*, Hog1 promotes stress tolerance and virulence, while Ssk1 and Hog1 influence antifungal susceptibility, cellular morphology, aggregation, cell-wall architecture and the activity of other MAPK pathways^19,20^. Together, these findings position the HOG pathway at the intersection of stress adaptation and cell-wall homeostasis^25,26^.

## Results

### Experimental evolution yields acquired echinocandin tolerance

To identify genetic paths of adaptation to echinocandins, we evolved 180 independent lineages of the clade I *C. auris* isolate B8441 (**Supplementary Table 1**) for 14 days at four concentrations of caspofungin (CAS), MCF or ANF, in liquid MOPS-buffered RPMI 1640 medium with serial culture dilution every 48 h (**Fig. 1a**). Six populations per drug concentration were randomly sampled, and one single-colony isolate from each population was tested in a broth dilution assay (BDA). Of the 72 isolates screened, 15 showed susceptibilities comparable to the parental wild-type (wt) strain and were excluded from further analysis. From the remaining isolates, we selected 25 drug-adapted strains based on decreased echinocandin susceptibility relative to the parental wt strain (resistance ratio based on the area under the curve (AUC), RR_AUC_ >1), while ensuring that the selection included isolates from all three echinocandins and all four drug concentrations used during experimental evolution. BDA results showed that, although relative 48-h MIC shifts (RR_MIC_) varied between the drug-adapted strains, RR_AUC_ and supra-MIC growth (SMG) ratios (TR_SMG_) were increased across most isolates (**Fig. 1b and Supplementary Fig. 1-2**). The high RR_MIC_ values for CAS in most strains are due to paradoxical growth, also known as the ‘Eagle effect’ in the BDA, which is known to occur in *C. auris* and complicates MIC determination by BDA^9^. The relative MIC (RR_MIC_) was calculated based on MIC measures of 50% growth reduction after 48 h of incubation, which is not standard for AFST (evaluated after 24 h of incubation). Increases in MIC in **Fig. 1b** therefore do not necessarily reflect standard resistance. To further discriminate tolerance from resistance, we therefore performed Etest susceptibility testing on the 25 adapted strains. Twenty-four retained an Etest MIC similar to that of the wt strain but showed increased growth within the inhibition ellipse after prolonged (48 h) incubation (**Fig. 1f** and **Supplementary Fig. 3 and Table 1**). Whereas classical resistance would shift the boundary of the inhibition ellipse towards higher drug concentrations (increased MIC), the increased supra-MIC growth indicated tolerance rather than resistance in 24 out of 25 evolved lineages. Growth rates and growth AUCs in drug-free conditions remained unchanged (**Fig. 1b**), suggesting that adaptation carried no detectable fitness cost (evaluated further in a later section) and that increased growth under echinocandin exposure was not due to adaptation to the culture conditions.

**Figure 1.**
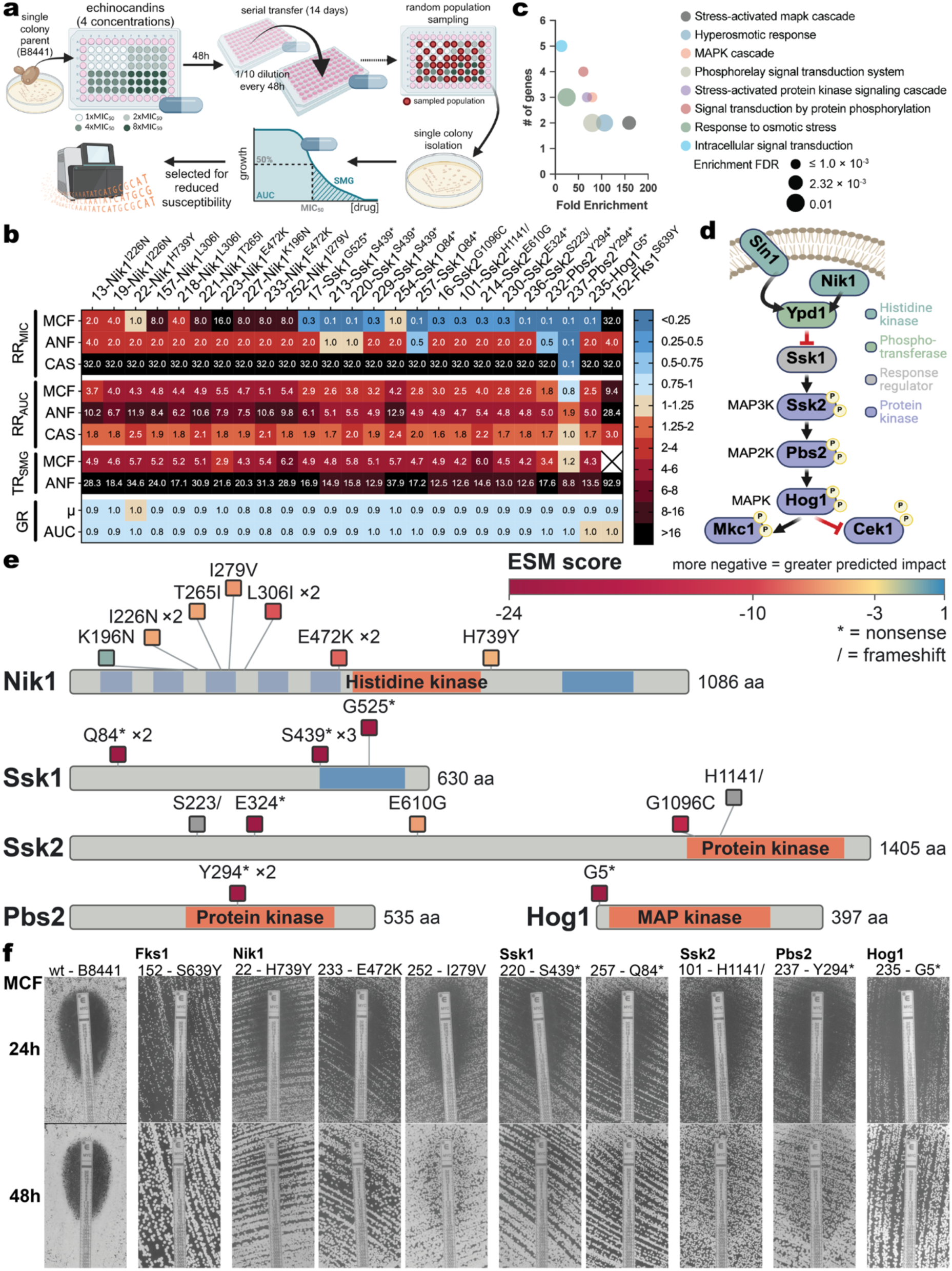
Experimental evolution identifies HOG pathway modulation as a genetic route to acquired echinocandin tolerance. **a)** Schematic representation of the workflow of experimental evolution, susceptibility screening and isolate selection for sequencing. **b)** Relative resistance ratio by MIC (RR_MIC_) and area under the curve (RR_AUC_), and tolerance ratio by supra-MIC growth (TR_SMG_), based on broth dilution assays (BDAs), for micafungin (MCF), anidulafungin (ANF) and caspofungin (CAS), together with the ratios of drug-free growth metrics (growth rate: µ and area under the growth curve: AUC) for the 25 sequenced isolates evaluated after 48 h of incubation. Relative ratios were calculated as 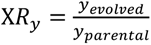 where X denotes the phenotype (resistance or tolerance) and y denotes the corresponding metric. TR_SMG_ could not be calculated for isolate 152 (*FKS1*^S639Y^), since no MIC was measured. **c)** Gene Ontology enrichment of the mutated genes from 25 drug-adapted strains, subjected to GO enrichment analysis using ShinyGO (v0.77). The x-axis indicates fold enrichment for each GO term, and the y-axis shows the number of genes assigned to each category. Dot size corresponds to enrichment significance (FDR-adjusted P value). **d)** Schematic representation of the *C. auris* HOG pathway based on literature^19,23,30^. **e)** Protein-level distribution of variants in Nik1, Ssk1, Ssk2, Pbs2 and Hog1 detected in 24 out of 25 sequenced adapted strains. Horizontal bars represent the proteins, with shaded regions indicating annotated domains and protein lengths given in amino acids (aa). Variant positions are indicated by labelled square markers, coloured according to their ESM scores: more negative scores indicate a greater predicted functional impact. Multipliers (×2 or ×3) indicate the number of isolates carrying the same variant. Asterisks (*) denote nonsense and slashes (/) denote frameshift variants. **f)** MCF Etests at 24 h and 48 h for the parental wt, resistant isolate 152 (Fks1^S639Y^) and a representative selection of evolved tolerant isolates carrying variants in Nik1, Ssk1, Ssk2, Pbs2 and Hog1. At least one isolate per mutated gene is shown, representing the shared phenotype observed across all tested evolved isolates carrying variants in that gene. Isolate 152 serves as the resistance comparator. Etests for all tested strains are provided in **Supplementary Fig. 3**.

The 25 adapted strains were sequenced (**Supplementary Table 2**) and Gene Ontology (GO) analysis of the complete mutated-gene set showed enrichment of stress-activated MAPK signaling, hyperosmotic response and protein-phosphorylation terms (**Fig. 1c**). Recurrently mutated genes mapped to the HOG pathway for 24 out of 25 mutated strains (**Fig. 1d,e**). Next, we estimated the functional impact of all HOG pathway variants with ESM1b, an evolutionary scale modelling (ESM) protein language model that captures evolutionary constraints from large protein-sequence datasets^27,28^. ESM scores represent the log-likelihood ratio of the variant versus the reference residue, with more negative scores indicating mutations that are less compatible with the protein sequence and therefore more likely to disrupt its function. We classified substitutions as predicted high impact using a loss of function (LoF) threshold derived from a two-component Gaussian mixture model fitted to the ESM-score distribution (**Supplementary Fig. 4**). Fourteen tolerant isolates carried a protein-altering variant in Ssk1, Ssk2, Pbs2 or Hog1, dominated by premature stop codons, frameshifts and high-impact substitutions, indicating LoF (**Fig. 1e**), whereas 10 isolates carried missense substitutions in Nik1, not predicted to confer a LoF (**Fig. 1e**). Only one sequenced strain (strain 152) showed a significant increase in MCF MIC (based on RR_MIC_ and Etest) and carried the known resistance-conferring substitution S639Y in Fks1 hotspot 1^29^.

### Loss of function of the HOG pathway confers echinocandin tolerance

Deletion of *SLN1*, *SSK1*, *SSK2*, *PBS2* or *HOG1* reproduced the tolerance phenotype seen in the experimentally evolved adapted strains (**Fig. 2a**). The *nik1Δ* strain did not show an increase in tolerance. The difference in susceptibility of the *nik1Δ* and the evolved *NIK1* missense variants, along with the predicted impact of *NIK1* mutations (**Fig. 1e**), suggests altered sensor regulation rather than LoF. These results show that the disruption of the HOG pathway, through the disruption or modulation of its upstream sensor kinases Sln1 and Nik1, respectively, or LoF of its response regulator Ssk1 or the Ssk2–Pbs2–Hog1 kinase cascade, causes echinocandin tolerance. Attempts to delete *YPD1* were unsuccessful, suggesting that it is an essential gene, as shown in several other fungi^31^. Without Ypd1, Ssk1 could stay unphosphorylated and Hog1 activation is constitutive, which is potentially lethal. We also investigate the roles of the related MAP kinases Mkc1 and Cek1 in echinocandin tolerance, by deleting them. The *mkc1Δ* strain showed increased MCF susceptibility, whereas deletion of *CEK1* did not alter echinocandin susceptibility relative to the wt strain (**Fig. 2a**). Mkc1 is the terminal MAP kinase of the cell wall integrity (CWI) pathway and is important for survival during echinocandin exposure^32^. The increased susceptibility of *mkc1Δ* and the tolerance of *hog1Δ* therefore indicate that the two MAPK pathways have opposing roles in the echinocandin response, with CWI signalling promoting survival and HOG pathway signalling limiting it. To test whether HOG-mediated tolerance depends on CWI signalling, we constructed a *hog1Δ mkc1Δ* double mutant, which was hypersusceptible to MCF, resembling *mkc1Δ* rather than the tolerant *hog1Δ* phenotype (**Fig. 2a**). This indicates that tolerance caused by HOG pathway disruption requires Mkc1-dependent CWI signalling.

**Figure 2.**
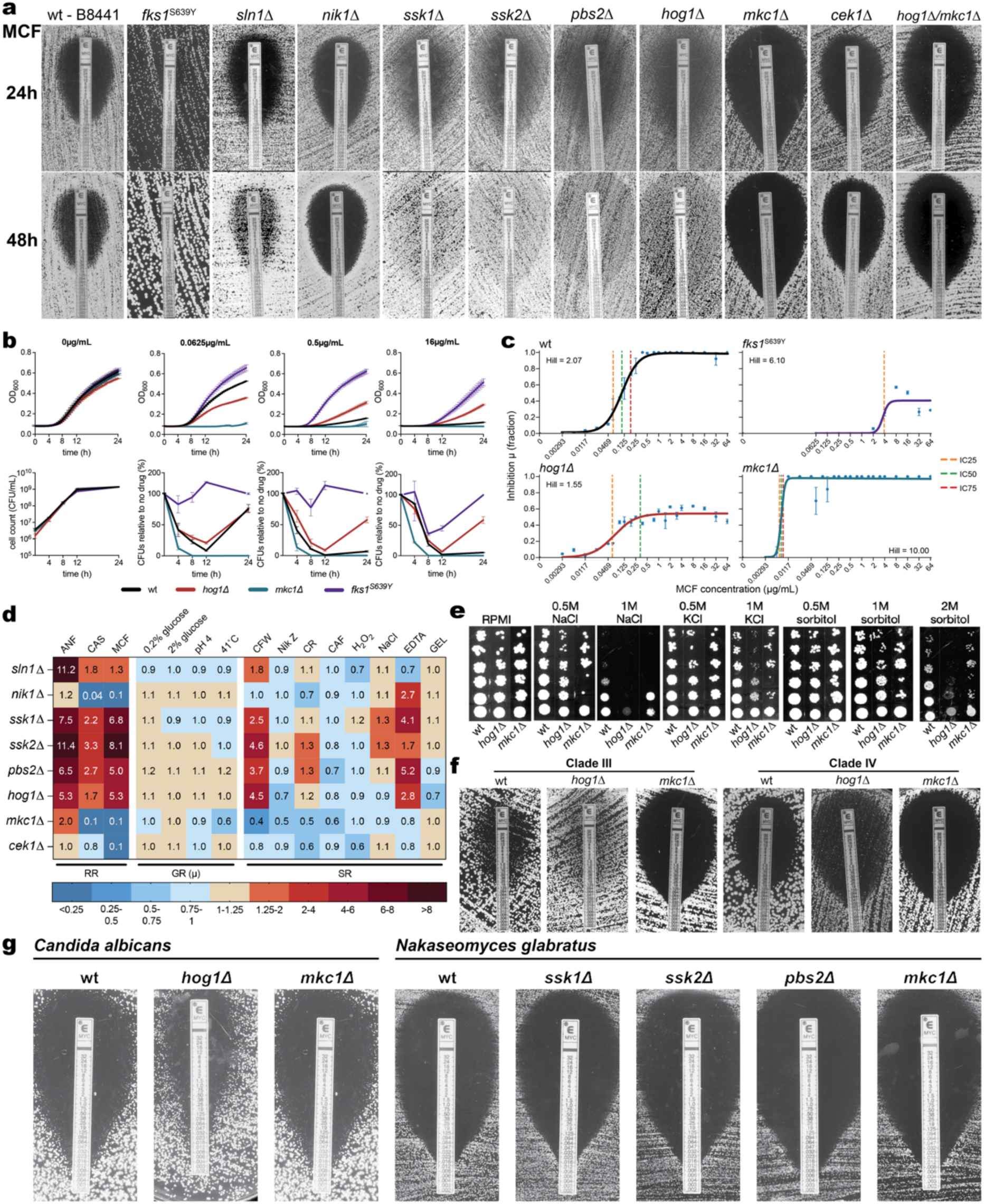
HOG pathway modulation produces a tolerant phenotype distinct from resistance. **a)** MCF Etests at 24 and 48 h for the wt and HOG pathway deletion strains, with the resistant Fks1^S639Y^ strain included as a resistance reference. **b)** Optical-density growth trajectories and CFU-based viable-cell counts for wt, *hog1Δ*, *mkc1Δ* and Fks1^S639Y^ during 24 h exposure to selected MCF concentrations (0, 0.0625, 0.5 or 16 µg mL^−1^ MCF). Viable-cell counts are shown as CFU mL⁻¹ in drug-free medium and as percentages relative to the corresponding drug-free controls during MCF exposure. The complete concentration gradient is shown in **Supplementary Fig. 5**. **c)** Concentration– inhibition curves fitted across the MCF gradient, showing inhibition of growth rate (µ) and the corresponding maximal inhibition and Hill coefficients for susceptible (wt), tolerant (*hog1Δ*), hypersusceptible (*mkc1Δ*) and resistant (Fks1^S639Y^) backgrounds. Points represent measured means of 3 replicates, error bars indicate the standard error of the mean (SEM, n=3) and solid lines indicate fitted curves. Dashed vertical lines indicate the concentrations producing 25%, 50% and 75% inhibition (IC₂₅, IC₅₀ and IC₇₅), where these levels are reached. **d)** Phenotypic-profile heatmap summarizing echinocandin resistance ratios (RR), drug-free relative growth-rate ratios (GR, µ) in different growth conditions and susceptibility ratios (SR) for various stressors. Colours and numbers indicate ratios relative to the wt under the corresponding condition. Relative ratios were calculated as 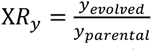 where X denotes the susceptibility metric measured under the corresponding stress condition. **e)** Osmotic-stress spot assays for wt, *hog1Δ* and *mkc1Δ* on RPMI-MOPS medium supplemented with the indicated concentrations of NaCl, KCl or sorbitol. Serial dilutions are shown from bottom to top. **f)** MCF Etests after 48 h for wt, *hog1Δ* and *mkc1Δ* strains generated in Clade III and IV *C. auris* genetic backgrounds. 24 and 72 h time points are shown in **Supplementary Fig. 6**. **g)** MCF Etests after 48 h for wt and the indicated HOG- and CWI-pathway deletion strains in *C. albicans* and *N. glabratus.* The 24 h time point is shown in **Supplementary Fig. 7**.

To determine whether these phenotypes extend beyond the Clade I background, we also deleted *HOG1* and *MKC1* in isolates belonging to the two other clinically most relevant clades: Clades III and IV. Consistent with results for Clade I, the deletion of *HOG1* produced the tolerance phenotype in both backgrounds (**Fig. 2f**). Deletion of *MKC1* abolished the basal tolerance observed in the Clade III wt strain but did not alter the Etest phenotype in Clade IV (**Fig. 2f**).

To test whether this mechanism of tolerance is conserved in other pathogenic *Candida* species, we evaluated MCF susceptibility by Etest in *Candida albicans hog1Δ* and *mkc1Δ* strains and *Nakaseomyces glabratus* (*Candida glabrata*) *pbs2Δ, ssk1Δ, ssk2Δ* and *mkc1Δ* strains. In *C. albicans,* the *hog1Δ* MIC was the same as in the wt strain although the halo was narrower, while *mkc1Δ* caused a fourfold reduction in MIC. In *N. glabratus,* the MIC of *ssk1Δ* and *ssk2Δ* was the same as that of wt, while *pbs2Δ* and *mkc1Δ* reduced the MIC twofold. Etests did not indicate increased tolerance in either species (**Fig. 2g**), suggesting that this mechanism is not conserved across the *Candida* phylogeny and possibly unique to *C. auris*.

Next, we further characterized the tolerant phenotype in *C. auris* by measuring colony-forming unit (CFU)-based survival, killing and growth of the wt, tolerant *hog1Δ*, hypersusceptible *mkc1Δ* and resistant Fks1^S639Y^ strains across an MCF concentration gradient (**Fig. 2b** and **Supplementary Fig. 5**). No significant growth or survival differences were observed in the absence of the drug. Across the MCF gradient, *hog1Δ* retained measurable growth at every tested concentration, exceeding wt at inhibitory and supra-MIC concentrations but remaining below that of the Fks1^S639Y^ strain. In contrast, *mkc1Δ* did not grow at any of the tested MCF concentrations (0.0625-16 µg/mL), including sub-MIC concentrations, with the MIC of the wt being 0.5 µg/mL (**Fig. 2b**). Killing curves showed that *hog1Δ* cells survived exposure to all tested concentrations of MCF, where the wt inoculum died off at MIC (0.5 µg/mL) and supra-MIC (16 µg/mL) concentrations. The resistant Fks1^S639Y^ strain showed the highest survival. Fitted growth-inhibition curves captured additional details of the drug dose response of the different strains: *hog1Δ* displayed a lower maximal level of inhibition and a reduced Hill coefficient compared to the wt (**Fig. 2c**), indicating that growth inhibition was incomplete and increased more gradually as MCF concentrations rose. In contrast, the Fks1^S639Y^ strain showed a clear increase in the fitted inhibitory concentrations (IC_25_, IC_50_ and IC_75_; **Fig. 2c**), whereas those of *hog1Δ* remained close to the wt values. The inhibitory concentrations or MIC indicate only where the drug begins to act, whereas the maximal level of inhibition and the Hill coefficient indicate how complete the effect becomes at clinically relevant supra-MIC concentrations. Isolates with identical MICs can therefore differ substantially in their response to the same exposure as tolerance illustrates: the MIC is unchanged, yet inhibition during drug exposure is reduced.

To test whether acquired tolerance was associated with any fitness costs, we measured the growth rate (µ, h^-1^) of wt, *hog1Δ, mkc1Δ* and Fks1^S639Y^ in several conditions and determined the relative (to wt) susceptibility to a range of stressors in BDAs in the panel of constructed deletion strains. Overall, no effect on fitness and no major decrease in stress resistance were detected (**Fig. 2d**). The HOG pathway is known for its role in osmotolerance^33^, and several deletion strains (*sln1Δ, nik1Δ, ssk1Δ, ssk2Δ* and *pbs2Δ)* showed a slight increase in tolerance to NaCl, while the *hog1Δ* strain was slightly more susceptible (**Fig. 2d**). We further assessed osmotic-stress responses using spot assays, which confirmed that *hog1Δ* is more sensitive to high concentrations of NaCl, KCl or sorbitol, compared to the wt (**Fig. 2e**). This finding agrees with the general protective role of Hog1 against osmotic stress, albeit only in extreme, clinically non-relevant conditions. In other species, the HOG pathway has been shown to be important for general fitness and stress tolerance^33^. Interestingly, HOG pathway deletion mutants showed increased stress resistance to the cell wall stressors calcofluor white (CFW), which binds chitin, and Congo red (CR), which binds β-glucans. In contrast, *mkc1Δ* showed reduced tolerance to both compounds. The tolerance to the divalent cation chelator ethylenediaminetetraacetic acid (EDTA) was also increased in HOG pathway deletion strains (except for *sln1Δ*). Divalent cation chelation compromises cell wall homeostasis, because Ca^2+^/calcineurin signalling and cation-dependent biosynthetic enzymes are required for the compensatory cell wall remodelling that maintains integrity under cell wall stress^34^. Taken together, these data suggest that HOG pathway disruption alters the response to cell-wall stress and that altered cell wall organization may underlie the mechanism of echinocandin tolerance.

### Drug-tolerant cells display a distinct cell-wall remodelling response

To investigate the cellular basis of echinocandin tolerance, we compared cell-wall composition and proteomic responses in wt and *hog1Δ* cells in the absence and presence of an echinocandin. We used CAS because MCF suppressed wt growth too strongly to yield sufficient biomass for these analyses.

In wt, CAS exposure significantly increased mannosyl-residue- and chitin-associated fluorescence by 2.7-fold and 7.9-fold, respectively, without a detectable increase in the β-glucan signal (**Fig. 3a**). In *hog1Δ*, CAS significantly increased β-glucan- and chitin-associated fluorescence by 1.7-fold and 2.6-fold, respectively, whereas the mannosyl-residue signal did not increase. These distinct staining profiles may reflect differences in the abundance or accessibility of cell-wall components. High-pressure ion chromatography (HPIC) analysis of hydrolysed cell wall polymers revealed changes in the relative proportions of mannan, glucan and chitin, inferred from mannose, glucose and glucosamine quantities, respectively (**Fig. 3b**). Following CAS exposure, the mannan proportion increased from approximately 25% to 45% in wt and from 30% to 65% in *hog1Δ*, while glucan decreased from approximately 70% to 50% and 30%, respectively. Chitin increased from below 1% to approximately 5% in both strains. These shifts accompanied distinct structural responses. Transmission electron microscopy (TEM) showed that in the wt, CAS exposure produced pronounced wall thickening by approximately 100 nm, most visibly in the inner glucan-chitin-rich layer. In contrast, *hog1Δ* showed no detectable increase in overall wall thickness but displayed a more prominent and dense fibrillar outer layer, consistent with its greater relative mannan proportion (**Fig. 3d**). Solid-state nuclear magnetic resonance (ssNMR) spectroscopy was used to identify the carbohydrate composition of intact fungal cells^35^ (**Supplementary Fig. 8**) and enabled quantification of carbohydrate components in rigid and mobile fractions (**Fig. 3c** and **Supplementary Fig. 8a-f**). Two-dimensional DARR spectra identified β-1,3-glucan, β-1,6-glucan, chitin and α-1,2-mannan. Complementary DP INADEQUATE spectra identified β-1,3-, β-1,6- and β-1,3,6-glucans, α-1,2- and α-1,6-mannans, galactose and glucose. These assignments were broadly consistent with previous ssNMR characterisation of untreated and CAS-exposed wt *C. auris* cells^36^. At least two chitin polymorphs were detected in all samples, while weak chitosan signals were additionally observed in *hog1*Δ cells. Furthermore, ¹H–¹⁵N cross-polarisation spectra indicated rigid protein components with amide ¹⁵N chemical shifts similar to those of chitin NH groups (**Supplementary Fig. 8g,h**). In both wt and *hog1Δ*, CAS exposure shifted the relative composition of the rigid fraction from β-1,3-glucan towards chitin. This broadly paralleled the decrease in glucose and increase in glucosamine measured by HPIC, although relative ssNMR signals do not quantify total polymer abundance or wall expansion. The CAS-treated *hog1Δ* cell wall was distinguished most clearly in its dynamic fractions. β-1,6-glucan signals were depleted from the mobile and highly mobile pools, while highly mobile β-1,3-glucan increased after treatment in *hog1Δ* but not in the wt (**Fig. 3c**). These changes are consistent with altered glucan packing or organisation and may help explain the increased Fc–Dectin-1 signal in *hog1Δ*, despite the decrease in its overall glucan proportion measured by HPIC. Mannan showed comparatively little redistribution between the ssNMR mobility fractions (**Fig. 3c**). Integrating these datasets, we propose that wt cells respond to CAS with pronounced wall thickening and increased surface exposure of mannan and chitin, whereas *hog1Δ* cells adopt a distinct reorganised wall state characterised by enhanced mannan and a selective redistribution of glucan mobility (**Fig. 3f**).

**Figure 3.**
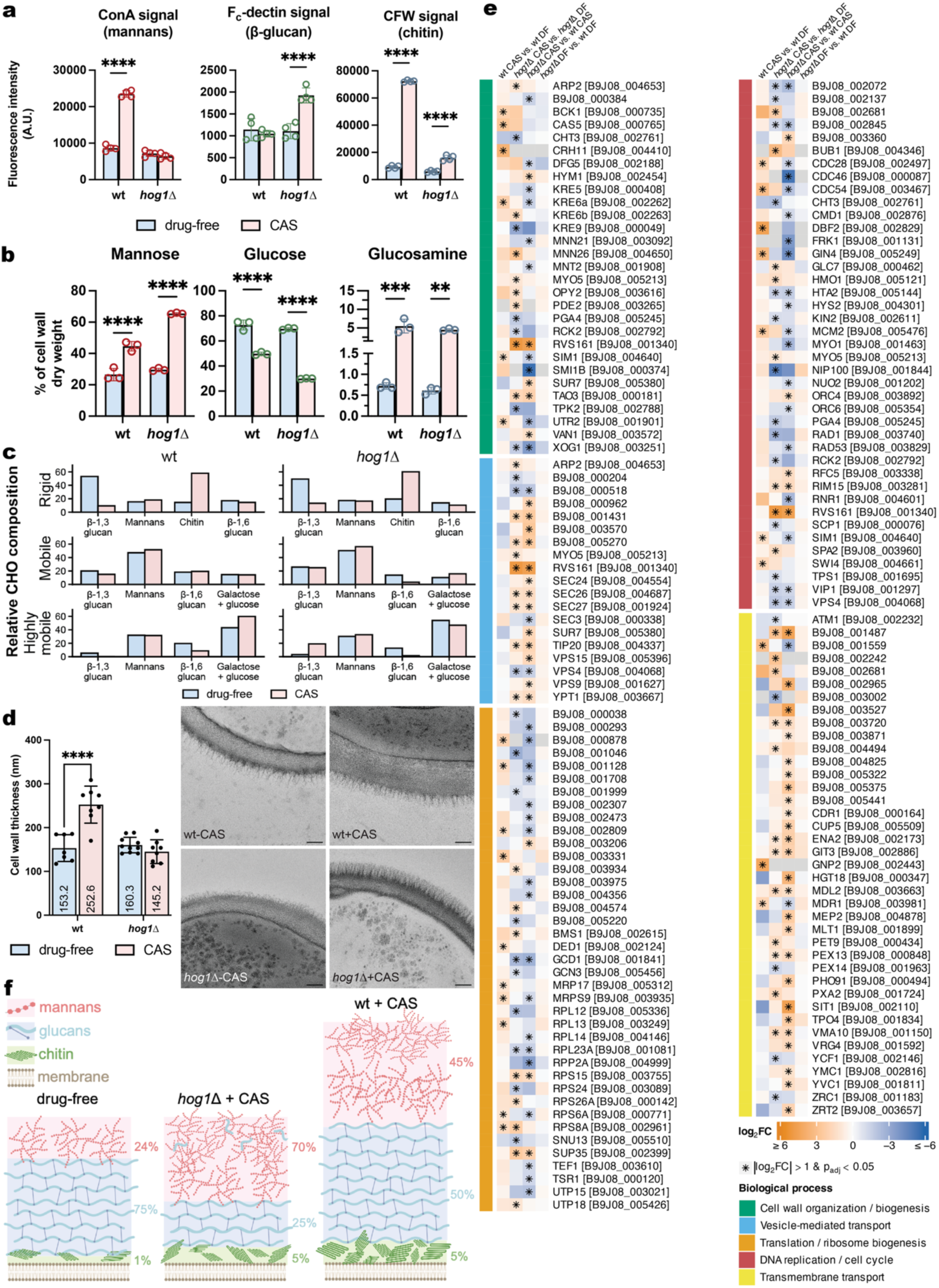
HOG-associated tolerance coincides with an altered cell-wall state and proteomic response. **a)** Flow-cytometric quantification of concanavalin A (ConA), Fc–Dectin-1 and calcofluor white (CFW) fluorescence, reflecting accessible mannosyl residues, β-glucan and chitin, respectively, in wt and *hog1Δ* cells grown with or without CAS. Fluorescence intensity is expressed in arbitrary units (A.U.). **b)** HPIC quantification of mannose, glucose and glucosamine in hydrolysed cell walls, used to estimate the relative mannan, glucan and chitin contents, respectively, and expressed as a percentage of cell-wall dry weight. **c)** Solid-state NMR-derived distribution of assigned carbohydrate (CHO) signals among rigid, mobile and highly mobile fractions of wt and *hog1Δ* cell walls grown with or without CAS. Values represent the relative distribution of signals within each mobility fraction and do not measure absolute polymer abundance. Assigned ¹³C chemical shifts are listed in **Supplementary Table 3**. **d)** Representative transmission electron micrographs and quantification of cell-wall thickness under the same conditions. Numbers within bars indicate mean thickness in nanometres. Scale bars are 100 nm. **e)** Heatmaps showing differential protein abundance for the treatment responses within each genotype (wt CAS versus wt drug-free, *hog1Δ* CAS versus *hog1Δ* drug-free) and the genotype differences under CAS-treated and drug-free conditions (*hog1Δ* CAS versus wt CAS, and *hog1Δ* drug-free versus wt drug-free, respectively). Colours indicate log₂ fold change, with orange and blue representing increased and decreased abundance, respectively. Asterisks indicate proteins with |log₂ fold change| >1 and adjusted P<0.05. Proteins are grouped by the indicated biological processes. DF denotes drug-free conditions. **f)** Schematic summary of cell-wall composition under drug-free conditions and following CAS exposure in wt and *hog1Δ,* based on flow cytometry, HPIC, ssNMR and TEM. For **a)**, **b)** and **d)**, bars represent means ± standard deviation (SD), with each data point representing the mean of three independent biological replicates for **a)** and **b)**, and seven cell wall measurements taken from micrographs of single cells for **d)**. Statistical significance was assessed by two-way ANOVA (n=7), followed by Šídák’s multiple-comparisons test, comparing drug-free and CAS-treated conditions within each genotype. **P < 0.01, ***P < 0.001 and ****P < 0.0001.

Proteome-wide responses to CAS differed markedly between wt and *hog1Δ* cells (**Fig. 3e** and **Supplementary Table 4**). CAS exposure significantly altered the abundance of 89 proteins in the wt (68 increased and 21 decreased), compared with 328 in *hog1Δ* (145 increased and 183 decreased). Under CAS exposure, 437 proteins differed significantly between the strains, with 241 more abundant and 196 less abundant in *hog1Δ* than in wt. Thus, the tolerant mutant showed more extensive proteomic changes than wt during drug exposure, whereas limited significant differences between the strains were detected under drug-free conditions. CAS exposure significantly increased the abundance of 6 cell wall-associated proteins in the wt, including the putative β-1,6-glucan synthesis protein Kre6a, the predicted chitin-associated glycosyltransferases Utr2 and Crh11 (B9J08_004410), together with the kinase Bck1 (cell-wall integrity pathway MAP kinase kinase) and transcription factor Cas5 (cell wall homeostasis regulator) (**Fig. 3e**). In *hog1Δ,* 10 cell wall proteins were less abundant than in wt under CAS exposure. These included Kre6a and Utr2, together with the putative exo-β-1,3-glucanase Xog1, the predicted β-1,3-glucanosyltransferase B9J08_000384 and the β- glucan biosynthetic protein Kre5. The same subset contained the mannoprotein Dfg5 and two predicted Golgi α-1,2-mannosyltransferases, Mnt2 (B9J08_001908) and Mnn21 (B9J08_003092), extending the differences beyond glucan and chitin remodelling. Van1, a predicted Golgi α-1,6-mannosyltransferase involved in mannoprotein glycosylation, showed the opposite pattern, being more abundant in *hog1Δ* than in wt under CAS exposure. Thus, the tolerant mutant displayed lower abundance of a functionally diverse set of cell wall proteins, including proteins that increased during the wt response to CAS. In addition, *hog1Δ* showed significantly higher abundance of 13 vesicle-mediated transport and 26 transmembrane transport proteins than wt under CAS exposure. These included Vrg4, which is predicted to transport GDP-mannose from the cytoplasm to the Golgi lumen, linking the transport response to mannosylation. Differences also extended to translation and DNA replication-associated proteins (**Fig. 3e**). In wt, CAS significantly increased 10 translation and 7 DNA replication proteins, whereas 18 translation and 19 DNA replication proteins were less abundant in *hog1Δ* than in wt under CAS. Together, these findings suggest that, under CAS exposure, the tolerant mutant adopts a different balance between biosynthetic and transport functions.

### Tolerance reduces treatment success and likely occurs in clinical isolates

The phenotypic assays we used so far profiled drug susceptibility under constant drug exposure. To evaluate the effect of tolerance during clinically relevant drug exposure, we assessed the population dynamics of tolerant and susceptible strains in a custom bioreactor which mimics pharmacokinetic drug exposure. We developed a perfusion bioreactor that mimics dynamic echinocandin exposure while simulating population loss through culture dilution (**Fig. 4a**). For this and other experiments that follow, we compared the wt strain with the tolerant Nik1^E472K^ mutant, since both have the same MIC, unlike the *hog1Δ,* which has a slightly decreased MIC compared to the wt strain (see **Fig. 2a**), which can confound the interpretation of results. Mixed populations of a tagged susceptible parent (wt with a nourseothricin resistance cassette in a neutral locus^37^) and the tolerant Nik1^E472K^ or resistant Fks1^S639Y^ mutant were grown with daily dynamic MCF administrations consisting of a 16 µg/mL peak (C_max_) and mono-exponential decay mimicking the pharmacokinetics of MCF in the human body^38^ (**Fig. 4a**, absolute burden in **Supplementary Fig. 9**). When introduced at 1% of the population, the resistant strain completely outcompeted wt, whereas the tolerant strain nearly displaced wt, with a small wt subpopulation of 3.75% ± 3.89 remaining at 72 h (**Fig. 4a**). Fks1^S639Y^ approached fixation within the first dosing interval (24 h), while Nik1^E472K^ reached a mean relative population share of 94.15% ± 3.47 after two drug administrations (48 h). Both tolerant and resistant strains completely outcompeted wt within 72 h when introduced at a 1:1 ratio (**Supplementary Fig. 10**). This demonstrates that tolerance can, like resistance, provide a direct competitive advantage under a clinically inspired drug exposure regimen, although displacement of wt occurred more slowly for the tolerant strain compared to the resistant one.

**Figure 4.**
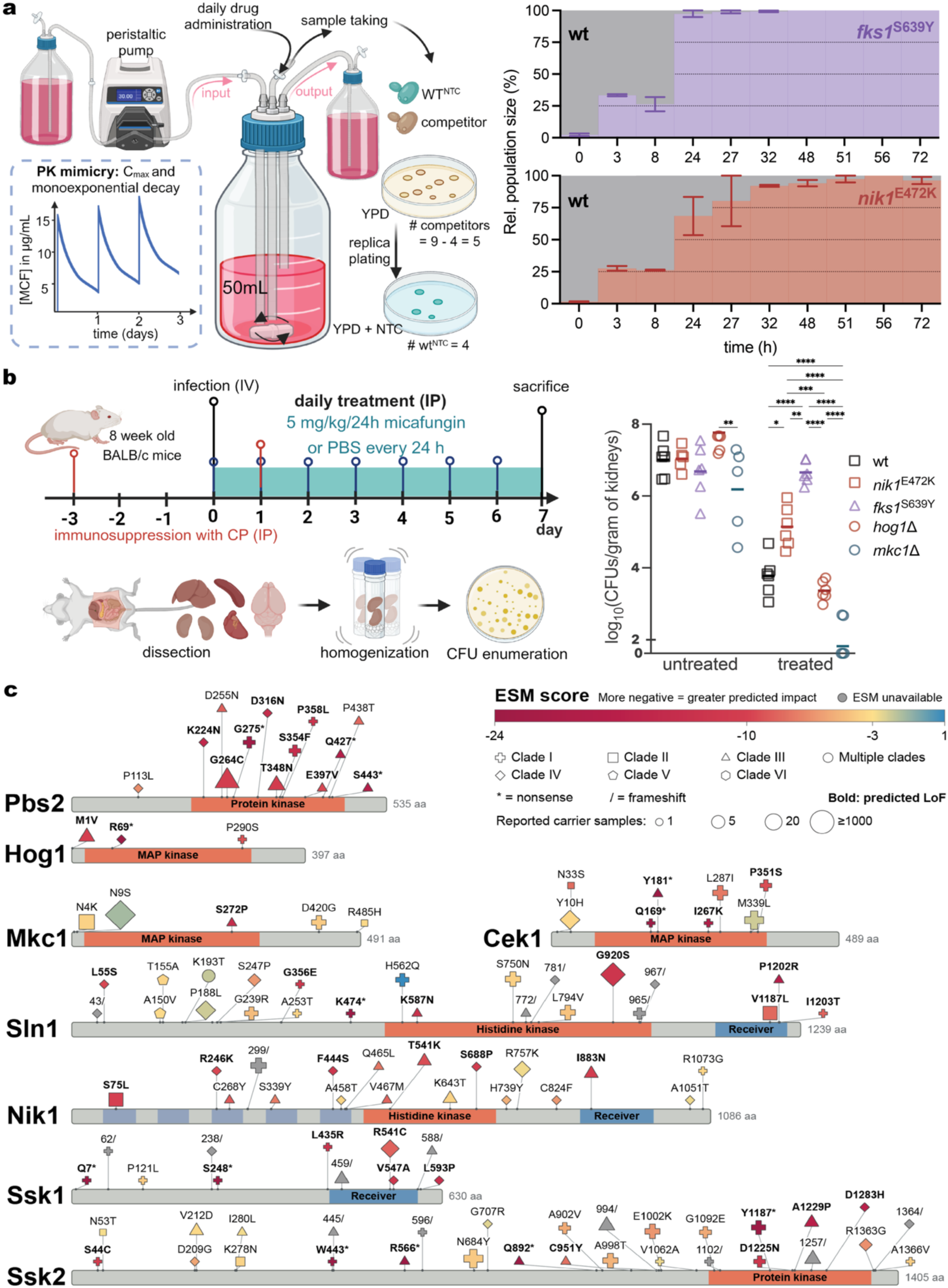
Tolerance is selected during treatment and represented in clinical populations. **a)** Schematic of the pharmacokinetic perfusion bioreactor reproducing once-daily MCF exposure and mono-exponential drug decay. Competitions were initiated with Fks1^S639Y^ or Nik1^E472K^ at 1% abundance with 99% wt, tagged with NAT to confer nourseothricin (NTC) resistance for differential plating, and sampled over 72 h. Relative population sizes were determined by plate replication of CFUs on YPD with or without NTC. Stacked plots show the relative abundance of wt (grey) and the resistant (purple) or tolerant (red) competitor over time. Data represent means ±SEM from 2 independent experiments per strain, per starting ratio. **b)** Experimental workflow and kidney burdens of susceptible wt, tolerant (Nik1^E472K^ and *hog1Δ*), resistant (Fks1^S639Y^) and hypersusceptible (*mkc1Δ*) strains recovered from untreated or MCF-treated immunosuppressed mice. Eight-week-old BALB/c mice were immunosuppressed with cyclophosphamide (CP) by intraperitoneal (IP) injection and infected intravenously (IV) on day 0. Mice received daily IP injections of MCF (5 mg kg⁻¹) or phosphate-buffered saline (PBS), and organs were collected on day 7 for CFU enumeration. Symbols represent individual animals and horizontal lines indicate means (n=6). Statistical significance was assessed by two-way ANOVA of log₁₀-transformed kidney burdens, with genotype and treatment as factors, followed by Šídák’s multiple-comparisons test. *P <0.05, **P <0.01, ***P <0.001 and ****P <0.0001. **c)** Distribution of epidemiological protein variants across Sln1, Nik1, Ssk1, Ssk2, Pbs2, Hog1, Mkc1 and Cek1 in the global 12,644-genome *C. auris* collection^39^. Protein domains are indicated, with protein lengths given in amino acids (aa). Marker colour represents the ESM score, with more negative values indicating greater predicted functional impact. Grey indicates that no score was available. Marker shape denotes clade, with circles indicating variants reported in multiple clades, and marker size represents the number of samples carrying the variant, capped at ≥1,000. Bold labels indicate variants with ESM scores below the corresponding gene-specific threshold. ESM score distributions and thresholds for each protein are shown in **Supplementary** Fig. 4. Asterisks (*) denote nonsense variants and slashes (/) denote frameshift variants.

We next tested whether the competitive advantage of the tolerant Nik1^E472K^ mutant over wt during MCF exposure holds in a mammalian host. Immunocompromised mice were infected with the wt, the tolerant *hog1Δ*, the tolerant Nik1^E472K^, the hypersusceptible *mkc1Δ* or the resistant Fks1^S639Y^ strain, followed by daily MCF treatment (**Fig. 4b**). Comparing the tolerant Nik1^E472K^ strain with the wt (same MIC) shows that Nik1^E472K^ caused a significantly higher fungal burden in the kidneys, compared to the wt, during MCF treatment (**Fig. 4b**). We also tested the tolerant *hog1Δ* and susceptible *mkc1Δ*, which shared the same MIC (lower MIC than the wt): during MCF treatment, *hog1Δ* led to a significantly higher fungal burden compared to *mkc1Δ*, again confirming that tolerance reduces treatment efficacy, independent of the MIC. However, the burden of *hog1Δ* did not differ significantly from the wt and was significantly lower than that of Nik1^E472K^, indicating that MIC, and thus relative resistance, remains important alongside tolerance during treatment. The resistant Fks1^S639Y^ strain showed significantly higher burden and *mkc1Δ* led to a significantly lower burden compared to all other tested strains. In untreated mice, *hog1Δ* showed higher burden than *mkc1Δ*, but no significant differences were observed between the other tested strains (**Fig. 4b**, other organs shown in **Supplementary Fig. 11**). The absence of a significant reduction in untreated fungal burden in the kidneys relative to wt indicates that HOG-pathway-deficient strains have no major virulence deficiency, as they retained the capacity to establish a systemic infection in this model.

To determine if potentially disruptive variants in HOG pathway genes occur in clinical *C. auris* isolates, we screened the global dataset of Gifford *et al.*^39^, comprising 12,644 genomes of *C. auris* collected between 1997 and 2024, and representing all six currently recognised clades. We considered epidemiological samples only and excluded laboratory-associated samples or samples of unknown origin. We extracted protein-altering variants in Sln1, Nik1, Ssk1, Ssk2, Pbs2 and Hog1, together with the related MAP kinases Mkc1 and Cek1 as controls. To limit the contribution of clade-specific variation, variants present in ≥95% of a given clade were arbitrarily excluded. We mapped missense, nonsense and frameshift variants onto the corresponding protein sequences and identified 110 distinct protein changes: 23 in Sln1, 18 in Nik1, 11 in Ssk1, 29 in Ssk2, 13 in Pbs2, 3 in Hog1, 5 in Mkc1 and 8 in Cek1 (**Fig. 4c** and **Supplementary Table 5**). Thus, most variation occurred in the upstream sensing and phosphorelay components of the HOG pathway. To estimate the potential functional impact of missense and nonsense variants, we used ESM-scoring^27,28^, which could not be used for frameshifts. Frameshifts were classified separately as putatively high-impact variants if the frameshift occurred in or upstream of a functional domain, as such variants likely result in LoF. Our analysis shows that 60 out of 110 detected variants (54.5%) are putative LoF variants. In the genes that we have shown to cause tolerance upon deletion (*SLN1, SSK1, SSK2, PBS2* and *HOG1*), 48 out of 79 variants (60.8%) were putative LoF variants, while putative LoF variation in *MKC1, CEK1* and *NIK1* was lower, occurring in 12 out of 31 (38.7%). This suggests that in the clinic, LoF variation in genes that confer putative echinocandin tolerance is more likely to occur, potentially due to the selective advantage during echinocandin treatment (**Fig. 4b**) and absence of fitness or virulence trade-offs (**Fig. 2d** and **Fig. 4b**). The number of putative LoF variants was especially high for *PBS2* (10 out of 13; 77%)*, SSK1* (9 out of 11; 82% excluding the terminal frameshift at position 588 as high impact)*, SSK2* (14 out of 29; 48% excluding the terminal frameshift at position 1364 as high impact) and *SLN1* (13 out of 23; 57%). High-impact variants *PBS2*^G264C^ and *SLN1*^G920S^ were detected most frequently, in 38 and 78 isolates, respectively. Although we did not detect mutations in *SLN1* in our experimentally evolved lineages (**Fig. 1e**), our genetic dissection shows that *sln1Δ* confers tolerance without decreasing the MIC (cf. *ssk1Δ, ssk2Δ, pbs2Δ* and *hog1Δ*; **Fig. 2a**). The lowest incidence of putative LoF variation was found in *MKC1 (1* out of 5, 20%, **Fig. 4c**), suggesting that the fitness trade-offs and hypersusceptibility to echinocandins (**Fig. 4a-d**) reduce fixation of LoF mutations in patients.

### Tolerance facilitates *FKS1*-mediated resistance evolution

Tolerance provides a competitive advantage both *in vitro* (**Fig. 4a**) and *in vivo* (**Fig. 4b**) where it enlarges the effective population size. This could increase the probability for resistance-conferring *FKS1* hotspot mutations to arise. To test this hypothesis, we experimentally evolved susceptible wt and tolerant Nik1^E472K^ populations for seven days under static, stepwise-doubling or stepwise-halving MCF regimens initiated across a range of starting drug concentrations (**Fig. 5a,b**). DNA was extracted from endpoint populations, and variation in the *FKS1* hotspot regions was assessed by deep sequencing. *FKS1* hotspot variants emerged in 6.5% (6/92) of susceptible wt populations and 23.9% (22/92) of tolerant Nik1^E472K^ populations (**Fig. 5b**). No mutations arose in the drug-free populations. The most frequently identified mutation was the HS1 variant Fks1^S639Y^ (detected in 21/92 Nik1^E472K^ populations and 1/92 wt populations under drug selection), followed by the HS1 variant Fks1^S639P^ (1/92 populations of both Nik1^E472K^ and wt) and the HS3 variant Fks1^W691L^ (4/92 wt populations). All three mutations are known to confer echinocandin resistance in *C. auris*^40–43^. Across starting concentrations, tolerant populations had a significantly higher probability of resistance under static and 72-h doubling exposure (**Fig. 5c**). Across the 23 conditions, resistance emerged at least as frequently in tolerant as in wt populations, except under 24-h doubling and halving initiated at 4× MIC. The difference was greatest under static exposure, with no clear dependence on starting concentration. Under doubling and halving regimens, the lowest starting concentrations yielded the most resistance-positive populations. These findings support the hypothesis that pre-existing tolerance promotes *FKS1*-mediated resistance emergence during MCF treatment.

**Figure 5.**
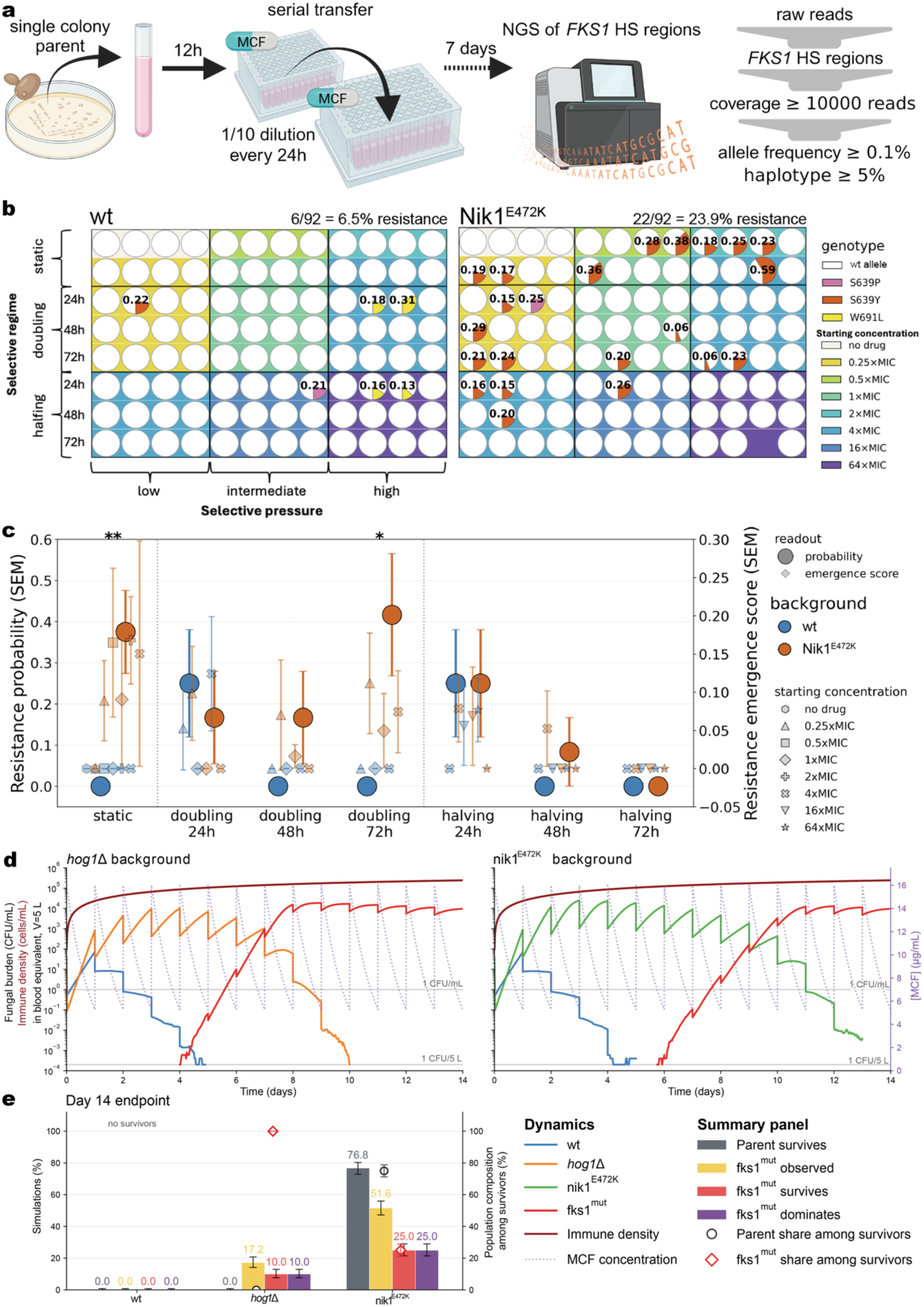
Tolerance promotes *FKS1*-mediated resistance evolution. **a)** Clonal wt or tolerant Nik1^E472K^ mutant populations were serially transferred for 7 days under MCF exposure, with a 1/10 dilution every 24 h. *FKS1* hotspot regions were analysed by deep sequencing using a minimum coverage of 10,000 reads, an allele-frequency threshold of 0.1% and a haplotype-frequency threshold of 5%. Static, stepwise-doubling and stepwise-halving regimens were initiated across a range of starting concentrations and selective pressures. For stepwise regimens, concentrations were doubled or halved every 24, 48 or 72 h. **b)** Each circle represents one independently evolved population, with coloured segments of the pie charts and numbers indicating the detected *FKS1* variant and its allele frequency, expressed as a fraction of total haplotypes. White segments indicate the wt allele of the three *FKS1* hotspots. Background colours indicate starting MCF concentrations relative to the parental MIC (MIC_wt_ = 0.5 µg/mL), grouped into low, intermediate and high selective pressure. Overall, *FKS1* variants were detected in 6/92 wt populations (6.5%) and 22/92 tolerant Nik1^E472K^ populations (23.9%). **c)** Large circles show the mean resistance probability summarised across starting concentrations (n = 20 for static and n = 12 for the doubling/halving regimes), whereas smaller symbols show emergence scores for the individual starting concentrations (n = 4). Colours indicate the strain (wt or tolerant Nik1^E472K^), symbol shapes indicate starting concentration, and error bars represent SEM. Resistance probability was compared between parental backgrounds within each exposure regimen using two-sided Fisher’s exact tests, and emergence scores were compared using two-sided Mann–Whitney U tests. *P < 0.05, **P < 0.01. **d)** Stochastic population model incorporating fungal growth, once-daily MCF exposure and immune-mediated clearance. Simulations were initiated separately with 2 × 10⁵ wt, *hog1*Δ or Nik1^E472K^ parental cells in a 5 L blood-equivalent compartment and followed for 14 days. MCF concentrations peaked at 16 µg mL⁻¹ every 24 h and declined with a 15-h half-life. Each dynamics panel shows one representative simulation per genotype with the blue wt trajectories representing separate simulations. Colours distinguish parental populations, emerging resistant lineages (Fks1^mut^), immune-effector density and MCF concentration. The resistant phenotype was parameterised using Fks1^S639Y^ as proxy. Horizontal reference lines indicate 1 CFU mL⁻¹ and one CFU in the entire 5 L compartment. **e)** Day-14 outcomes from an ensemble of 500 independently seeded simulations per parental background. Bars indicate the percentages of simulations in which the parental population survived (grey), an Fks1^mut^ lineage had been observed by day 14, or an Fks1^mut^ lineage survived or was dominant at day 14. Numbers above bars indicate percentages, and error bars show binomial Wilson 95% confidence intervals. Open circles and diamonds indicate mean parental and Fks1^mut^ population shares, respectively, among simulations with survivors, plotted against the right y-axis. Their error bars show 95% confidence intervals for the mean. Corresponding day-7 outcomes are shown in **Supplementary Fig. 13**.

To further explore how MCF tolerance could influence resistance development during treatment, we used a stochastic computational model informed by experimentally obtained growth and survival measurements and literature derived parameters. The model compared resistance emergence in wt and tolerant backgrounds under repeated MCF exposure and immune-mediated clearance (**Fig. 5d,e**, parameter details in **Supplementary Table 8**). The fitted concentration–response functions showed strong suppression of wt growth by MCF, whereas the tolerant strains retained substantially higher growth rates (**Fig. 2c**, **Supplementary Fig. 12**). At the simulated peak concentration of 16 µg mL⁻¹, fitted mean growth rates were 0.038 h⁻¹ for wt, 0.251 h⁻¹ for *hog1Δ* and 0.298 h⁻¹ for Nik1^E472K^, compared with 0.398 h⁻¹ for the resistant proxy. In separate simulations used to illustrate first-day recovery of each strain individually, fungal burdens after 24 h were approximately 71, 803, 902 and 1,611 CFU mL⁻¹ for wt, *hog1Δ*, Nik1^E472K^ and Fks1^mut^, respectively (**Supplementary Fig. 12**). Representative trajectories of the wt or tolerant populations and emerging resistant lineages within the same simulation illustrate how tolerant populations could expand between doses and generate resistant lineages before being cleared (**Fig. 5d**). Outcomes across all replicate simulations were summarised at days 7 and 14 (**Fig. 5e, Supplementary Fig. 13**). Both tolerant parental populations survived in 100% of simulations at day 7 (**Supplementary Fig. 13**). Thus, the subsequent clearance of the *hog1Δ* parent by day 14 does not contradict its recovery at the seven-day endpoint of the murine experiment, although the experimental and simulated host conditions differ. By day 14, the parental subpopulation died out in all simulations, and no resistant lineage emerged in the wt condition (**Fig. 5e)**. In the *hog1Δ* background, Fks1^mut^ lineages emerged in 17.2% of simulations and survived to day 14 in 10.0%. The *hog1Δ* parental population was cleared in every simulation, leaving the resistant lineage as the only surviving fungal population in those runs. In the Nik1^E472K^ background, the parent survived to day 14 in 76.8% of simulations, while an Fks1^mut^ lineage was observed in 51.6% of simulations. Resistant lineages survived and became numerically dominant in 25.0% of simulations. Here, dominance means that resistant cells outnumbered parental cells at the endpoint. It does not necessarily indicate complete replacement (**Fig. 5e**). Together, these simulations suggest that tolerance can increase opportunities for resistance evolution by maintaining fungal populations during treatment. The Nik1^E472K^ background supported prolonged parental survival, whereas *hog1Δ* provided a transient window during which resistance could emerge and sustain the infection after clearance of the tolerant parent.

## Discussion

Echinocandin resistance in *C. auris*, for which these drugs are first-line treatment^6^, mainly arises through mutations in Fks1 hotspots^9^, although mutations in *CRZ1*, which encodes a calcineurin-responsive transcription factor, have also been reported^44^. Here, *C. auris* evaded treatment without alterations in *FKS1* or *CRZ1* or an increased MIC. Increased growth and survival above an unchanged MIC are consistent with tolerance^7,11,12,45^. Although echinocandin tolerance has been described^46^, acquired tolerance through validated genetic mechanisms remains rarely reported, especially in clinical isolates. To date, out-of-hotspot *FKS1* mutations are the only validated mechanism of stable, mutation-based acquired echinocandin tolerance in clinical pathogenic fungi^47,48^. Narayanan *et al.* recently showed acquisition of tolerance through a small supernumerary chromosome in *C. auris*^49^, but aneuploidies are unstable and often carry fitness costs in the absence of selection^50^. Our findings identify stable, mutation-based HOG pathway changes that confer tolerance without detectable fitness or virulence trade-offs.

In the canonical HOG pathway, the sensor histidine kinases Sln1 and Nik1 regulate a phosphorelay through Ypd1 that keeps Ssk1 phosphorylated and inactive. Interrupting this allows dephosphorylated Ssk1 to activate the downstream Ssk2–Pbs2–Hog1 kinase cascade^21^; loss of Ssk1 or any of these kinases therefore disrupts signalling. The exclusively missense variants in Nik1 and absence of tolerance in *nik1Δ* instead suggest altered sensor regulation. We hypothesise that these substitutions may favour Ssk1 phosphorylation, suppressing Hog1 activation. Interestingly, *SLN1* deletion also conferred tolerance, despite the canonical model predicting Hog1 hyperactivation upon loss of Sln1, indicating that upstream HOG pathway regulation in *C. auris* may differ from other yeasts^51^. Finally, loss of tolerance in *mkc1Δ* and *hog1Δmkc1Δ* implies that tolerance depends on Mkc1, the terminal MAP kinase of the PKC-dependent cell-wall integrity (CWI) pathway^32^, and thus crosstalk between HOG and CWI pathways. This putative dependence is consistent with CAS-induced Mkc1 phosphorylation in *C. auris ssk1Δ* and *hog1Δ*^20^.

Loss of Hog1 appears to redirect rather than abolish the response to echinocandin-induced cell-wall damage. Although increased chitin synthesis supports echinocandin survival in *C. albicans* and *C. auris*^52,53^, neither greater relative chitin enrichment nor wall thickening explained *hog1Δ* tolerance. Balanced rather than maximal compensation has precedent in *C. albicans*, where calcineurin restrains Mkc1-dependent chitin accumulation and excessive chitin is associated with growth arrest and loss of viability^54^. Increased resistance to CFW and CR, but not the chitin synthesis inhibitor nikkomycin Z (NikZ), suggests selective protection against cell-wall perturbations rather than broadly enhanced stress resistance. Increased β-glucan accessibility despite a reduced glucan proportion, mannan enrichment without increased ConA binding, and altered ssNMR-defined glucan mobility fractions instead point towards changes in wall polymer organisation.

Reduced abundance of the β−1,6-glucan synthase Kre6a in CAS-treated *hog1Δ* could contribute to its reduced mobile β−1,6-glucan. Recently, *kreCaΔ* was shown to decrease *C. auris* echinocandin susceptibility, with the effect most evident in spot assays but hardly visible in liquid BDA^36^, consistent with the tolerance phenotype we describe here. *KRECa* deletion modestly but significantly increased *FKS1* expression, whereas overexpression had the opposite effect^36^. However, CAS restored the relative contribution of β-1,6-glucan to the mobile fraction in *kreCaΔ*^36^, contrasting with our findings and suggesting that reduced Kre6a alone cannot fully explain the *hog1Δ* wall phenotype. Lower abundance of the glycosyltransferase Utr2 and B9J08_000384, a putative β-1,3-glucanosyltransferase orthologous to *C. albicans* Phr2, suggests altered polymer incorporation and connectivity. In *C. albicans*, changing Crh-family protein abundance, including Utr2, alters wall elasticity and osmotic-stress survival, consistent with their proposed chitin–glucan cross-linking roles^55^. Thus, *hog1Δ* could retain chitin compensation while incorporating it differently into the glucan network, potentially explaining altered architecture without pronounced thickening. Reduced abundance of Xog1, an exo-β-1,3-glucanase, could explain greater Dectin-1 binding despite a lower glucan proportion: *C. albicans* Xog1 trims exposed β-glucan epitopes, so reduced activity can increase glucan accessibility without increasing abundance^56^. Significantly higher abundance of Van1, a putative α-1,6-mannosyltransferase, and Vrg4, a putative GDP-mannose transmembrane transporter, could contribute to the increased relative mannan proportion and more prominent fibrillar outer layer. Together, these changes suggest dampened cell-wall biosynthetic responses and selective reorganisation of wall assembly and connectivity, rather than uniformly increased repair or globally reduced biosynthesis. Whether Hog1 or Mkc1 regulates these effectors directly or indirectly, remain to be established.

HOG pathway disruption has been reported to both increase and decrease echinocandin susceptibility in pathogenic fungi, with differences in assay conditions and genetic backgrounds complicating comparisons. Traditional AFST measures the MIC, usually after 24 h, and would miss the tolerance observed here. Comparing 24- and 48-h Etests and measuring growth and survival across concentrations clearly distinguished tolerance from resistance, whereas the traditional BDA readout alone could not. Drug choice also matters: CAS-specific paradoxical growth in *C. auris*^9^ complicates this distinction. Moreover, increased sensitivity at low concentrations can coexist with enhanced growth and survival above the MIC, as our data show. This assay dependence helps reconcile increased growth of *C. auris hog1Δ* on CAS agar reported by Day *et al.*^19^ with decreased growth of *hog1Δ* and *ssk1Δ* during 24-h exposure to 0.1 µg/mL CAS reported by Shivarathri *et al.*^20^, resembling our low-concentration MCF response. The absence of increased tolerance in *C. albicans* and *N. glabratus* HOG-pathway mutants also agrees with prior findings. Alonso-Monge *et al.* found no effect of *C. albicans hog1Δ/Δ* on 24-h MCF MIC, growth on MCF agar or 1-h survival at a supra-MIC concentration of MCF^57^. Harrington *et al.* likewise detected no difference in survival of *N*. *glabratus hog1Δ, ssk2Δ* and *pbs2Δ* during 30-h exposure to 1 µg/mL MCF^58^. Loss of the Hog1 orthologue SakA in *Aspergillus fumigatus* increased CAS sensitivity and reduced high-dose paradoxical regrowth in liquid growth assays^59^. Together, these findings suggest that HOG-mediated gain of tolerance is unique to *C. auris*. Reitler *et al.* nevertheless reported increased growth at wt-inhibitory CAS concentrations in *C. albicans hog1Δ/Δ, pbs2Δ/Δ* and *ssk2Δ/Δ*, but their categorical screening phenotypes did not distinguish MIC shifts or quantitatively increased trailing growth from paradoxical growth in *hog1Δ/Δ*^60^. Polvi *et al.* experimentally evolved *C. albicans* under CAS and the metal chelator DTPA, obtaining *NIK1* substitutions that impaired drug-induced Hog1 activation and reduced susceptibility to both agents. They also observed increased CAS resistance and/or tolerance in *ssk1Δ/Δ, ssk2Δ/Δ*, *pbs2Δ/Δ* and *hog1Δ/Δ*, but not *nik1Δ/Δ*, in BDA^61^. These observations support a potentially broader capacity for HOG modulation to increase growth or survival during echinocandin exposure, although they do not all establish the MIC-independent, growth-associated phenotype described here.

Beyond susceptibility, an important determinant of whether HOG disruption can promote survival during treatment is its effect on fitness and virulence, which seems to be different for *C. auris* compared to other species. In *C. albicans*, *Cryptococcus neoformans* and *A. fumigatus,* abrogating HOG signalling compromises stress adaptation and fitness^23^. In *C. albicans, HOG1* loss attenuated systemic virulence, increasing mean survival from 1–3 to 30–60 days^62^, and compromised competitive gastrointestinal colonisation^63^ in mice. In *N. glabratus*, it impaired survival within macrophages and reduced fungal burdens after seven days of disseminated murine infection^64^. In contrast, we detected no fitness or virulence reduction in tolerant *C*. *auris* strains. Nevertheless, the sensitivity of *hog1Δ* to high osmotic stress reveals a condition-dependent fitness cost. Consistent with our findings, drug-free growth of *C. auris hog1Δ* and *ssk1Δ* mutants remained broadly similar to parental controls across several backgrounds^20^, and Hog1 was dispensable for growth at pH 4 and 8 and resistance to sorbic acid and nitrosative stress^19^. However, reduced virulence of *hog1Δ* was reported in nematodes^19^, and reduced renal and brain, but not splenic or hepatic, burdens were observed in immunocompetent mice after 72 h^30^. Differences in assays, strain backgrounds, virulence models, sampling times and host immunity limit comparison. Together, the evidence suggests that loss of HOG signalling carries lower fitness and virulence costs in *C. auris* than in other species, potentially indicating pathway rewiring or repurposing. This may reflect its ecological history: *C. auris* has been isolated from high-salinity tidal marshes and beaches^65^, and marine niches have been proposed for related *C. haemulonii*-complex species^66,67^. *C. auris* withstands cationic stress more effectively than *C*. *albicans* despite weaker Hog1 activation, although *HOG1* remains required under severe NaCl, KCl and sorbitol stress*^1S^*. We therefore speculate that adaptation to saline environments may have reduced the dependence of core fitness and stress adaptation on HOG signaling, unlike predominantly host-associated species such as *C. albicans* and *N. glabratus*, in which substantial fitness and virulence costs may limit pathway disruption during infection.

HOG-mediated tolerance threatens echinocandin efficacy directly by reducing treatment efficacy and indirectly by promoting resistance evolution. Putative LoF variants in tolerance-associated genes in clinical *C. auris* isolates suggest that this threat may be present in the clinic. The breadth of the genetic target for tolerance is particularly concerning: *SLN1, SSK1, SSK2, PBS2* and *HOG1* collectively encode 4,206 residues, with 31.36% (25,079 of 79,966) of variants predicted by ESM to cause LoF. By comparison, Durand *et al.* classified 26% (121 of 465) of variants as echinocandin resistant in a deep mutational scan of all amino acid substitutions across the 28 residues of the three *Saccharomyces cerevisiae* Fks1 hotspots^68^. Weighting target lengths by these fractions predicts an approximately (4,206 × 0.3136)/(28 × 0.26) ≍ 180-fold higher probability of generating tolerance-conferring variants than variants conferring resistance to echinocandins under an equal-mutability model. This crude estimate uses predicted LoF as a proxy for tolerance, the *S. cerevisiae* resistance fraction as a reference, and excludes *NIK1* mutations and tolerance-associated indels and frameshifts in the five HOG pathway-encoding genes. The actual probability of tolerance versus resistance fixation is therefore most probably higher. The central implication is that HOG disruption offers a substantially broader opportunity for drug adaptation than specific changes within short drug-target hotspots. This may explain the predominance of tolerant HOG mutants over *FKS1* mutants among evolved clones we selected for sequencing – initially just based on reduced susceptibility by RR_AUC_ > 1 and not based on tolerance. Our experiments and simulations show that tolerance facilitates resistance emergence. This facilitation also occurs during bacterial adaptation to antibiotics^69,70^. Our simulations suggest how this can occur in *C. auris*: enhanced survival and inter-dose proliferation preserve a larger reproducing population during daily MCF exposure, increasing opportunities for resistant descendants to arise and establish despite immune-mediated clearance. Resistant lineages could consequently sustain infections after the tolerant parent was eliminated.HOG-mediated tolerance is therefore not just an additional susceptibility phenotype, but a determinant of both treatment response and subsequent resistance risk.

We suggest to evaluate tolerance alongside resistance in clinical susceptibility testing of echinocandins in *C. auris*. Extending incubation beyond the 24-h MIC readout and measuring supra-MIC growth provides a practical approach to detecting tolerance by standard AFST. We suggest using MCF or ANF in traditional BDA and evaluating AUC and SMG after both 24 and 48 h. Alternatively, it can be detected by measuring trailing growth within the Etest inhibition ellipse after prolonged incubation, or the maximal inhibition and Hill coefficient from 24-h dose–response curves. Genomic surveillance should include *SLN1, NIK1, SSK1, SSK2, PBS2* and *HOG1* alongside resistance determinants such as *FKS1*, capturing changes that can influence both resistance and tolerance. Additionally, future work should further dissect the signalling and interaction network sustaining tolerance, test whether clinical HOG variants predict tolerance and are enriched in tolerant isolates, and link these associations to antifungal exposure and treatment outcome. Finally, the putative dependence on Mkc1 suggests a potential therapeutic vulnerability: Mkc1 inhibition or restoration of Hog1 activity could suppress tolerance and improve echinocandin efficacy. Combined, phenotypic detection, genomic surveillance and mechanistically informed interventions could improve echinocandin efficacy directly, in patients.

## Methods

### Ethical statement

All animal experiments were approved in accordance with the ethical guidelines of the Ethics Committee of KU Leuven (project numbers 079/2020 and 148/2022).

### Strains and growth conditions

The strains used in this study are listed in **Supplementary Table 1**. The clade I *Candidozyma auris* isolate B8441 was used as the parental wt strain for experimental evolution and CRISPR–Cas9 genome editing. Engineered mutants were additionally constructed in the clade III isolate B11222 and the clade IV isolate B11245. For cross-species comparisons, *C. albicans* GRACE conditional-expression strains (CaSS1::*hog1*Δ/p_TET_*HOG1* and CaSS1::*mkc1*Δ/p_TET_*MKC1*) and *N. glabratus* deletion mutants were obtained from the collections described by Roemer *et al.*^71^ and Schwarzmüller *et al.*^72^, respectively. Unless stated otherwise, cells were cultured at 37 °C in RPMI 1640 buffered with MOPS to pH 7.0 and supplemented with 2% glucose. Strains were maintained on YPD agar (1% yeast extract, 2% peptone and 2% glucose) and stored at −80 °C in 27% glycerol PBS. For the *C. albicans* strains, the medium was supplemented with 20 µg/mL doxycycline.

### Experimental evolution under echinocandin exposure

Experimental evolution was performed with B8441 in MCF, ANF or CAS following the same method described elsewhere^37,73^. For each drug, 15 independent populations were initiated at each of four concentrations (1×, 2×, 4× and 8× the wt MIC), yielding 60 lineages per drug and 180 lineages in total. Each population was founded with approximately 1 × 10^6^ cells in 200 µl RPMI–MOPS containing 2% glucose and 1% dimethyl sulfoxide (DMSO). Populations were transferred 1:10 into fresh medium containing the corresponding drug concentration every 48 h for six transfers over 14 d. Populations were archived after each transfer by adding glycerol to a final concentration of 27% and storing at −80 °C.

### Broth dilution assays and tolerance quantification

Antifungal susceptibility was assessed using broth dilution assays (BDAs). Twofold dilution series of MCF, ANF or CAS were prepared over a concentration range of 0.0625-16 µg mL⁻¹ in a final volume of 200 µl RPMI–MOPS containing 2% glucose and 1% DMSO. Wells were inoculated with 100-500 cells and incubated at 37 °C for 48 h. Growth was subsequently measured as optical density at 600 nm (OD_600_) using a Synergy H1 microplate reader (BioTek). The initial screen of evolved isolates comprised two technical replicates. Validation experiments included three independent biological replicates per strain, each measured in technical duplicate.

The BDA MIC_50_ was defined as the lowest tested drug concentration producing at least a 50% reduction in growth relative to the corresponding drug-free control after 48 h. The area under the concentration–response curve (AUC) summarised growth across the tested drug concentration range and was calculated by trapezoidal integration using a linear concentration axis in GraphPad Prism. Supra-MIC growth (SMG) was calculated as the mean of relative growth at all tested concentrations strictly above each strain’s own 48-h MIC_50_, with growth expressed relative to that strain’s drug-free control. Thus, the concentration equal to the MIC_50_ was excluded from the SMG calculation. Relative susceptibility was expressed using strain-to-parent ratios. Resistance ratios based on MIC and AUC were calculated as 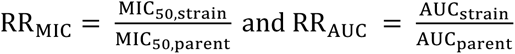. The tolerance ratio based on SMG was calculated as 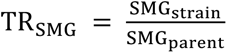. All comparisons used the corresponding parental wt strain under the same drug and assay conditions. SMG and TR_SMG_ could not be calculated when no MIC_50_ was reached within the tested concentration range. Concentration–inhibition curves were additionally fitted to estimate maximal inhibition and Hill coefficients.

Susceptibility was further assessed using ETEST® strips (bioMérieux). Overnight cultures grown in RPMI–MOPS containing 2% glucose at 37 °C were diluted to OD_600_ 0.1 in 1× PBS and spread evenly onto RPMI–MOPS agar plates containing 2% glucose and 2% agar. ETEST strips were placed centrally on the inoculated plates. Plates were scanned after 24 and 48 h, with an additional assessment at 72 h for slow-growing strains. The Etest MIC was read after 24 h at the intersection of the inhibition-ellipse boundary with the strip’s concentration scale. The tolerance phenotype was assessed visually as growth within the inhibition ellipse after 48 h relative to the corresponding parental wt strain. Etest assessments included three independent biological replicates per strain. No numerical threshold for colony number, growth density or occupied area within the ellipse was applied. Etests therefore provided a qualitative assessment of tolerance, complemented by quantitative SMG and TR_SMG_ measurements in BDAs.

### Growth kinetics, stress phenotyping and killing assays

For growth-curve experiments, approximately 1 × 10⁶ cells were inoculated into 200 µl medium in flat-bottom 96-well plates. Plates were incubated in a Multiskan GO reader at 37 °C, with 1 min of pulsed shaking before OD₆₀₀ measurements at 30-min intervals for 48 h. MCF responses were measured across 0.0625–64 µg mL⁻¹ in RPMI–MOPS containing 2% glucose. Additional growth experiments were performed at 41 °C or at pH 4.0 using RPMI buffered with 50 mM MES. Apparent specific growth rates were calculated from blank-corrected OD measurements within the OD₆₀₀ range of 0.02–0.80. For consecutive measurements, the interval-specific rate was calculated as 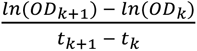 where *ODₖ* is the blank-corrected optical density at measurement time *tₖ* (ℎ), and *k* indexes successive measurements. For each well, the apparent maximum specific growth rate was defined as the median of the five highest interval-specific estimates and expressed in h⁻¹. When fewer than five valid intervals were available, the median of the available interval-specific estimates was used. Wells with fewer than two measurements within the predefined OD range were assigned a rate of zero. Growth rates were derived from the first 24 h.

For the concentration–inhibition curves, the apparent specific growth rate at each MCF concentration was normalised to the matched drug-free control as 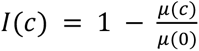, where *c* is the MCF concentration, *μ*(*c*) is the apparent specific growth rate under drug exposure and *μ*(0) is that of the matched control. Mean inhibition at each non-zero concentration was fitted by unweighted nonlinear least squares to a Hill function with a freely fitted maximum: 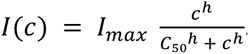. Here, *I_max_* is the fitted maximal fractional inhibition, *C*_50_ is the concentration producing half of *I_max_*, and ℎ is the Hill coefficient describing curve steepness. Concentrations producing 25%, 50% or 75% inhibition relative to the drug-free control (IC₂₅, IC₅₀ and IC₇₅) were reported only when supported by the fitted maximum and measured concentration–response range. *C*_50_ therefore denotes half-maximal inhibition rather than necessarily 50% inhibition of the drug-free growth rate. Plotted points and error bars show means and standard errors, respectively. For population modelling, individual-well growth rates were fitted directly using the four-parameter inhibitory *E_max_* function described under Population-dynamic modelling.

Stress susceptibility was assessed using BDAs with twofold dilution series of H₂O₂ (0.31– 80 mM), calcofluor white (CFW, 0.078–20 mM), Congo red (CR, 0.047–12 µM), EDTA (0.0078–2 mM), nikkomycin Z (NikZ, 0.125–32 µg mL⁻¹), caffeine (0.2–50 mM) or geldanamycin (0.25–64 µg mL⁻¹). NaCl was tested over 0.58–2.5 M using successive dilutions with a factor of 0.85. For osmotic-stress spot assays, overnight cultures were adjusted to OD₆₀₀ = 1, serially diluted tenfold and spotted onto RPMI–MOPS agar supplemented with NaCl or KCl (0.5 or 1 M), or sorbitol (0.5, 1 or 2 M). Plates were imaged after 48 h at 37 °C.

For time–kill experiments, overnight cultures were precultured from OD₆₀₀ = 0.2, and approximately 5 × 10⁶ cells were exposed to 0.0625–64 µg mL⁻¹ MCF in 1 mL medium at 37 °C with shaking at 300 rpm. Samples were collected after 0, 4, 8, 12, 24, 48 and 72 h, serially diluted and plated onto agar containing 1:10000-diluted YPD for microcolony enumeration. Microcolony development was monitored by time-lapse imaging using a Nikon Eclipse Ti2 inverted microscope equipped with a 4× objective and a DS-Qi2 monochrome camera, allowing cells that resumed growth to be distinguished from those that remained non-growing during the observation period. Counts were obtained using a custom Python-based image-analysis script developed and supplied by B.V.d.B. and J.M., corrected for the plated volume and dilution factor, and expressed as CFU mL⁻¹.

Relative viability was calculated as 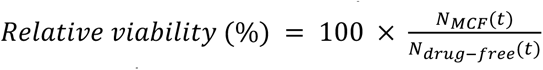 where *N_MCF_*(*t*) and *N_dru_*_g–*free*_(*t*) denote CFU mL⁻¹ for the MCF-exposed culture and the corresponding drug-free control, respectively, for the same strain and sampling time. This measure expresses viable-cell abundance relative to the matched control rather than the fraction of the initial inoculum remaining. Growth-curve experiments comprised three biological replicates per strain, whereas time–kill experiments comprised one biological replicate per strain. Both included two technical repeats per strain– concentration combination performed on separate plates. Technical repeats were not treated as independent biological replicates.

### Whole-genome sequencing and variant identification

Genomic DNA was extracted using the MasterPure Yeast DNA Purification Kit and sequenced by Eurofins Genomics on an Illumina NovaSeq 6000 platform. Read quality was inspected using FastQC and reads were trimmed using Trimmomatic^74^. Variants were identified with perSVade v.1.0.6^75^ against the B8441 reference assembly (s01-m01-r27). Reads were aligned using BWA-MEM, and haploid small variants were called with BCFtools, GATK and FreeBayes. Variants were retained when supported by at least 30-fold coverage, an allele frequency of at least 0.9 and the required caller consensus. Variants present in the wt strain were removed using BCFtools isec. Nuclear variants were annotated using Ensembl Variant Effect Predictor with NCBI translation tables 12 and 3.

### Gene Ontology analysis

Functional enrichment of mutated genes was carried out using ShinyGO v.0.77 with a false-discovery rate (FDR) threshold of 0.05^76^. Proteomics data were analysed in R. Log₂ fold-change (log_2_FC) values were interpreted according to the direction of each comparison, such that positive values indicated higher protein abundance in the first condition named. Proteins were selected for functional visualisation if they exhibited a log₂FC greater than 1 or less than −1 and an adjusted P value below 0.05 in at least one comparison. Gene Ontology (GO) annotations were obtained from the Candida Genome Database (CGD) in Gene Association File (GAF) format (generated 2 April 2026). We combined *C. auris* annotations with annotations transferred from orthologous genes in *C. albicans*. Orthologue relationships were obtained from the CGD orthologue mapping file (generated 14 October 2025). Functional categorisation focused on twelve broad categories. The GO hierarchy was accessed using the R package GO.db^77^. For each predefined biological-process category, the corresponding GO term and its descendant terms were retrieved using the GO biological-process offspring relationships. A protein could be assigned to multiple categories if its annotations matched the descendants of more than one selected GO term.

### CRISPR–CasG genome editing

Targeted gene deletions in *C. auris* were generated using Cas9 ribonucleoprotein (RNP) complexes and the recyclable *SAT1*-flipper selection system. The deletion panel comprised *SLN1*, *NIK1*, *SSK1*, *SSK2*, *PBS2*, *HOG1*, *CEK1* and *MKC1* in the B8441 background. *HOG1* and *MKC1* were additionally deleted in B11222 (clade III) and B11245 (clade IV). The *hog1Δmkc1Δ* double mutant was derived from a marker-recycled *hog1Δ* strain. Because the *FLP* recombinase in the *SAT1*-flipper cassette is under the control of the maltose-inducible *MAL2* promoter, marker excision was induced by growing the *hog1Δ* strain in YP medium containing 2% maltose. Marker loss was assessed by nourseothricin susceptibility and confirmed by PCR before subsequent deletion of *MKC1*. Each strain was assessed in three biological replicates. Guide RNA sequences, construction and verification primers, and repair-cassette details are provided in **Supplementary Table 6**.

Gene-specific deletion constructs were assembled in pSFS2. Homology arms of approximately 500 bp flanking each target gene were PCR-amplified from genomic DNA using StaTaq polymerase and gel-purified. The upstream arm was cloned into pSFS2 digested with XhoI and ApaI, and the downstream arm into the vector digested with NotI and SacII (37 °C for 1 h, followed by heat inactivation at 65 °C for 20 min). Assembly was performed with the NEBuilder HiFi DNA Assembly kit (100 ng vector and 200 ng insert in a final volume of 10 µL; 50 °C for 15 min), including a no-insert control. Reactions were transformed into chemically competent *E. coli* TOP10F′ (CaCl₂ method; heat shock at 42 °C for 45 s) and selected on LB agar containing 100 µg/mL ampicillin. Colonies were screened by colony PCR using primers flanking the insertion site, and plasmids were purified (Macherey-Nagel), screened by restriction analysis and sequence-verified before use.

For transformation of *C. auris*, verified plasmids (6 µg) were linearized with KpnI and SacI and gel-purified. Cells were grown overnight in YPD at 37 °C (220 rpm), diluted to an OD₆₀₀ of approximately 0.4 and grown for about 3 h to an OD₆₀₀ of 1.6–2.2. Cells were then incubated in TE buffer with 100 mM lithium acetate (37 °C, 1 h), followed by addition of 25 mM DTT for a further 30 min, washed with ice-cold water and 1 M sorbitol, and resuspended in 200 µL of 1 M sorbitol. For each transformation, tracrRNA and gene-specific crRNA were annealed (1:1 ratio in duplex buffer; 95 °C for 5 min, then cooled to room temperature), and 3.6 µL of the duplex was combined with 3 µL of 4 µM Cas9 to form the RNP. The RNP was mixed with 500 ng linearized donor DNA and 40 µL of competent cells and electroporated in a 0.2 cm cuvette (1.8 kV, 200 Ω, 25 µF). Cells were recovered in 2 mL YPD at 37 °C for at least 3 h with shaking and plated on YPD containing 200 µg/mL nourseothricin, followed by incubation at 37 °C until colonies appeared. Mutant genotypes were assessed by diagnostic PCR across both cassette junctions and by an internal locus-specific PCR (Q5 polymerase), and positive candidates underwent targeted sequencing.

### Proteome analysis

Wild-type and *hog1Δ* cells were grown for 48 h at 37 °C on RPMI–MOPS agar with or without 2 µg mL−1 CAS. Cells were lysed in 7 M urea, 100 mM ammonium bicarbonate and 5 mM MgCl2 by 12 cycles of bead beating (30 s, speed 6), with cooling between cycles. Lysates were clarified by centrifugation at 16,000g for 2 h and proteins were digested with trypsin. Peptides (200 ng per injection) were analysed using an Orbitrap Astral mass spectrometer coupled to a NeoVanquish liquid chromatography system and a 5.5-cm µPAC column. Peptides were separated over a 7.6-min gradient at 1.25 µl min^−1^ using 0.1% formic acid in water as mobile phase A and 80% acetonitrile containing 0.1% formic acid as mobile phase B. A blank wash injection was included after every 12 injections to assess carryover. Narrow-window data-independent acquisition covered 380-980 m/z. MS1 scans were acquired at a resolution of 240000 with an automatic gain control target of 500% and a maximum injection time of 5 ms. DIA scans used 2-Th isolation windows, a maximum injection time of 3 ms and a resolution of 80000.

Raw data were processed using DIA-NN v.1.8 in library-free mode against the *C. auris* UniProt proteome UP000230249. A two-step match-between-runs workflow was used: an in silico spectral library was generated from the reference proteome, refined using the experimental DIA data and subsequently used to reanalyse the dataset^78^. Protein abundances were log₂-transformed and normalised to the mean protein abundance within each sample. Missing abundance values were set to zero. Differential protein abundance was assessed using protein-wise t-tests, with P values adjusted for multiple testing using the Benjamini–Hochberg procedure. Four pairwise comparisons were evaluated: CAS-treated versus drug-free wt, CAS-treated versus drug-free *hog1Δ*, *hog1Δ* versus wt under CAS exposure, and *hog1Δ* versus wt under drug-free conditions. Positive log₂ fold changes indicated higher abundance in the first condition named in each comparison. Proteins were considered differentially abundant when |log₂ fold change| > 1 and the Benjamini–Hochberg-adjusted P value was < 0.05. Proteins meeting both criteria in at least one comparison were eligible for functional visualisation and were grouped by biological process.

### Cell-wall sample preparation for flow cytometry and TEM

Wt and *hog1Δ* cells were grown overnight in RPMI–MOPS (pH 7, 2% glucose), diluted to OD_600_ 0.1 in PBS, and 50 µL of suspension was spread onto RPMI–MOPS agar (pH 7, 2% glucose, 2% agar) with or without 2 µg mL^−1^ CAS. Plates were incubated for 48 h at 37 °C, and cells were collected by scraping into deionised water and resuspended by gentle aspiration. For fluorescence analysis, cells were pelleted (4,500 g, 1 min), fixed for 2 h at room temperature in 500 µL of 50 mM thimerosal (Sigma-Aldrich), pelleted again (4,000 g, 1 min), washed twice with deionised water, resuspended in 200 µL deionised water and stored at 4 °C until staining.

### Fluorescent staining and flow cytometry

On the day of analysis, 2.5 × 10^6^ fixed cells were transferred to a 96-well round-bottom plate and pelleted (4,500 g, 3 min). Accessible β-glucan was labelled with 1.5 µg mL^−1^ Fc– hDectin-1a (InvivoGen) in 100 µL for 45 min at room temperature in the dark. Cells were pelleted (4,500 g, 5 min) and resuspended in 100 µL staining solution containing Alexa Fluor 488-conjugated anti-human IgG (1:200, Invitrogen H10120) to detect bound Dectin-1, concanavalin A–Texas Red (25 µg mL^−1^, Invitrogen C825) to detect mannan, and calcofluor white (CFW, 25 µg mL^−1^, Sigma-Aldrich) to detect chitin. After 30 min at room temperature in the dark, cells were pelleted (4,500 g, 5 min), washed once with PBS, resuspended in 50 µL PBS and analysed immediately on an Attune NxT flow cytometer (Thermo Fisher). Fluorescence was recorded for 30000 single cells per sample using an identical gating strategy for all samples: debris was excluded by side-scatter versus forward-scatter area, and singlets were selected by forward-scatter height versus area. Data were analysed in Floreada.io, and mean fluorescence intensities were compared by two-way ANOVA with Šídák’s multiple-comparisons test (drug-free versus CAS within each genotype) in GraphPad Prism v10.6.0.

### Transmission electron microscopy

Cells were grown on agar with or without CAS and scraped as described above, without chemical fixation. Aliquots (5 µL) of concentrated cells were loaded into 100 µm flat specimen carriers and cryofixed using a Leica EM PACT2 high-pressure freezer^79^ with rapid transport system (2100 bar). Frozen carriers were freeze-substituted in acetone containing 1% (w/v) osmium tetroxide in a Leica EM AFS2 at −90 °C, gradually warmed to room temperature, infiltrated with Spurr’s resin through a graded acetone–resin series and polymerised. Ultrathin sections (70–90 nm) were cut with a diamond knife (Diatome) on a Leica UC6 ultramicrotome, collected on copper grids and post-stained with uranyl acetate and lead citrate. Sections were imaged at 80 kV on a JEOL 1400 Plus microscope equipped with an AMT UltraVUE camera. Microscopy was conducted in the University of Aberdeen’s Microscopy and Histology Core Facility. Cell-wall thickness was measured in ImageJ and compared in GraphPad Prism.

### Cell-wall monosaccharide composition with HPIC

Cell-wall composition was determined for cells grown for 48 h at 37 °C on RPMI–MOPS agar with or without 2 µg mL^−1^ CAS, as described above, and scraped into PBS. Cells were disrupted with glass beads in a FastPrep instrument (12 cycles of 30 s at speed 6, with 2 min on ice between cycles). The insoluble cell-wall fraction was collected by centrifugation (4,000 g, 5 min), washed five times with 1 mL ice-cold 1 M NaCl and five times with 20 mL ice-cold 1 M NaCl, resuspended in 5 mL deionised water and pelleted again (11,000 g, 5 min). Proteins were extracted by resuspending the pellet in 1 mL SDS extraction buffer (50 mM Tris-HCl pH 7.5, 2% w/v SDS, 1 mM EDTA, 150 mM NaCl, 0.3 M β-mercaptoethanol) and heating at 100 °C for 10 min. Walls were washed four times with deionised water to remove residual detergent, frozen in liquid nitrogen and freeze-dried. Dried walls were hydrolysed in 2 M trifluoroacetic acid (TFA, Sigma-Aldrich) for 3 h at 100 °C to release glucose, mannose and glucosamine, used as measures of glucan, mannan and chitin, respectively. Hydrolysates (0.4 µL) were analysed by high-performance anion-exchange chromatography with pulsed amperometric detection (HPAEC-PAD, a form of HPIC) on a Dionex carbohydrate analyser equipped with a CarboPac PA20 column (0.4 × 150 mm), guard column and ED50 detector. Monosaccharides were eluted with a 5-100 mM KOH gradient at 8 µL min^−1^ for 25 min. Each condition was measured in three biological replicates.

### Solid-state NMR spectroscopy

Wt and *hog1*Δ cells were grown for 48 h at 37 °C on RPMI–MOPS (pH 7.0) containing ¹³C-labelled glucose and ^15^N ammonium chloride, with or without 2 µg mL^−1^ CAS. Cells were harvested, fixed with PFA and washed with H_2_O. All spectra were acquired on a Bruker 18.8 T (800 MHz ¹H Larmor frequency) spectrometer using 3.2 mm H/F-X or HCN MAS probes at 12 kHz MAS and a probe temperature of 273 K. ¹³C chemical shifts were referenced. Rigid polysaccharides were detected by ¹³C cross-polarisation (CP), mobile components by quantitative ¹³C direct polarisation (DP) with a 30 s recycle delay, and highly mobile components by refocused J-INEPT. The 1D ^1^H-^13^C CP and INEPT spectra were recorded with 64 scans and a 3 s recycle delay, and DP spectra with 64 scans. Carbohydrate resonances were assigned from 2D ¹³C–¹³C correlation spectra (**Supplementary Fig. 8**), analysed in CcpNMR 3. Relative abundances of β-1,3-glucan, β-1,6-glucan, chitin and mannan within each mobility fraction were estimated from the volumes of resolved cross-peaks and expressed as the relative distribution within that fraction. These values describe the composition within each fraction and not absolute polymer abundance.

The water accessibility of rigid polysaccharides was assessed using ¹³C-detected ¹H T2-filter experiments (128 scans, 3 s recycle delay). A 2 × 3.0 ms ¹H T2 filter suppressed up to 95% of the polysaccharide magnetization while retaining the water signal. Water magnetization was then transferred to the polysaccharides during a ¹H mixing period of 0.02–100 ms, followed by ¹H–¹³C CP and ¹³C detection. Relative signal intensity versus the control spectrum was used to describe hydration. Spectra were processed in Bruker TopSpin 4.0.0. The ssNMR acquisition and processing parameters for 2D spectra are listed in **Supplementary Table 7**.

### Dynamic MCF exposure and *in vitro* competition

Competition experiments under dynamic MCF exposure were performed in custom continuous-flow bioreactors, using a nourseothricin (NTC)-resistant marked wild-type strain and either the evolved Nik1^E472K^ isolate or the resistant Fks1^S639Y^ isolate. The marked strain and competition framework were based on previously described methods^37^. Marked wild type and competitor were mixed at 1:1 or 99:1 and adjusted to a total final OD_600_ of 0.04 in 50 mL RPMI 1640 containing 0.2% glucose, 0.165 M MOPS (pH 7.0) and 100 µg mL^−1^ ampicillin. Cultures were maintained at 37 °C with stirring at 240 rpm, and each condition was run in duplicate. To mimic drug clearance, drug-free medium was delivered by peristaltic pump at 14.13 mL min^−1^ in 20 bursts of 13.2 s per day (≈3.1 mL per burst). Each burst displaced an equal volume of drug-containing culture through a passive siphon overflow, which kept the working volume constant at 50 mL. Repeated replacement of drug-containing medium with drug-free medium progressively diluted the MCF concentration over time, approximating mono-exponential decay (calculated half-life ≈ 14 h). MCF (800 µg, 16 µg mL^−1^ final) was added at 0, 24 and 48 h, immediately after sampling. Samples (2 mL) were collected at 0, 3, 8, 24, 27, 36, 48, 51, 56 and 72 h, serially diluted in PBS and plated on YPD, then replica-plated onto YPD containing 100 µg mL^−1^ NTC. Frequencies of the marked wt were calculated as the proportion of NTC-resistant colonies among total colonies, and the competitor frequency as the remainder.

### Murine systemic infection and *in vivo* competition

Eight-week-old female BALB/c mice (Janvier Labs) were housed three per cage with food and water ad libitum. Mice were randomly allocated to treatment groups of six animals per strain and treatment condition. Immunosuppression was induced by intraperitoneal cyclophosphamide (200 mg kg^−1^, in a 200 µL volume) 3 d before infection and this was repeated on day 1 after infection (150 mg kg^−1^, in a 200 µL volume). Mice were infected through the lateral tail vein with 1 × 10^7^ cells in 100 µL PBS. MCF treatment (5 mg kg^−1^, intraperitoneal) started 4 h after infection and was repeated every 24 h for seven doses. Control animals received PBS. Animals were humanely killed on day 7. Liver, spleen, kidneys, brain and heart were weighed and homogenized in 1 mL PBS using two 30-s FastPrep cycles. Homogenates were serially diluted, plated on YPD agar and incubated for 48 h at 37 °C. Colonies were enumerated using the automated image-analysis software AutoCellSeg^80^, which removes observer bias from the counts. Organ burden was expressed as CFU per gram of tissue. All animal procedures were approved by the KU Leuven animal ethics committee (projects 079/2020 and 148/2022).

### Clinical and interspecies HOG pathway variation

Protein-altering variation in *SLN1*, *NIK1*, *SSK1*, *SSK2*, *PBS2*, *HOG1*, *CEK1* and *MKC1* was assessed using the global *C. auris* genomic dataset of Gifford *et al.*^39^. The dataset comprised 13,193 publicly available Illumina paired-end whole-genome sequencing datasets from the NCBI Sequence Read Archive and seven newly sequenced isolates. Reads were aligned to the Clade I B8441 v3 nuclear reference genome (GCA_002759435.3), and 12,644 genomes with >10 Mb aligned nuclear sequence were retained. We used the variant calls and sample classifications generated by Gifford *et al.*^39^ directly; reads had been mapped using BWA v.0.7.17 within Snippy v.4.6.0, with variants called using FreeBayes-parallel v.1.3.6 and normalised to the B8441 reference. No independent remapping or variant recalling was performed. The clade-level dataset contained 12,367 epidemiological and 275 laboratory-derived genomes, of which only epidemiological samples were retained. For each variant, carrier frequency was calculated within the corresponding clade as the number of genomes carrying the variant divided by the number assessed (*Number/Total* × 100). Because all genomes were mapped to the Clade I B8441 reference, variant–clade combinations present in ≥95% of epidemiological genomes of a clade were removed as clade-associated background variation. For the epidemiological lollipop plots (**Fig. 4c**), missense, stop-gained and explicitly annotated frameshift variants were retained, whereas complex/multiresidue and in-frame insertion/deletion variants were retained only in **Supplementary Table 5**. This yielded 110 distinct protein-altering variants: 81 missense, 13 stop-gained and 16 frameshift variants.

### Protein variant-effect prediction

Variant effects were predicted using ESM1b, an evolutionary-scale protein language model^27,28^. For missense substitutions, ESM scores were calculated as the difference between the log likelihood of the mutant and reference amino acid, with more-negative scores indicating lower sequence compatibility. For each reference protein, all possible single-amino-acid substitutions were scored and a two-component Gaussian mixture model was fitted to the resulting distribution. The intersection between the components defined the protein-specific high-impact threshold (**Supplementary Fig. 4**). Observed missense variants below this threshold were classified as predicted high impact. For stop-gained variants, the lowest ESM score among substitutions preceding the introduced stop position was used and evaluated against the corresponding protein-specific threshold. Frameshifts were not ESM-scored because the multiresidue approach was not validated for frameshifts. Instead, they were classified as putatively high impact when located within or upstream of an annotated functional domain, whereas terminal frameshifts downstream of the final annotated domain were not. Complex/multiresidue and in-frame insertion/deletion variants were excluded from the predicted loss-of-function (LoF) analysis. Putative LoF variants therefore comprised high-impact missense and stop-gained variants together with qualifying frameshifts.

### Experimental evolution of *FKS1*-mediated resistance

Susceptible wt and tolerant Nik1^E472K^ populations (both with an MIC_50_ of 0.5 µg mL^−1^) were evolved for 7 days in MCF in 200 µL RPMI–MOPS containing 2% glucose and 1% DMSO, with a 1:10 transfer into fresh medium every 24 h. Four wells per plate were drug-free controls, leaving 92 drug-exposed populations per strain, each well being treated as an independent population. Three exposure regimens were applied, each with four replicate populations per starting concentration. Under static exposure, the MCF concentration was kept constant at 0.125, 0.25, 0.5, 1 or 2 µg mL^−1^ (20 populations). Under stepwise-doubling exposure, the concentration was doubled every 24, 48 or 72 h from a starting concentration of 0.125, 0.5 or 2 µg mL^−1^ (36 populations). Under stepwise-halving exposure, the concentration was halved every 24, 48 or 72 h from a starting concentration of 2, 8 or 32 µg mL^−1^ (36 populations). This resulted in 23 regimen– concentration combinations (5 static and 18 stepwise).

### Deep sequencing and variant analysis

Genomic DNA was extracted from endpoint populations using the MasterPure Yeast DNA Purification Kit (Biosearch Technologies/LGC) according to the manufacturer’s instructions. The *FKS1* HS1/HS3 and HS2 regions were amplified using KAPA polymerase and the region-specific primers listed in **Supplementary Table 6**. Amplicons were purified using SPRI beads, dual-indexed in a second PCR and purified again before sequencing. Libraries were sequenced at the Plateforme d’Analyses Génomiques (IBIS, Université Laval, Québec, Canada) on an AVITI instrument (Element Biosciences) using a paired-end 300-base sequencing kit. Reads were demultiplexed by sample indexes before downstream analysis.

Endpoint populations were analysed by paired-end deep sequencing of two *FKS1* amplicons covering HS1/HS3 and HS2, respectively. Reads from wt and Nik1^E472K^-background populations were processed using gyōza v1.2.0 within a Snakemake workflow configured for random-variant detection. Annotated haplotype read counts and allele frequencies were collated using custom Python scripts. Each well was analysed as a separate population, with frequencies evaluated within the corresponding amplicon. Sequence haplotypes were retained when supported by ≥10,000 reads and present at a frequency of ≥0.1%. Allele frequencies were expressed as *f* = 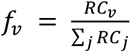 where *f*ᵥ is the frequency of sequence haplotype *v*, *RC*ᵥ is its read count, and *j* indexes the sequence haplotypes contributing to the frequency calculation within the same population and amplicon. The denominator is the sum of their read counts. Frequencies are fractions and are multiplied by 100 when expressed as percentages. Downstream analysis focused on amino acid changes within HS1 (residues 635–645), HS2 (1351– 1357) and HS3 (686–696). Hotspot variants reaching ≥5% frequency within a population were scored as emerged. A population-level emergence score was calculated by summing the frequencies of qualifying variants across the three hotspots. Variant incidence was defined as the number of populations containing at least one qualifying variant divided by the total number of analysed drug-exposed populations within the corresponding exposure regimen, pooling starting concentrations. Incidence was compared between wt and Nik1^E472K^ backgrounds using two-sided Fisher’s exact tests, and emergence scores were compared using two-sided Mann–Whitney U tests.

### Population-dynamic modelling

We used a stochastic population model to examine how pre-existing echinocandin tolerance could influence resistance emergence during MCF exposure. The model links experimentally measured growth and viable-cell responses to changing drug concentrations, population turnover, mutation and immune-mediated clearance. Repeated simulations represent chance variation in cell division, loss and resistance emergence. Model settings and strain-specific inputs are summarised in **Supplementary Table 8**.

Separate simulations were initiated with wt, *hog1*Δ or Nik1^E472K^ as the sole parental background. Wt and tolerant trajectories were overlaid from separate simulations. The Fks1^S639Y^ phenotype represented resistant descendants in all backgrounds. Populations were represented as absolute viable-cell counts in a single, well-mixed 5 L blood-equivalent compartment.

Experimental growth response inputs

OD_600_ growth curves described how rapidly each strain could grow at each MCF concentration. After blank subtraction, interval-specific growth rates (µ_j_) were calculated within the corrected OD_600_ range of 0.02-0.80 as

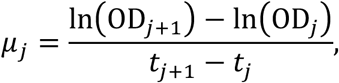

where *j* indexes successive measurements. For each well, the apparent maximum specific growth rate was the median of the five highest interval-specific estimates. Wells without measurements in the predefined OD_600_ range were assigned a rate of zero. These estimates have units of h^−1^ and describe apparent growth, not a separately measured killing rate.

Individual-well growth rates were fitted against MCF concentration using a four-parameter inhibitory Emax function:

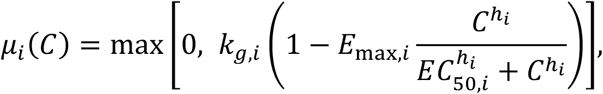

where *C* is the MCF concentration, *k*_g,*i*_ is the drug-free apparent growth rate, *E*_max,*i*_ is the maximum fractional inhibition, *EC*_50,*i*_ is the concentration producing half-maximal inhibition and the Hill coefficient ℎ*_i_* describes the steepness of the response for genotype *i*. Each genotype, including Nik1^E472K^, retained its own fitted growth parameters. Means and standard errors at each concentration were used for visualisation, and fitting used individual-well data. Dose-specific lag times were derived from the same growth curves. When growth onset was not observed within 24 h, lag was treated as right-censored and 24 h was used as the model boundary.

CFU-derived killing and recovery

Viable-cell counts supplied information about early loss and recovery that OD-derived growth rates do not resolve. For each genotype and concentration, two technical CFU replicates were summarised on the log_10_ scale. Measurements outside the countable range were excluded. The early change and subsequent recovery were calculated as

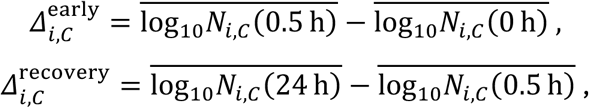

where *N_i_*_,*C*_(*t*) denotes viable cell abundance for genotype *i* at concentration *C* and time *t*. *C* = 0 denotes the drug-free control. Changes were separated into an initial 0-0.5 h decline and a 0.5-24 h recovery phase, accounting for changes also present in the matched drug-free control. Effects were interpolated between positive concentrations on a logarithmic concentration scale, connected linearly to the no-drug condition and held at the highest measured effect above the experimental concentration range. A strain-specific CFU profile was not available among the model inputs for Nik1^E472K^, so the *hog1*Δ CFU profile supplied its early-loss and recovery terms, including drug-free control changes. This proxy was restricted to the CFU-derived layer. The substitution is a modelling assumption, not evidence that the strains have identical viable-cell dynamics.

CFU-derived dynamics governed the first 24 h. At later daily MCF peaks, only the additional early decline relative to the matched drug-free control was repeated. The same CFU proxy therefore also supplied the later killing pulses for Nik1^E472K^. The genotype-specific log-killing pulse was defined as

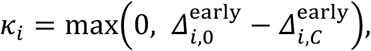

and was applied at each new MCF peak as an instantaneous survival probability

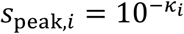

The first dose was already represented in the empirical first-day CFU trajectory, so no additional initial killing pulse was applied.

Population dynamics and resistance emergence

Each simulation tracked one parental population, *P*, and its derived resistant population, *R*. Initial conditions were *P*(0) = 2 × 10^5^ cells and *R*(0) = 0, equivalent to 40 CFU mL^-1^. Thus, each run started with 100% of the selected parent and no resistant cells. Parent and resistant descendants shared a carrying capacity of *K* = 2 × 10^8^ viable cells. Outside the CFU-calibrated first day, division depended on each population’s fitted growth response to the current MCF concentration. Crowding increased cell loss while allowing division to continue, so a stable total population could still undergo turnover and generate mutations. The total division flux and shared crowding-loss hazard were

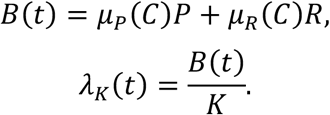

Outside the CFU-calibrated first day and between the instantaneous killing pulses, the deterministic expectation of the stochastic system can be written as

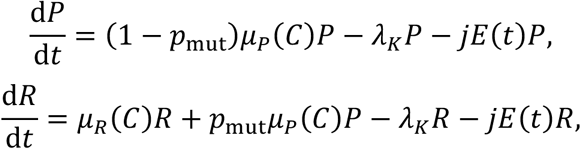

where *p*_mut_ is the mutation probability per parental division opportunity and *jE*(*t*) is the immune-mediated per-capita removal rate. The parental equation accounts for growth, transfer of mutant daughters to the resistant population, crowding and immune clearance. The resistant equation additionally receives those mutant daughters.

Individual simulations sampled births and losses as random events using a tau-leaping procedure, with a maximum time step of 0.1 h. For a step of duration *τ*, births were sampled from

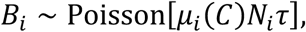

and crowding-associated losses were sampled as

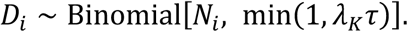

Here, the index *i* denotes the parental or resistant population, *N* is its current viable-cell count, and *B* and *D* denote births and crowding losses during the step. Mutation opportunities were based on the expected number of parental divisions rather than on net population increase:

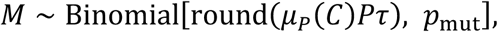

with *p*_mut_ = 5 × 10^−10^ per division opportunity and *M* the number of mutant daughters during a simulation step. Mutant daughters were reassigned from the parental birth compartment to the resistant population, preserving total births. Mutations could therefore occur during turnover even when net population growth was small.

MCF exposure

MCF concentration was set to a peak of 16 µg mL^−1^ every 24 h and declined between doses with a 15 h half-life:

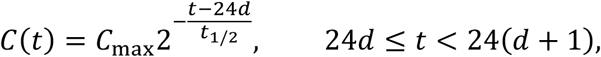

where *C*_max_ = 16 µg mL^-1^, *t*_1/2_ = 15 h and *d* is the dosing-day index. Each new peak was the total post-dose concentration and residual drug was not added on top of that peak. No serial dilution or population transfer was applied during the 14-day treatment simulation.

Immune-mediated clearance

A separate immune-effector compartment represented a common clearance pressure on all fungal genotypes. Effector recruitment was independent of fungal population density, and effectors were also lost by decay. The density, *E*, followed

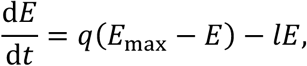

where *E*_max_ = 10^6^ cells mL^-1^ is the reservoir parameter, *q* = 10^−3^ h^-1^ is the recruitment coefficient and *l* = 10^−3^ h^-1^ is the decay coefficient. Initial effector density was *E*(0) = 10^2^ cells mL^-1^. The common interaction coefficient was *j* = 1.2 × 10^−6^ mL effector-cell^- 1^ h^-1^. Over one simulation step, the corresponding fungal survival probability was

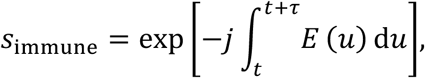

and fungal cells were sampled binomially. Immune parameters were adopted from the previous modelling frameworks^37,73^ and were not fitted to the present MCF data. This compartment represents a shared clearance assumption rather than a strain-specific measurement of host immunity.

### Simulations and outcome classification

For each parental background, 500 simulations were performed. A master random seed generated separate random streams. Outcomes were evaluated at days 7 and 14. At each endpoint, parental survival meant that at least one parental cell remained. Resistance was classified as observed if any resistant cell had appeared by that time, surviving if at least one resistant cell remained, and dominant if resistant cells outnumbered parental cells. Dominance did not require complete replacement. Wilson 95% confidence intervals are given for the summarising percentages. Population composition was calculated within simulations with surviving cells and then summarised across batches. These summaries describe stochastic variation under fixed model inputs, not uncertainty in biological parameters or clinical event probabilities. Population counts were converted to blood-equivalent CFU mL^−1^ by division by 5000 mL.

### Statistical analysis and software

Statistical tests, biological replicate numbers and definitions of independent experiments are specified in the corresponding Methods sections and figure legends. Unless stated otherwise, tests were two-sided. Corrections for multiple comparisons are described for the relevant analyses.

Statistical analyses and concentration–response AUC calculations were performed using GraphPad Prism. Growth-curve analysis and concentration–response fitting were performed using custom Python scripts, with NumPy for numerical operations, pandas for data handling, SciPy for nonlinear curve fitting and Matplotlib for visualisation. The stochastic population model was implemented in Python using custom scripts, with NumPy for numerical calculations and random sampling and Matplotlib for plotting simulated population trajectories. Additional custom Python scripts were used for sequencing-data processing and microcolony enumeration.

Whole-genome sequencing analyses used FastQC, Trimmomatic and perSVade v1.0.6, including BWA-MEM, BCFtools, GATK and FreeBayes, with variant annotation using Ensembl Variant Effect Predictor. Amplicon-sequencing data were processed using gyōza v1.2.0 within a Snakemake workflow. Gene Ontology enrichment was performed using ShinyGO v0.77.

Proteomics data were processed using DIA-NN v1.8, with downstream analyses performed in R, including functional categorisation using GO.db. Flow-cytometry data were analysed using Floreada and GraphPad Prism, cell-wall thickness was quantified using ImageJ, and tissue colony counts were obtained using AutoCellSeg. ssNMR data were processed using Bruker TopSpin v4.0.0 and analysed using CcpNMR v3.

## Acknowledgments

D.S. C S.J. were supported by PhD fellowships from the Fund for Scientific Research Flanders – FWO (11PRR24N C 11J8122N respectively). H.C. was supported by a postdoctoral fellowship granted by KU Leuven Internal Funds (PDMT2/23/032) and a postdoctoral fellowship from the European Molecular Biology Organization – EMBO (ALTF 1105-2024). B.V.d.B. was supported by an FWO postdoctoral fellowship (12O1922N).

A.M. was supported by funding from the European Research Council (ERC) under the European Union’s Horizon 2020 research and innovation programme ERC Synergy “FungalTolerance” (grant No. ERC-SyG-2020 951475) awarded to J.B. and M.R. Mass spectrometry instrumentation was funded by the Berlin University Alliance under grant (515_BUA_Proteomics). We thank Michael Mülleder and the Charité/BUA proteomics core facility (Charité Berlin) for hands-on support in conducting the proteomic experiments.

A.L. and A.Zv. were funded by the Latvian Council of Science, project No. lzp-2023/1-0312, “Characterisation of fungal cell wall architecture using solid-state nuclear magnetic resonance spectroscopy at fast spinning regime.”

R.A.F. is part of the Medical Research Council Centre for Medical Mycology MR/N006364/2 and is supported by a Wellcome Trust Career Development Award (225303/Z/22/Z). R.A.F. and H.G. also received support from the MRC Doctoral Training Grants (MR/P501955/2, MR/W502649/1) and the NIHR Exeter Biomedical Research Centre. The views expressed are those of the authors and not necessarily those of the NIHR or the Department of Health and Social Care.

N.A.R.G. acknowledges the support of Wellcome Trust Investigator, Collaborative, Equipment, Strategic and Biomedical Resource awards (200208, 215599, 224323). They also thank the MRC (MR/M026663/2), MR/Y002164/1 and the MRC Centre for Medical Mycology (MR/N006364/2) for support. This study/research is funded by the National Institute for Health and Care Research (NIHR) Exeter Biomedical Research Centre (BRC). The views expressed are those of the author(s) and not necessarily those of the NIHR or the Department of Health and Social Care. For the purpose of open access, the author has applied a ‘Creative Commons Attribution (CC BY) licence’ to any Author Accepted Manuscript version arising from this submission.

The group of T.G. acknowledges support from the Spanish Ministry of Science and Innovation, grant number PID2025-176398OB-I00 funded by MICIU/AEI /10.13039/501100011033 and by FEDER, UE and grant number PLEC2023-010225 funded by MICIU/AEI /10.13039/501100011033, as well as support from “La Caixa” foundation (grant numbers LCF/PR/HR21/00737, CI23-20260, LCF/PR/HR21/52410006 and LCF/PR/GN18/50310010); Instituto de Salud Carlos III (CIBERINFEC CB21/13/00061- ISCIII-SGEFI/ERDF and DTS25/00141); European Commission, Horizon Europe-HORIZON-MSCA-2023-DN-01-01 (grant number 101168618).

C.R.L. was supported by a Genome Canada and Genome Québec grant (no. 6569) and holds the Canada Research Chair in Cellular Systems and Synthetic Biology.

The group of P.V.D. was supported by FWO under the framework of the JPIAMR – Joint Programming Initiative on Antimicrobial Resistance fund (project CycleDrug) and by a C3 grant from the Industrial Research Fund of KU Leuven (C3/22/007).

We thank Terry Roemer and colleagues for providing the *C. albicans* GRACE strains used in this study, and Karl Kuchler and colleagues for sharing the *N. glabratus* deletion collection.

## Author contributions

Conceptualization: H.C. Methodology: D.S., S.J., A.Zi., A.M., A.Zv., A.P., G.B., A.K.D., L.W., B.V.d.B. C H.C. Investigation: D.S., S.J., A.Zi., A.M., A.Zv., A.P., G.B., H.G., V.B., A.K.D., L.W., D.M., I.L., Z.W., C.L.R. C H.C. Formal analysis: D.S., S.J., A.Zi., A.M., A.Zv., A.P., H.G., V.B., I.L., D.M-J, N.A.G C H.C. Resources: J.M., R.A.F., N.A.R.G., T.G., A.L., M.R., C.A.M., C.R.L., P.V.D. Data curation: D.S., S.J. C H.C. Writing – original draft: D.S., S.J. C H.C. Writing – review and editing: D.S., S.J., A.Zi., A.M., A.Zv., A.P., G.B., H.G., V.B., A.K.D., I.L., D.M-J, L.W., B.V.d.B., C.L.R., P.v.d.B., J.M., R.A.F., J.B., N.A.G., T.G., M.A.L., M.R., C.A.M., C.R.L., P.V.D. C H.C. Visualization: D.S., S.J., A.M., A.Zv., A.P. C H.C. Supervision: P.v.d.B., J.M., R.A.F., J.B., N.A.G., T.G., M.A.L., M.R., C.A.M., C.R.L., P.V.D. C H.C. Funding acquisition: J.B., T.G., C.R.L., P.V.D. C H.C.

## Data and materials availability

Accession numbers for raw sequencing data, repository links for code and strain availability information will be provided upon formal publication.

