## supplementary for "HOG pathway modulation is a determinant of echinocandin treatment failure in *Candidozyma auris*"

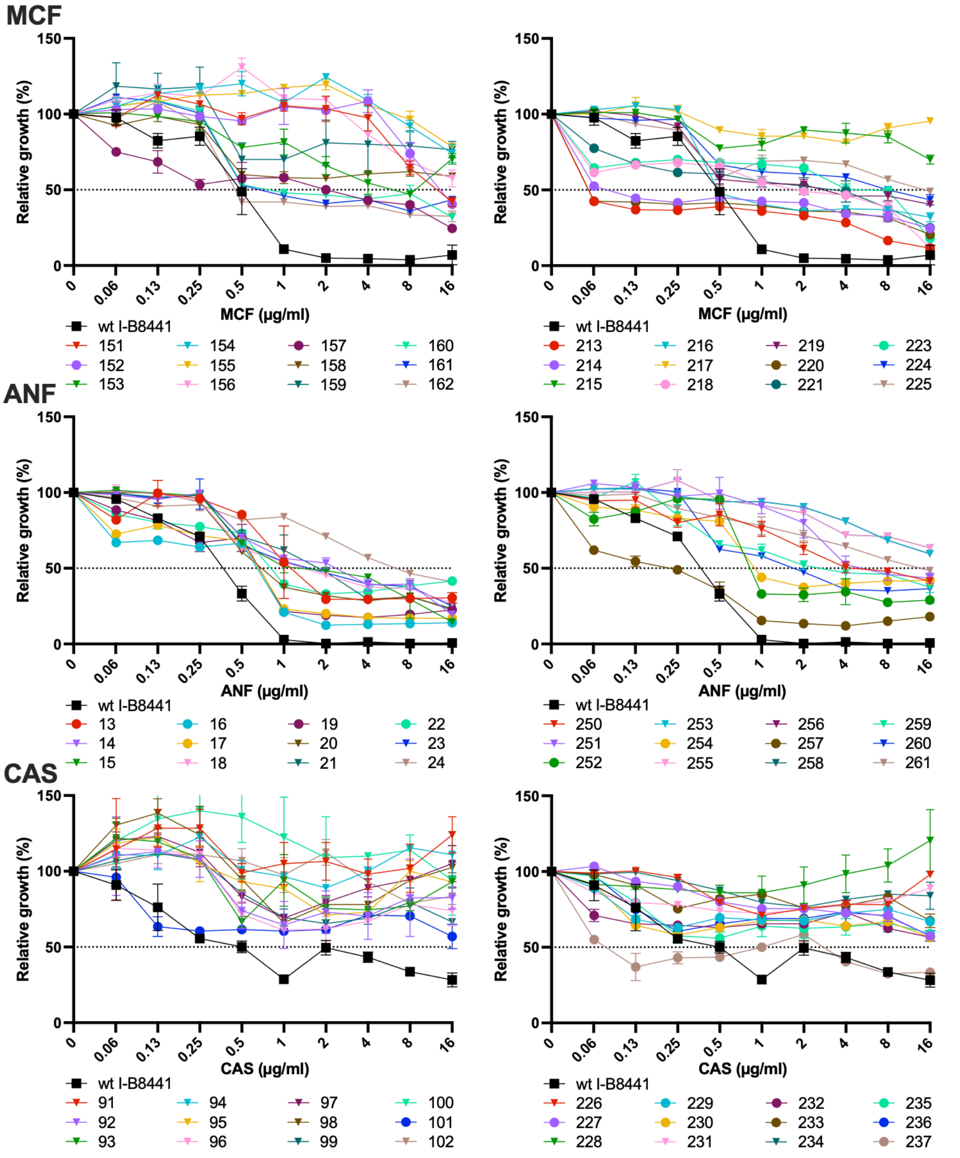


**MCF**

| | 151 | **152** | 153 | 154 | 155 | 156 | **157** | 158 | 159 | 160 | 161 | 162 | **213** | **214** | 215 | 216 | 217 | **218** | 219 | **220** | **221** | **223** | 224 | 225 |
|---|---|---|---|---|---|---|---|---|---|---|---|---|---|---|---|---|---|---|---|---|---|---|---|---|
| $RR_{MIC}$ | 32.0 | 32.0 | 16.0 | 64.0 | 64.0 | 64.0 | 8.0 | 64.0 | 64.0 | 2.0 | 2.0 | 1.0 | 0.1 | 0.3 | 64.0 | 1.0 | 64.0 | 4.0 | 8.0 | 0.1 | 8.0 | 16.0 | 32.0 | 32.0 |
| $RR_{AUC}$ | 8.8 | 9.4 | 7.3 | 11.7 | 12.0 | 9.5 | 4.8 | 7.5 | 9.7 | 5.5 | 5.2 | 4.6 | 2.6 | 4.0 | 10.1 | 4.6 | 11.0 | 4.4 | 5.8 | 3.8 | 4.9 | 5.5 | 6.5 | 7.3 |
| $TR_{SMG}$ | | | 11.4 | | | | 5.2 | | | 6.9 | 6.6 | 6.0 | 4.8 | 6.0 | | 5.9 | | 5.2 | 7.0 | 5.8 | 5.1 | 2.9 | | |
| GR µ | 0.8 | 0.9 | 0.9 | 0.9 | 0.8 | 0.7 | 0.9 | 0.8 | 0.9 | 0.7 | 0.8 | 0.8 | 0.9 | 0.9 | 1.0 | 0.8 | 0.8 | 0.9 | 0.9 | 0.9 | 0.9 | 1.0 | 0.9 | 0.9 |
| GR AUC | 1.1 | 1.0 | 1.0 | 1.1 | 1.1 | 0.8 | 0.8 | 0.9 | 0.9 | 0.9 | 1.0 | 0.9 | 0.9 | 0.9 | 1.0 | 1.0 | 1.0 | 0.9 | 1.3 | 0.9 | 0.9 | 0.9 | 1.2 | 1.1 |

**ANF**

| | **13** | 14 | 15 | **16** | **17** | 18 | **19** | 20 | 21 | **22** | 23 | 24 | 250 | 251 | **252** | 253 | **254** | 255 | 256 | **257** | 258 | 259 | 260 | 261 |
|---|---|---|---|---|---|---|---|---|---|---|---|---|---|---|---|---|---|---|---|---|---|---|---|---|
| $RR_{MIC}$ | 4.0 | 8.0 | 4.0 | 2.0 | 2.0 | 4.0 | 2.0 | 2.0 | 4.0 | 2.0 | 4.0 | 16.0 | 16.0 | 16.0 | 2.0 | 64.0 | 2.0 | 64.0 | 64.0 | 0.5 | 64.0 | 8.0 | 4.0 | 32.0 |
| $RR_{AUC}$ | 10.2 | 11.5 | 10.0 | 4.9 | 6.1 | 11.4 | 6.7 | 9.4 | 10.4 | 11.9 | 11.6 | 15.9 | 15.4 | 16.3 | 9.8 | 21.9 | 12.9 | 21.9 | 21.9 | 4.9 | 21.9 | 14.1 | 12.0 | 17.9 |
| $TR_{SMG}$ | 28.3 | 27.6 | 27.4 | 12.5 | 16.9 | 31.7 | 18.4 | 27.0 | 26.5 | 34.6 | 32.2 | 38.4 | 39.0 | 41.1 | 28.9 | | 37.9 | | | 17.2 | | 39.5 | 34.0 | |
| GR µ | 0.9 | 0.7 | 0.7 | 0.9 | 0.9 | 0.7 | 1.0 | 0.6 | 0.7 | 1.0 | 0.8 | 0.8 | 1.0 | 1.1 | 0.9 | 1.0 | 1.0 | 1.0 | 0.9 | 0.9 | 0.9 | 0.9 | 0.9 | 0.9 |
| GR AUC | 0.9 | 0.9 | 0.9 | 0.9 | 0.9 | 0.8 | 0.9 | 0.7 | 0.8 | 1.0 | 1.1 | 0.8 | 1.3 | 1.4 | 0.8 | 1.3 | 1.0 | 1.3 | 1.1 | 1.0 | 1.2 | 1.2 | 1.2 | 1.1 |

**CAS**

| | 91 | 92 | 93 | 94 | 95 | 96 | 97 | 98 | 99 | 100 | **101** | 102 | 226 | **227** | 228 | **229** | **230** | 231 | **232** | **233** | 234 | **235** | **236** | **237** |
|---|---|---|---|---|---|---|---|---|---|---|---|---|---|---|---|---|---|---|---|---|---|---|---|---|
| $RR_{MIC}$ | 32.0 | 32.0 | 32.0 | 32.0 | 32.0 | 32.0 | 32.0 | 32.0 | 32.0 | 32.0 | 32.0 | 32.0 | 32.0 | 32.0 | 32.0 | 32.0 | 32.0 | 32.0 | 32.0 | 32.0 | 32.0 | 32.0 | 32.0 | 0.1 |
| $RR_{AUC}$ | 3.0 | 2.2 | 2.2 | 2.9 | 2.5 | 2.0 | 2.6 | 2.5 | 2.0 | 3.0 | 1.8 | 2.4 | 2.3 | 1.9 | 2.9 | 2.0 | 1.7 | 2.2 | 1.7 | 2.1 | 2.3 | 1.7 | 1.8 | 1.0 |
| GR µ | 0.8 | 0.8 | 0.8 | 0.8 | 0.9 | 0.8 | 1.0 | 0.7 | 0.8 | 0.6 | 0.9 | 0.8 | 0.9 | 0.8 | 0.9 | 0.9 | 0.9 | 0.9 | 0.9 | 0.8 | 0.9 | 0.9 | 0.9 | 0.9 |
| GR AUC | 1.0 | 1.0 | 0.9 | 1.0 | 1.2 | 1.1 | 1.0 | 0.9 | 0.8 | 0.7 | 0.9 | 0.8 | 1.2 | 0.9 | 1.2 | 1.0 | 0.9 | 1.2 | 1.0 | 0.8 | 1.0 | 1.0 | 0.9 | 1.0 |

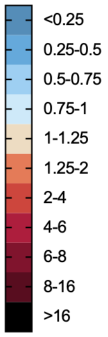

**Supplementary Fig. 1: Echinocandin susceptibility and drug-free growth of all 72 screened evolved isolates.** Broth dilution assays (BDAs) for the 72 evolved isolates, comprising 24 isolates evolved in micafungin (MCF), anidulafungin (ANF) or caspofungin (CAS), evaluated after 48 h of incubation against their respective selection drug. For each drug, isolates are displayed in two groups of 12, with the parental wt strain B8441 included in each graph as a control (black squares). Growth is expressed relative to the corresponding drug-free control. Curves show the mean ± s.e.m. of two technical replicates. Dotted horizontal lines indicate 50% growth. Isolates selected for whole-genome sequencing (WGS) are depicted with circular symbols.

Heat maps show relative resistance ratios by MIC ($RR_{MIC}$) and area under the concentration–response curve ($RR_{AUC}$), and tolerance ratios by supra-MIC growth ($TR_{SMG}$), together with drug-free growth metrics (growth rate, μ, and area under the growth curve, AUC). Relative ratios were calculated as $XR_y = \frac{y_{\text{evolved}}}{y_{\text{parental}}}$, where $X$ denotes the phenotype (resistance or tolerance) and $y$ denotes the corresponding metric. Drug-free growth metrics were normalized to the wt in the same way and determined under control growth conditions containing 2% glucose over 48 h. Isolate identifiers in bold indicate strains selected for WGS. Crossed cells indicate that $TR_{SMG}$ could not be calculated.

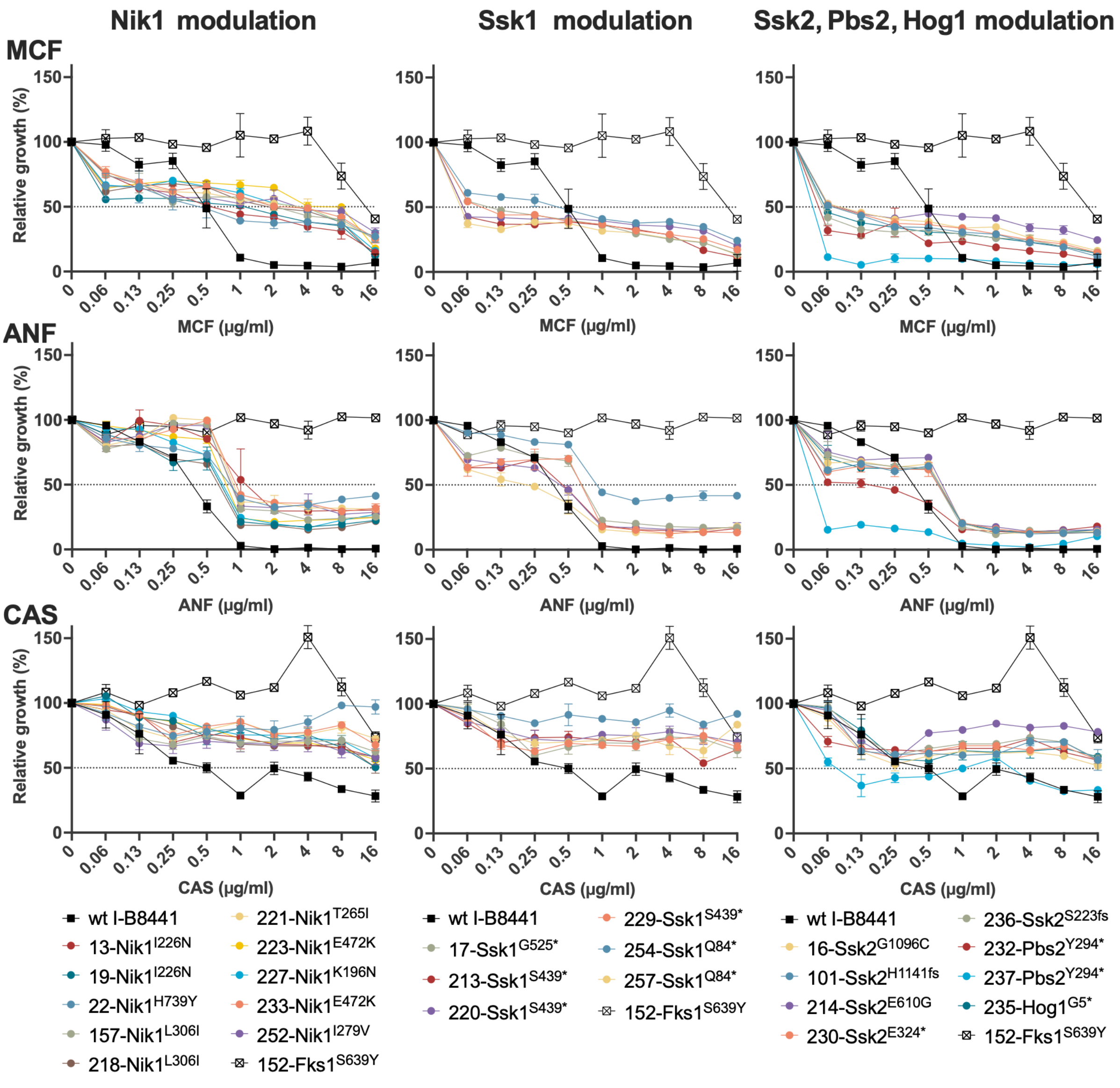

**Supplementary Fig. 2: Echinocandin concentration–response profiles of the sequenced evolved isolates.** Broth dilution assays (BDAs) for the 25 evolved isolates selected for whole-genome sequencing, evaluated against micafungin (MCF), anidulafungin (ANF) and caspofungin (CAS) after 48 h of incubation. Isolates are grouped by variants in Nik1; Ssk1; or Ssk2, Pbs2 and Hog1 (columns), with each group evaluated against all three echinocandins (rows). The parental wt strain B8441 (filled black squares) and resistant isolate 152 carrying the Fks1$^{S639Y}$ substitution (crossed open squares) are included in every graph as susceptible and resistant controls, respectively. Growth is expressed relative to the corresponding drug-free control. Dotted horizontal lines indicate 50% growth. Strain labels indicate the isolate number and defining protein variant. Curves show the mean ± s.e.m.

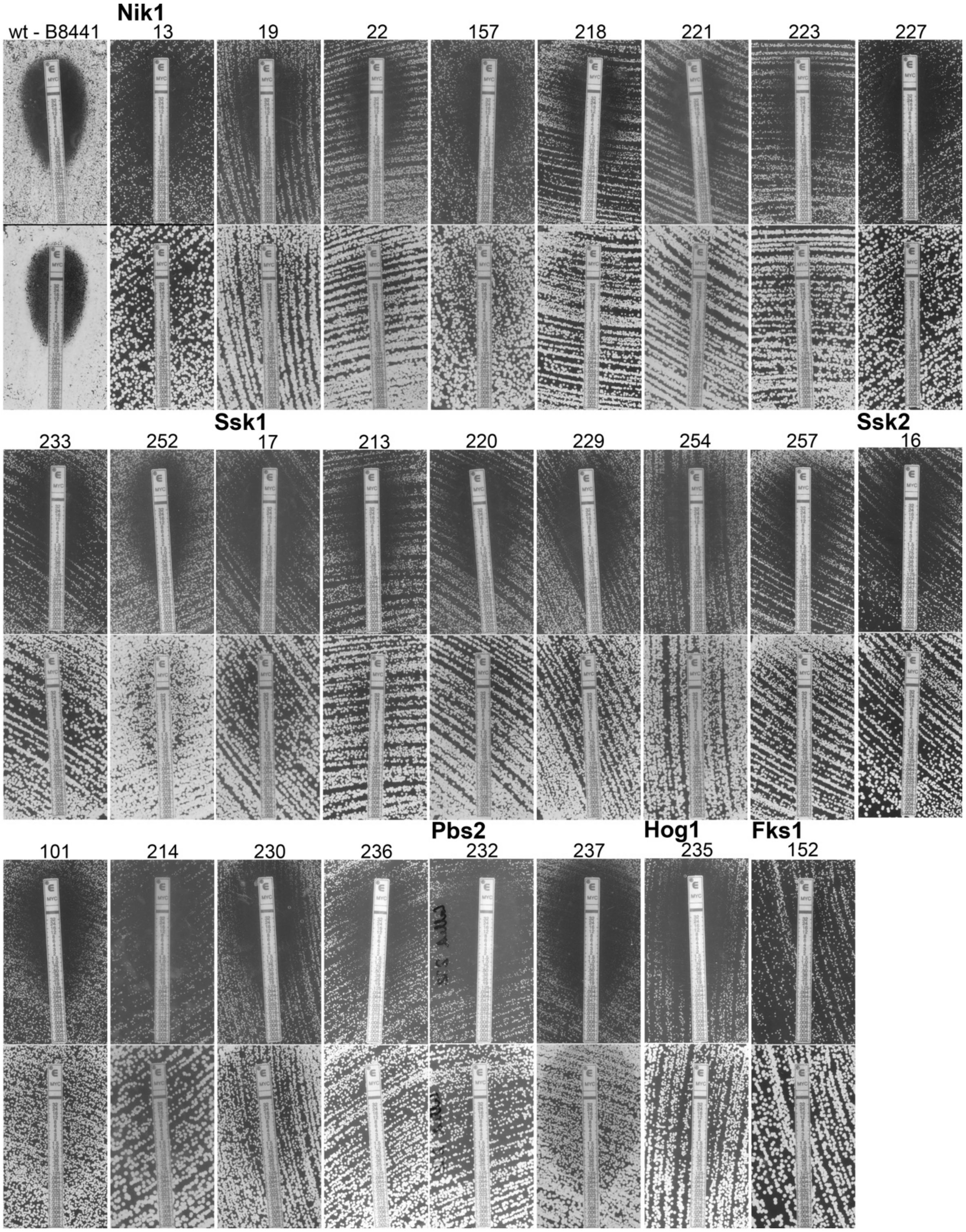


**Supplementary Fig. 3: Micafungin Etest profiles of all sequenced evolved isolates.** Micafungin (MCF) Etests at 24 h (top) and 48 h (bottom) for the parental wild-type (wt) strain B8441 and all 25 evolved isolates selected for whole-genome sequencing. Isolates are grouped by the mutated protein (Nik1, Ssk1, Ssk2, Pbs2, Hog1 or Fks1), with isolate numbers indicated above each pair of images. Increased growth within the inhibition ellipse after prolonged incubation (48 h) illustrates the tolerance phenotype of the HOG-pathway adapted strains. Isolate 152, carrying the resistance-conferring Fks1$^{S639Y}$ substitution, serves as the resistance comparator.

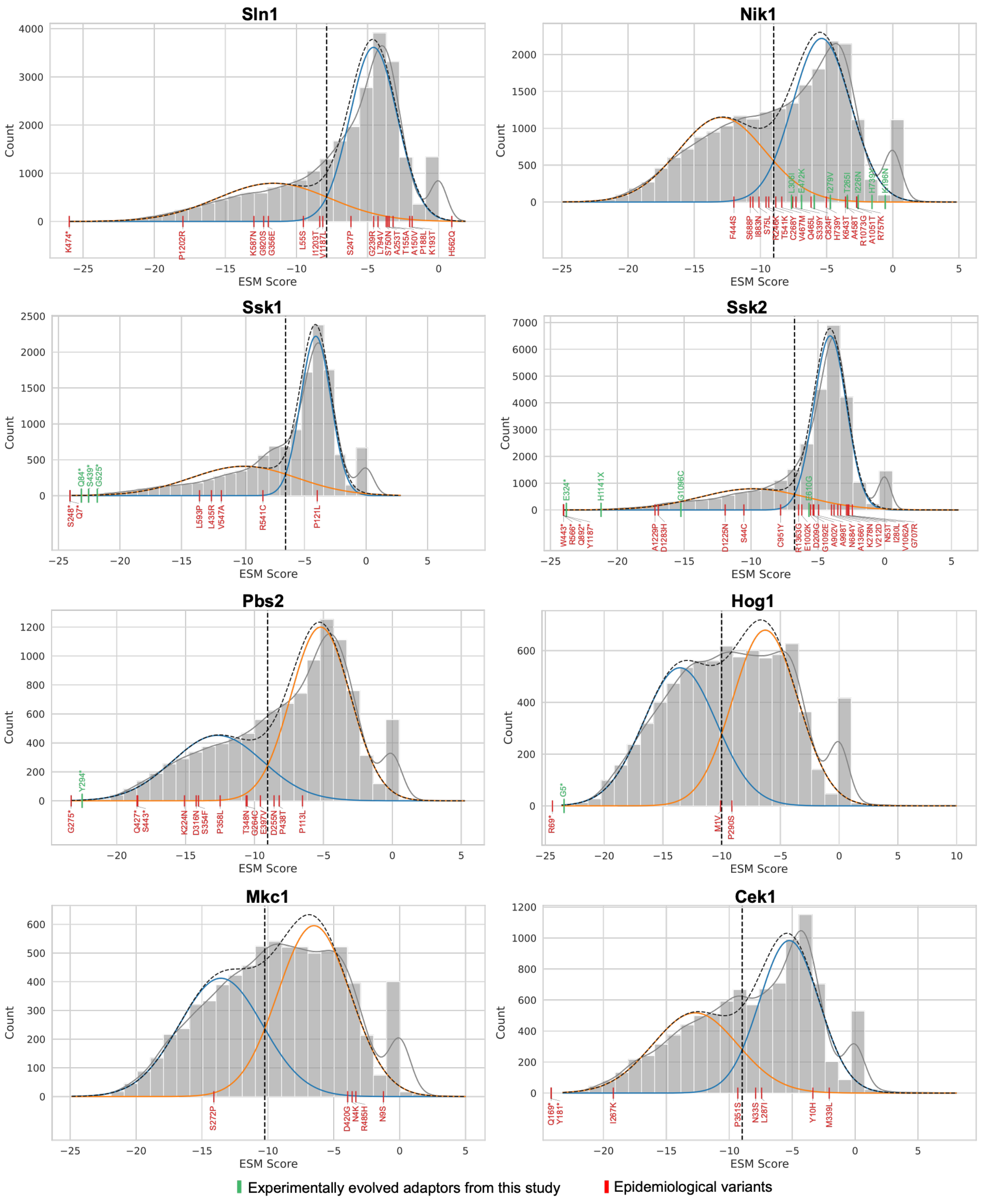


**Supplementary Fig. 4: ESM-score distributions and predicted variant impact for HOG pathway proteins.** Grey histograms show the distributions of evolutionary scale modelling (ESM) scores obtained from FungAMR[1] for Sln1, Nik1, Ssk1, Ssk2, Pbs2, Hog1, Mkc1 and Cek1. A two-component Gaussian mixture model was fitted to each protein-specific distribution, excluding synonymous variants. Blue and orange curves indicate the individual Gaussian components, and black dashed curves indicate the combined fits. Vertical black dashed lines mark the intersection of the two components, defining the protein-specific threshold below which variants were classified as having a predicted high impact. More negative ESM scores indicate lower compatibility with the reference protein sequence and a greater predicted likelihood of functional disruption. Experimentally evolved variants identified in this study (green)

and epidemiological variants (red) are indicated by labelled vertical markers at their corresponding scores.

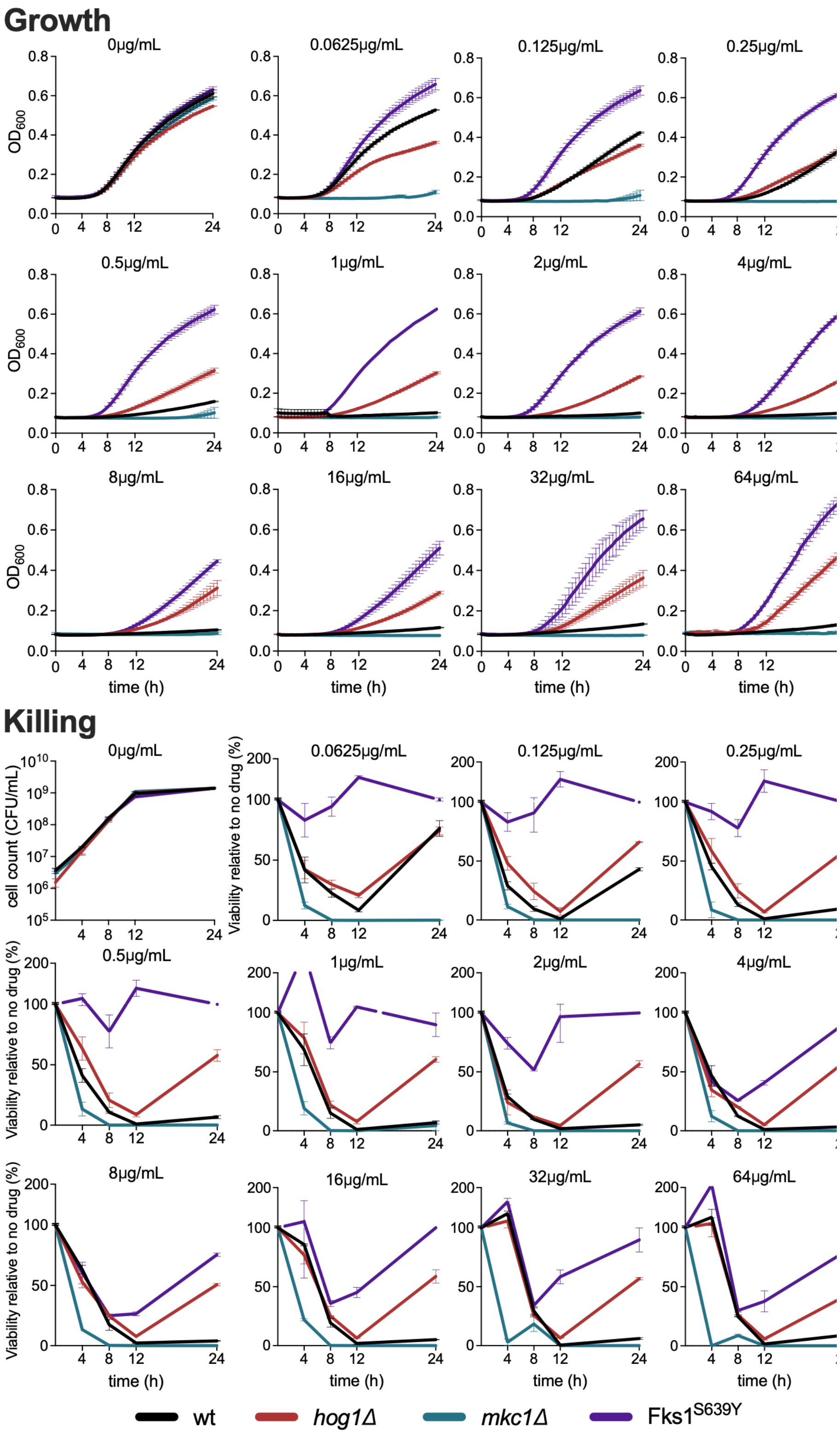


**Supplementary Fig. 5: Growth and viable-cell dynamics across the complete micafungin concentration gradient.** Optical-density growth trajectories (upper panels) and CFU-based viability measurements (lower panels) for *C. auris* wt, tolerant *hog1*Δ, hypersusceptible *mkc1*Δ and resistant Fks1$^{S639Y}$ during 24 h of exposure to micafungin (MCF, 0.0625–64 μg ml$^{-1}$), alongside drug-free controls. Growth was monitored by optical density at 600 nm ($OD_{600}$). Viable-cell counts are expressed as CFU $ml^{-1}$ in drug-free medium and as percentages relative to the corresponding drug-free control at the same time point during MCF exposure.

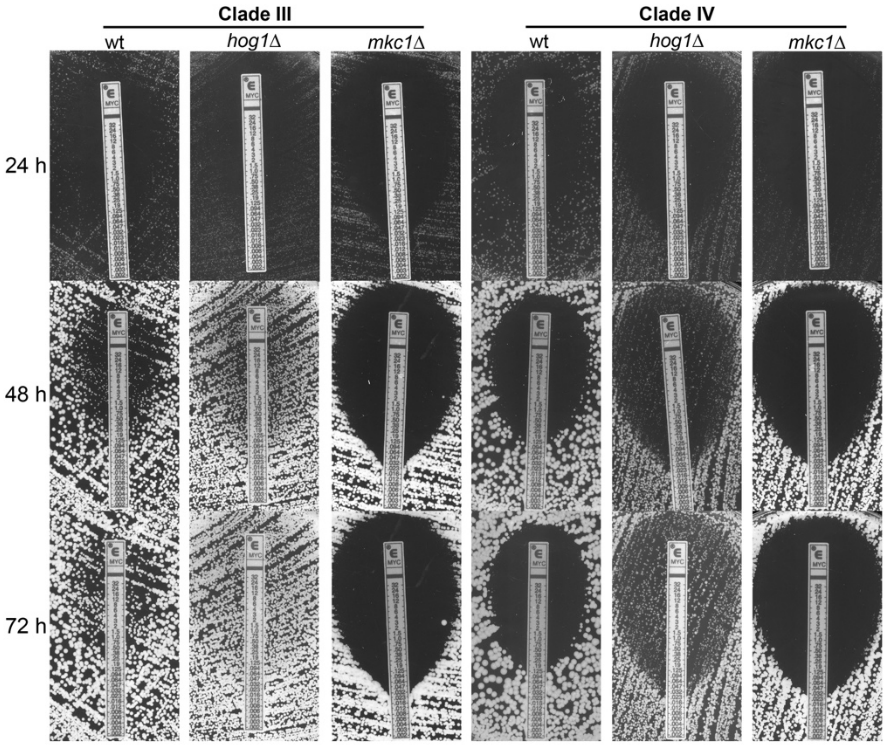


**Supplementary Fig. 6: Micafungin Etest profiles of *HOG1* and *MKC1* deletion strains in other clades.** MCF Etests for *C. auris* wt, *hog1*Δ and *mkc1*Δ strains in clade III (left) and clade IV (right) genetic backgrounds. Images show the inhibition profiles after 24 h, 48 h and 72 h, allowing comparison of the inhibition ellipse and growth within the ellipse during prolonged incubation. The corresponding parental wt strain is included for each clade.

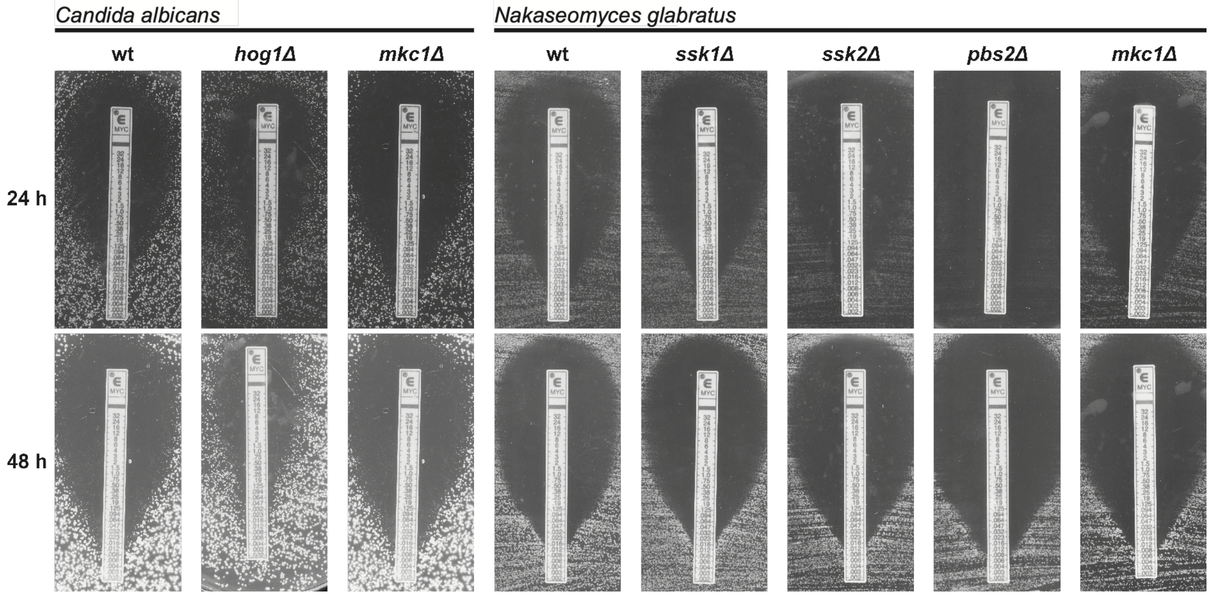

**Supplementary Fig. 7: Micafungin Etest profiles of HOG- and cell-wall-integrity-pathway deletion strains in *C. albicans* and *N. glabratus*.** MCF Etests for *C. albicans* wt, *hog1*Δ and *mkc1*Δ strains, and *N. glabratus* wt, *ssk1*Δ, *ssk2*Δ, *pbs2*Δ and *mkc1*Δ strains (right). Images show the inhibition profiles after 24 h and 48 h, with the corresponding parental wt controls included for each species.

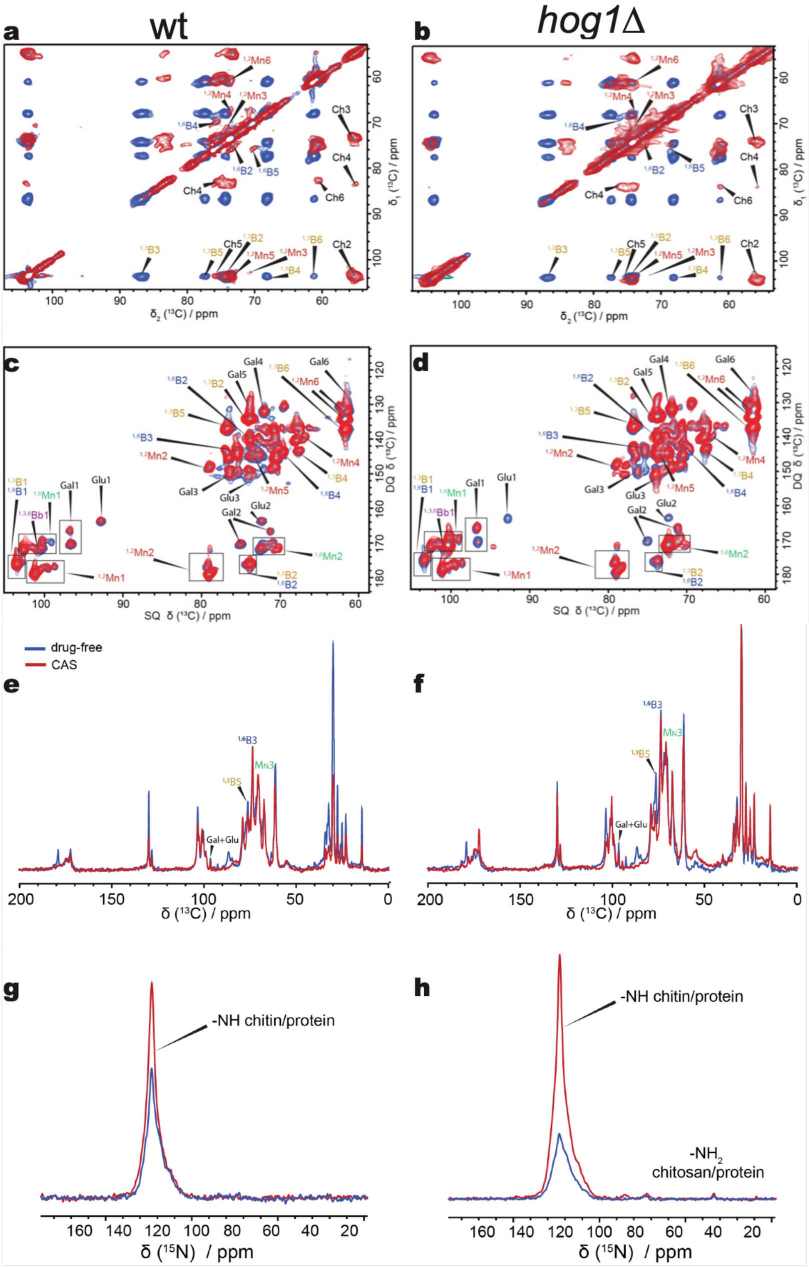


**Supplementary Fig. 8: Solid-state NMR spectra supporting compositional analysis of intact *C. auris* cells.** Spectra compare wt (left column) and *hog1Δ* (right column) cells grown under drug-free conditions (blue) or with 2 μg $mL^{-1}$ CAS (red). **a,b)** Two-dimensional $^{13}C$–$^{13}C$ dipolar-assisted rotational resonance (DARR) spectra used to identify comparatively rigid carbohydrate components. **c,d)** Two-dimensional direct-polarisation INADEQUATE (DP-INADEQUATE) spectra used to identify comparatively mobile carbohydrate components. Labels indicate carbohydrate assignments and carbon positions: B, β-glucan; Mn, mannan; Ch, chitin; Gal, galactose; and Glu, glucose, with glycosidic linkages indicated where applicable. **e,f)** One-dimensional $^{13}C$ direct-polarisation spectra acquired with a 30-s recycle delay and used for carbohydrate quantification. **g,h)** One-dimensional $^{15}N$ spectra acquired using $^{1}H$–$^{15}N$ cross-polarisation (CP). Annotated signals correspond to amide NH groups of chitin/proteins and amino $NH_2$ groups of chitosan/proteins. Overlapping blue and red contours indicate shared chemical-shift positions between treatment conditions; contour intensities in the two-dimensional spectra do not directly quantify absolute carbohydrate abundance. Chemical shifts are expressed in parts per million (ppm).

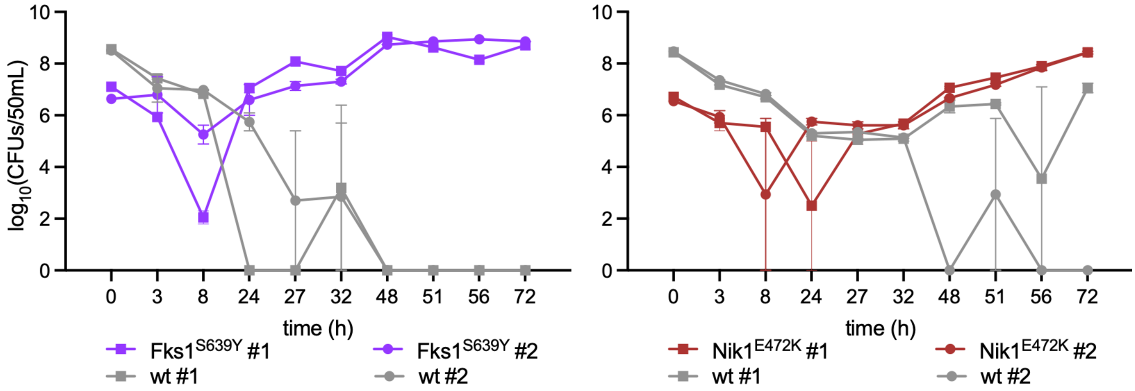


**Supplementary Fig. 9: Absolute population sizes during bioreactor competition under dynamic micafungin exposure.** Viable-cell numbers of wt competing with the resistant Fks1$^{S639Y}$ isolate or tolerant Nik1$^{E472K}$ isolate over 72 h. Competitions were initiated with 99% wt and 1% competitor. Population sizes are expressed as $\log_{10}$ CFU per 50-mL bioreactor, representing the total viable-cell number of each strain throughout the working volume. Labels #1 and #2 denote separate bioreactor runs, distinguished by square and circular symbols. Each time point within each run was assessed using two technical plating replicates, with points showing their mean.

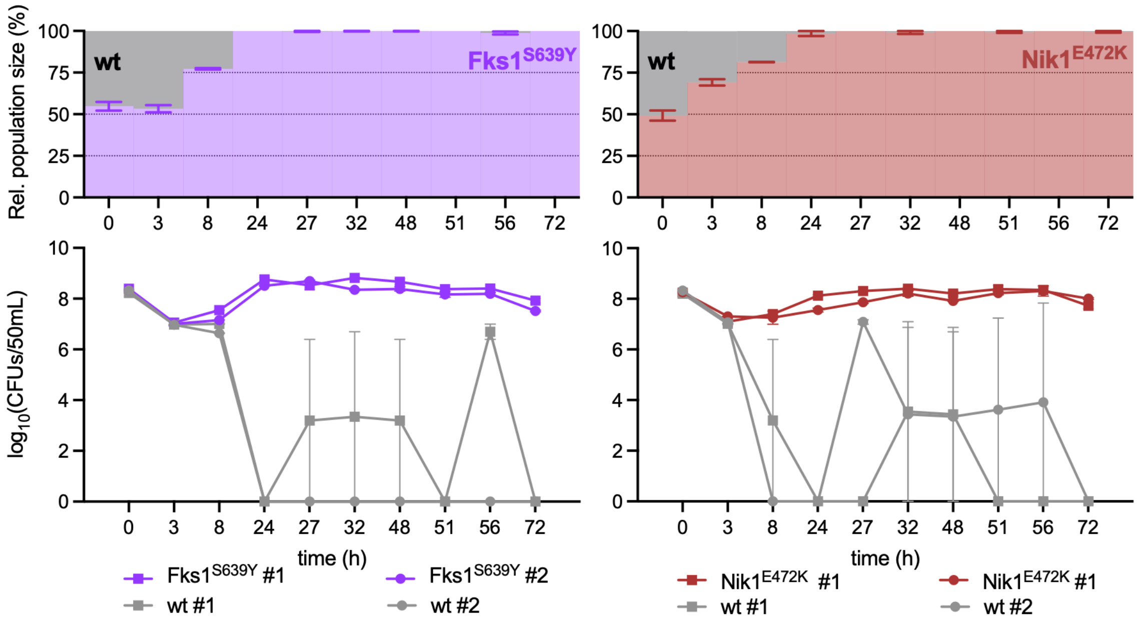


**Supplementary Fig. 10: Relative and absolute population sizes during bioreactor competition initiated at equal strain frequencies.** Wt competed with the resistant Fks1$^{S639Y}$ isolate or tolerant Nik1$^{E472K}$ isolate over 72 h under dynamic micafungin exposure. Competitions were initiated with 50% wt and 50% competitor. Upper panels show stacked plots with relative abundance of wt and the resistant or tolerant competitor, expressed as percentages of the combined population. Strain proportions were determined by differential plating using the nourseothricin-resistance marker in the wt background. Lower panels show absolute population sizes, expressed as $\log_{10}$ CFU per 50-mL bioreactor, representing the total viable-cell number of each strain throughout the working volume. Labels #1 and #2 denote separate bioreactor runs, distinguished by square and circular symbols, respectively. Each time point within each run was assessed using two technical plating replicates, with lower-panel points showing their mean.

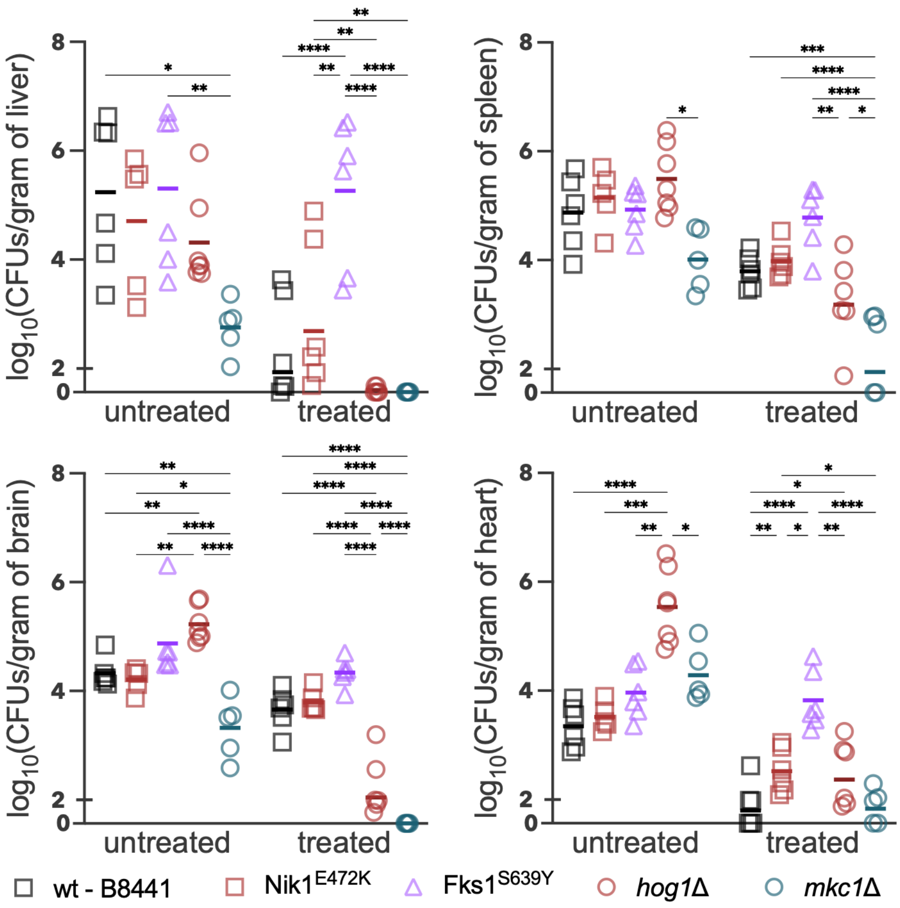


**Supplementary Fig. 11: Fungal burden in additional organs following micafungin treatment.** Liver, spleen, brain and heart fungal burdens in immunosuppressed BALB/c mice infected intravenously with susceptible wt (B8441), tolerant Nik1$^{E472K}$ or *hog1*Δ, resistant Fks1$^{S639Y}$, or hypersusceptible *mkc1*Δ strains. Mice received daily intraperitoneal injections of MCF (5 mg kg$^{-1}$) or phosphate-buffered saline (untreated controls), and organs were collected on day 7 for CFU enumeration. Fungal burdens are expressed as $\log_{10}$ CFU per gram of tissue. Symbols represent individual animals, and horizontal lines indicate means. Statistical significance was assessed separately for each organ by two-way ANOVA of $\log_{10}$-transformed fungal burdens, with genotype and treatment as factors, followed by Šídák's multiple-comparisons test. Lines indicate the comparisons shown. *$P < 0.05$, **$P < 0.01$, ***$P < 0.001$ and ****$P < 0.0001$.

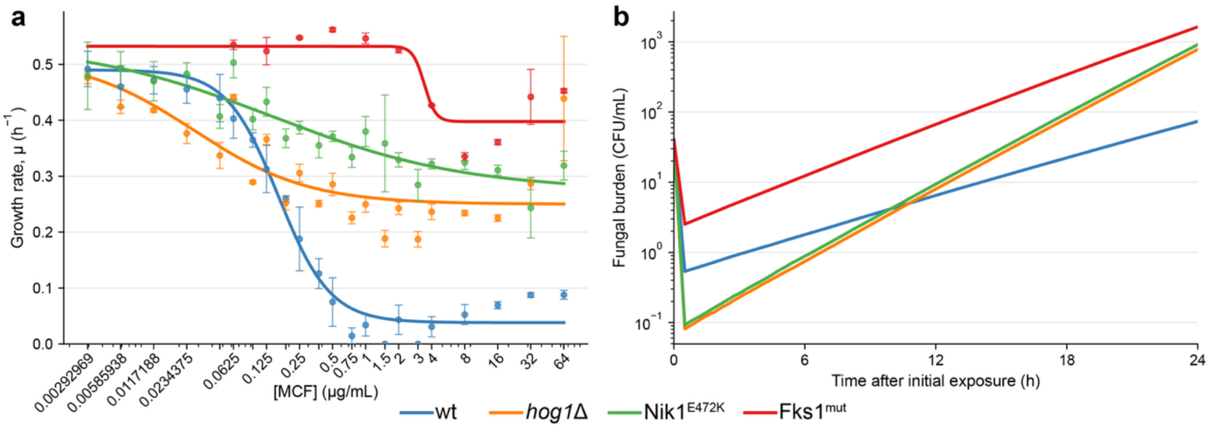


**Supplementary Fig. 12: Experimental growth-response inputs and simulated first-day population dynamics. a)** Original μ data and fitted growth response. Growth rates were derived from $OD_{600}$ measurements recorded over 24 h in RPMI containing 2% glucose and the indicated MCF concentrations. Points represent means of three repeats, error bars indicate s.e.m., and solid lines show the fitted concentration–growth relationships used as model inputs. **b)** Simulated first-day population dynamics. Lines show separate single-strain simulations of the initial decline and subsequent recovery in viable-cell numbers. These first-day dynamics were parameterised using absolute CFU counts from the killing experiments in **Fig. 2b** and **Supplementary Fig. 5**, comprising two technical repeats, each plated in triplicate. CFU-derived inputs describe the initial 0–0.5-h decline and the subsequent 0.5–24-h recovery. The *hog1Δ* CFU profile was used for both tolerant strains, but Nik1$^{E472K}$ retained its own OD-derived growth response in panel **a**. In the treatment simulations, CFU-derived dynamics (**b**) governed the first 24 h, whereas the concentration-dependent growth relationships (**a**) governed growth thereafter, alongside the other model processes described in the Methods and **Supplementary Table 8**. Fks1$^{mut}$ denotes the resistant phenotype represented by Fks1$^{S639Y}$.

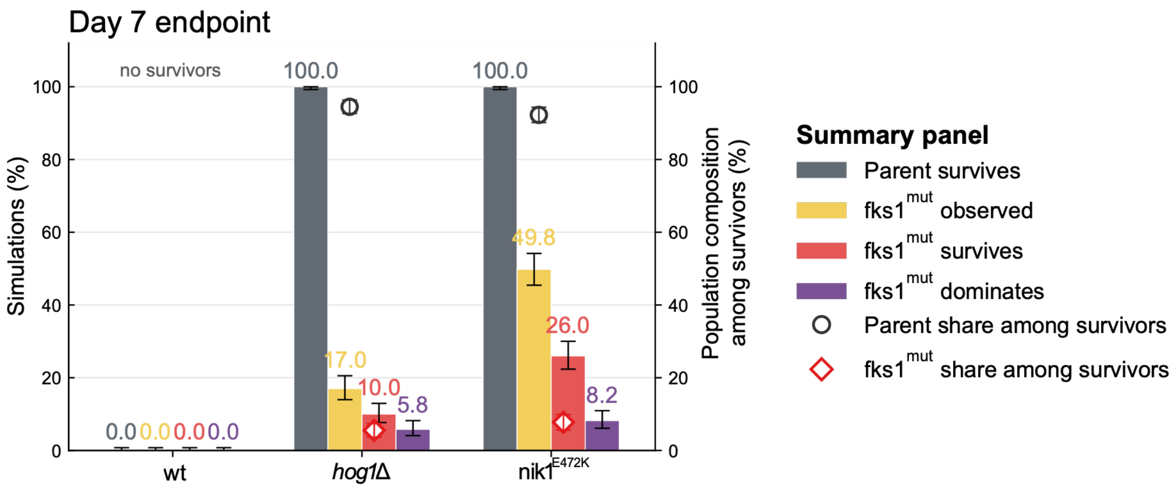


**Supplementary Fig. 13: Day-7 outcomes of stochastic population simulations under MCF exposure.** Day-7 outcomes from the same ensemble of 500 independently seeded simulations per wt, *hog1Δ* or Nik1$^{E472K}$ parental background used in **Fig. 5d,e**. Bars indicate the percentages of simulations in which the parental population survived (grey), an Fks1$^{mut}$ lineage had been observed by day 7, or an Fks1$^{mut}$ lineage survived or was dominant at day 7. Numbers above bars indicate percentages, and error bars show binomial Wilson 95% confidence intervals. Open circles and diamonds indicate mean parental and Fks1$^{mut}$ population shares, respectively, among simulations with survivors, plotted against the right y-axis. Their error bars show 95% confidence intervals for the mean.

**Supplementary Table 1: Strain inventory.** Species, clade per species, isolate designation, relevant genotype, parental background and source are listed for parental strains, engineered mutants, the nourseothricin-resistant wt and the 25 sequenced experimentally evolved isolates. For evolved isolates, only the defining HOG-pathway or *FKS1* genotype is shown. Susceptibility to micafungin (MCF), anidulafungin (ANF) and caspofungin (CAS) is summarised using the MIC, area under the growth–concentration curve (AUC) and supra-MIC growth (SMG), derived from broth dilution assays. Micafungin MICs determined by Etest are reported separately. All MIC values are expressed in μg $ml^{-1}$. Micafungin MICs for the *C. albicans* isolates were determined by Etest on medium supplemented with 20 μg $ml^{-1}$ doxycycline.

| Species | Clade | Isolate / stock ID | Relevant genotype | Parental background | MCF MIC (μg ml−1) | MCF AUC | MCF SMG | MCF MIC Etest | ANF MIC (μg ml−1) | ANF AUC | ANF SMG | CAS MIC (μg ml−1) | CAS AUC | CAS SMG | Source / reference |
|---|---|---|---|---|---|---|---|---|---|---|---|---|---|---|---|
| *C. auris* | I | B8441 wt | wt | - | 0.5 | 130.79 | 0.062 | 0.250 | 0.5 | 53.49 | 0.011 | 1 | 585.42 | 0.387 | 2 |
| *C. auris* | I | NAT-marked wt | NAT-CauNEUT | B8441 | 0.5 | 128.63 | 0.102 | 0.250 | 0.5 | 49.85 | 0.084 | 1 | 503.33 | 0.356 | 3 |
| *C. auris* | I | *sln1Δ* | *sln1Δ* | B8441 | 0.2 | 169.40 | 0.184 | 0.125 | 1 | 599.10 | 0.335 | 32 | 1074.00 | NA | This study |
| *C. auris* | I | *nik1Δ* | *nik1Δ* | B8441 | 0.1 | 14.16 | 0.000 | 0.125 | 0.5 | 62.39 | 0.016 | 0.125 | 25.19 | 0.022 | This study |
| *C. auris* | I | *ssk1Δ* | *ssk1Δ* | B8441 | 16.0 | 889.70 | NA | 0.125 | 1 | 401.50 | 0.223 | 32 | 1336.00 | NA | This study |
| *C. auris* | I | *ssk2Δ* | *ssk2Δ* | B8441 | 16.0 | 1061.00 | NA | 0.125 | 1 | 609.90 | 0.368 | 32 | 1957.00 | NA | This study |
| *C. auris* | I | *pbs2Δ* | *pbs2Δ* | B8441 | 0.5 | 652.30 | 0.424 | 0.125 | 1 | 347.40 | 0.202 | 32 | 1636.00 | NA | This study |
| *C. auris* | I | *hog1Δ* | *hog1Δ* | B8441 | 2.0 | 699.50 | 0.416 | 0.940 | 1 | 282.80 | 0.154 | 32 | 1020.00 | NA | This study |
| *C. auris* | I | *mkc1Δ* | *mkc1Δ* | B8441 | 0.1 | 15.82 | 0.000 | 0.008 | 0.5 | 104.90 | 0.013 | 1 | 83.91 | 0.006 | This study |
| *C. auris* | I | *cek1Δ* | *cek1Δ* | B8441 | 0.1 | 16.55 | 0.000 | 0.047 | 0.5 | 55.05 | 0.017 | 2 | 478.20 | 0.222 | This study |
| *C. auris* | I | *hog1Δ mkc1Δ* | *hog1Δ mkc1Δ* | B8441 | - | - | - | 0.023 | - | - | - | - | - | - | This study |
| *C. auris* | I | Evolved 13 | *NIK1* I226N | B8441 | 1.0 | 488.76 | 0.305 | 0.190 | 2 | 543.58 | 0.301 | 32 | 1043.01 | NA | This study |
| *C. auris* | I | Evolved 19 | *NIK1* I226N | B8441 | 2.0 | 523.22 | 0.287 | 0.190 | 1 | 358.00 | 0.196 | 32 | 1052.53 | NA | This study |
| *C. auris* | I | Evolved 22 | *NIK1* H739Y | B8441 | 0.5 | 566.82 | 0.356 | 0.500 | 1 | 636.79 | 0.368 | 32 | 1472.66 | NA | This study |
| *C. auris* | I | Evolved 157 | *NIK1* L306I | B8441 | 4.0 | 634.08 | 0.325 | 0.500 | 1 | 451.74 | 0.254 | 32 | 1101.39 | NA | This study |
| *C. auris* | I | Evolved 218 | *NIK1* L306I | B8441 | 2.0 | 581.94 | 0.323 | 0.380 | 1 | 330.99 | 0.181 | 32 | 1025.13 | NA | This study |
| *C. auris* | I | Evolved 221 | *NIK1* T265I | B8441 | 4.0 | 646.08 | 0.317 | 0.380 | 1 | 565.30 | 0.328 | 32 | 1243.82 | NA | This study |
| *C. auris* | I | Evolved 223 | *NIK1* E472K | B8441 | 8.0 | 721.14 | 0.179 | 0.250 | 1 | 421.12 | 0.229 | 32 | 1043.38 | NA | This study |
| *C. auris* | I | Evolved 227 | *NIK1* K196N | B8441 | 4.0 | 614.17 | 0.269 | 0.250 | 1 | 402.25 | 0.215 | 32 | 1108.65 | NA | This study |
| *C. auris* | I | Evolved 233 | *NIK1* E472K | B8441 | 4.0 | 669.41 | 0.337 | 0.250 | 1 | 567.86 | 0.333 | 32 | 1242.37 | NA | This study |
| *C. auris* | I | Evolved 252 | *NIK1* I279V | B8441 | 4.0 | 710.57 | 0.382 | 0.125 | 1 | 526.07 | 0.307 | 32 | 1033.72 | NA | This study |
| *C. auris* | I | Evolved 17 | *SSK1* G525* | B8441 | 0.1 | 372.81 | 0.302 | 0.125 | 1 | 324.83 | 0.180 | 32 | 1126.69 | NA | This study |
| *C. auris* | I | Evolved 213 | *SSK1* S439* | B8441 | 0.1 | 339.00 | 0.297 | 0.250 | 0.5 | 272.34 | 0.158 | 32 | 1020.52 | NA | This study |
| *C. auris* | I | Evolved 220 | *SSK1* S439* | B8441 | 0.1 | 493.21 | 0.358 | 0.250 | 0.5 | 294.56 | 0.168 | 32 | 1194.59 | NA | This study |
| *C. auris* | I | Evolved 229 | *SSK1* S439* | B8441 | 0.1 | 417.60 | 0.319 | 0.125 | 1 | 261.45 | 0.137 | 32 | 1141.35 | NA | This study |
| *C. auris* | I | Evolved 254 | *SSK1* Q84* | B8441 | 0.5 | 549.71 | 0.353 | 0.190 | 1 | 689.46 | 0.402 | 32 | 1421.01 | NA | This study |
| *C. auris* | I | Evolved 257 | *SSK1* Q84* | B8441 | 0.1 | 365.84 | 0.291 | 0.190 | 0.25 | 264.37 | 0.183 | 32 | 1144.54 | NA | This study |

| Species | Clade | Isolate / stock ID | Relevant genotype | Parental background | MCF MIC (µg ml−1) | MCF AUC | MCF SMG | MCF MIC Etest | ANF MIC (µg ml−1) | ANF AUC | ANF SMG | CAS MIC (µg ml−1) | CAS AUC | CAS SMG | Source / reference |
|---|---|---|---|---|---|---|---|---|---|---|---|---|---|---|---|
| *C. auris* | I | Evolved 16 | *SSK2* G1096C | B8441 | 0.1 | 389.38 | 0.303 | 0.250 | 1 | 260.23 | 0.132 | 32 | 942.42 | NA | This study |
| *C. auris* | I | Evolved 101 | *SSK2* H1141/ | B8441 | 0.1 | 324.57 | 0.259 | 0.250 | 1 | 253.59 | 0.134 | 32 | 1047.69 | NA | This study |
| *C. auris* | I | Evolved 214 | *SSK2* E610G | B8441 | 0.1 | 522.46 | 0.373 | 0.380 | 1 | 289.74 | 0.156 | 32 | 1296.52 | NA | This study |
| *C. auris* | I | Evolved 230 | *SSK2* E324* | B8441 | 0.1 | 360.96 | 0.282 | 0.125 | 1 | 256.61 | 0.138 | 32 | 1021.90 | NA | This study |
| *C. auris* | I | Evolved 236 | *SSK2* S223/ | B8441 | 0.1 | 337.88 | 0.262 | 0.250 | 1 | 259.04 | 0.134 | 32 | 1079.20 | NA | This study |
| *C. auris* | I | Evolved 232 | *PBS2* Y294* | B8441 | 0.1 | 237.00 | 0.212 | 0.125 | 0.25 | 269.69 | 0.187 | 32 | 1016.46 | NA | This study |
| *C. auris* | I | Evolved 237 | *PBS2* Y294* | B8441 | 0.1 | 103.57 | 0.077 | 0.094 | 0.0625 | 99.15 | 0.093 | 0.125 | 610.51 | 0.431 | This study |
| *C. auris* | I | Evolved 235 | *HOG1* G5* | B8441 | 0.1 | 330.59 | 0.268 | 0.125 | 1 | 269.85 | 0.143 | 32 | 1010.39 | NA | This study |
| *C. auris* | I | Evolved 152 | *FKS1* S639Y | B8441 | 16.0 | 1235.35 | NA | NA | 32 | 1587.87 | NA | 32 | 1753.07 | NA | This study |
| *C. auris* | III | B11222 | wt | - | 0.1 | 221.50 | 0.173 | 0.125 | 0.0625 | 145.40 | 0.167 | 0.5 | 334.60 | 0.201 | 2 |
| *C. auris* | III | hog1Δ | *hog1Δ* | B11222 | 0.1 | 212.60 | 0.172 | 0.125 | 0.0625 | 90.98 | 0.148 | 0.25 | 316.60 | 0.220 | This study |
| *C. auris* | III | mkc1Δ | *mkc1Δ* | B11222 | 0.1 | 19.51 | 0.048 | 0.023 | 0.0625 | 27.51 | 0.032 | 0.125 | 23.97 | 0.018 | This study |
| *C. auris* | IV | B11245 | wt | - | 0.1 | 12.97 | 0.017 | 0.125 | 0.0625 | 24.35 | 0.047 | 0.25 | 639.90 | 0.299 | 2 |
| *C. auris* | IV | hog1Δ | *hog1Δ* | B11245 | 0.1 | 20.99 | 0.043 | 0.023 | 0.0625 | 37.01 | 0.053 | 0.125 | 395.60 | 0.213 | This study |
| *C. auris* | IV | mkc1Δ | *mkc1Δ* | B11245 | 0.1 | 26.48 | 0.020 | 0.023 | 0.0625 | 14.75 | 0.042 | 0.0625 | 32.75 | 0.104 | This study |
| *C. albicans* | I | CaSS1 | wt | SC5314 | - | - | - | 0.008 | - | - | - | - | - | - | 4 |
| *C. albicans* | I | hog1Δ | HOG1/*hog1Δ* | SC5314 | - | - | - | 0.023 | - | - | - | - | - | - | |
| *C. albicans* | I | mkc1Δ | MKC1/*mkc1Δ* | SC5314 | - | - | - | 0.008 | - | - | - | - | - | - | |
| *N. glabratus* | I | ATCC2001 wt | wt | - | - | - | - | 0.012 | - | - | - | - | - | - | 5 |
| *N. glabratus* | I | ssk1Δ | *ssk1Δ* | ATCC2001 | - | - | - | 0.016 | - | - | - | - | - | - | |
| *N. glabratus* | I | ssk2Δ | *ssk2Δ* | ATCC2001 | - | - | - | 0.012 | - | - | - | - | - | - | |
| *N. glabratus* | I | pbs2Δ | *pbs2Δ* | ATCC2001 | - | - | - | 0.008 | - | - | - | - | - | - | |
| *N. glabratus* | I | mkc1Δ | *mkc1Δ* | ATCC2001 | - | - | - | 0.008 | - | - | - | - | - | - | |

**Supplementary Table 2: Genetic variation in 25 selected evolved *C. auris* isolates identified by whole-genome sequencing.** Variants were identified relative to the sequenced clade I parental strain B8441. Gene identifiers refer to the systematic annotation of the B8441 reference genome. Gene names marked with an asterisk (*) correspond to *C. albicans* orthologue names used where no standard *C. auris* gene name was available. Nucleotide change indicates the reference and mutant alleles at the affected codon(s). Protein position indicates the affected amino-acid residue or residue range. Amino-acid changes are reported as the reference residue(s) followed by the mutant residue(s). Frameshift mutations are denoted by X, whereas premature stop codons are denoted by * in the 'aa change' column. ESM scores provide a sequence-based estimate of the predicted functional impact of protein-altering variants, with more negative values indicating substitutions that are less compatible with the sequence constraints of the respective protein and therefore potentially more disruptive. ESM scores are not reported for frameshift variants, as the multiresidue ESM approach has not been validated for these variants.

| Strain no. | Evolved in | Gene | Chromosome | Position | Nucleotide change | Gene name | Protein position | aa change | ESM score |
|---|---|---|---|---|---|---|---|---|---|
| 13 | ANF | B9J08_001959 | PEKT02000005 | 40415 | T/A | NIK1* | 226 | I/N | -5.49 |
| | | B9J08_000829 | PEKT02000002 | 708032 | ATACGTGAGCAAGAACAAGTTCTTGAGCACGTCACACTGGGCGTGCACCAGAGAATG/A | TRY5* | 327-345 | HSSVHAQCDVLKNLFLLTY/X | |
| 19 | ANF | B9J08_001959 | PEKT02000005 | 40415 | T/A | NIK1* | 226 | I/N | -5.49 |
| | | B9J08_000829 | PEKT02000002 | 708032 | ATACGTGAGCAAGAACAAGTTCTTGAGCACGTCACACTGGGCGTGCACCAGAGAATG/A | TRY5* | 327-345 | HSSVHAQCDVLKNLFLLTY/X | |
| 22 | ANF | B9J08_001296 | PEKT02000003 | 493880 | C/A | FAS1 | 239 | P/T | |
| | | B9J08_001959 | PEKT02000005 | 41953 | C/T | NIK1* | 739 | H/Y | -5.02 |
| | | B9J08_000829 | PEKT02000002 | 708032 | ATACGTGAGCAAGAACAAGTTCTTGAGCACGTCACACTGGGCGTGCACCAGAGAATG/A | TRY5* | 327-345 | HSSVHAQCDVLKNLFLLTY/X | |
| 157 | MCF | B9J08_002676 | PEKT02000007 | 215643 | GA/G | BNR1* | 650 | E/X | |
| | | B9J08_001959 | PEKT02000005 | 40654 | C/A | NIK1* | 306 | L/I | -9.13 |
| 218 | MCF | B9J08_002676 | PEKT02000007 | 215643 | GA/G | BNR1* | 650 | E/X | |
| | | B9J08_001959 | PEKT02000005 | 40654 | C/A | NIK1* | 306 | L/I | -9.13 |
| 221 | MCF | B9J08_002676 | PEKT02000007 | 215643 | GA/G | BNR1* | 650 | E/X | |
| | | B9J08_000199 | PEKT02000001 | 433366 | C/A | - | 702 | S/Y | |
| | | B9J08_001959 | PEKT02000005 | 40532 | C/T | NIK1* | 265 | T/I | -5.65 |
| 223 | MCF | B9J08_002676 | PEKT02000007 | 215643 | GA/G | BNR1* | 650 | E/X | |
| | | B9J08_001959 | PEKT02000005 | 41152 | G/A | NIK1* | 472 | E/K | -8.37 |
| 227 | CAS | B9J08_002676 | PEKT02000007 | 215643 | GA/G | BNR1* | 650 | E/X | |
| | | B9J08_001959 | PEKT02000005 | 40326 | A/T | NIK1* | 196 | K/N | -0.75 |
| 233 | CAS | B9J08_002676 | PEKT02000007 | 215643 | GA/G | BNR1* | 650 | E/X | |
| | | B9J08_001959 | PEKT02000005 | 41152 | G/A | NIK1* | 472 | E/K | -8.37 |
| 252 | ANF | B9J08_001959 | PEKT02000005 | 40573 | A/G | NIK1* | 279 | I/V | -6.20 |
| | | B9J08_000829 | PEKT02000002 | 708032 | ATACGTGAGCAAGAACAAGTTCTTGAGCACGTCACACTGGGCGTGCACCAGAGAATG/A | TRY5* | 327-345 | HSSVHAQCDVLKNLFLLTY/X | |
| 17 | ANF | B9J08_003819 | PEKT02000007 | 2558663 | T/G | GLR1 | 128 | F/V | |
| | | B9J08_000711 | PEKT02000002 | 443800 | C/A | RPP0 | 200 | S/Y | |
| | | B9J08_005450 | PEKT02000010 | 1230551 | C/A | SSK1 | 525 | G/* | -22.62 |
| 213 | MCF | B9J08_002676 | PEKT02000007 | 215643 | GA/G | BNR1* | 650 | E/X | |
| | | B9J08_004796 | PEKT02000009 | 763554 | C/T | - | 43 | M/I | |
| | | B9J08_002227 | PEKT02000005 | 615522 | C/T | - | 6 | R/H | |

| Strain no. | Evolved in | Gene | Chromosome | Position | Nucleotide change | Gene name | Protein position | aa change | ESM score |
|---|---|---|---|---|---|---|---|---|---|
| | | B9J08_005450 | PEKT02000010 | 1230808 | G/T | ***SSK1*** | 439 | S/* | -23.91 |
| 220 | MCF | B9J08_002676 | PEKT02000007 | 215643 | GA/G | *BNR1** | 650 | E/X | |
| | | B9J08_005450 | PEKT02000010 | 1230808 | G/T | ***SSK1*** | 439 | S/* | -23.91 |
| 229 | CAS | B9J08_002676 | PEKT02000007 | 215643 | GA/G | *BNR1** | 650 | E/X | |
| | | B9J08_005450 | PEKT02000010 | 1230808 | G/T | ***SSK1*** | 439 | S/* | -23.91 |
| 254 | ANF | B9J08_005450 | PEKT02000010 | 1231874 | G/A | ***SSK1*** | 84 | Q/* | -23.91 |
| 257 | ANF | B9J08_005450 | PEKT02000010 | 1231874 | G/A | ***SSK1*** | 84 | Q/* | -23.91 |
| 16 | ANF | B9J08_005216 | PEKT02000010 | 679734 | G/T | ***SSK2*** | 1096 | G/C | -15.42 |
| | | B9J08_000829 | PEKT02000002 | 708032 | ATACGTGAGCAAGAACAAGTTCTTGAGCACGTCACACTGGGCGTGCACCAGAGAATG/A | *TRY5** | 327-345 | HSSVHAQCDVLKNLFLLTY/X | |
| | | B9J08_004888 | PEKT02000009 | 970961 | G/T | - | 225 | L/F | |
| 101 | CAS | B9J08_005216 | PEKT02000010 | 679870 | AT/A | ***SSK2*** | 1141 | H/X | |
| 214 | MCF | B9J08_002676 | PEKT02000007 | 215643 | GA/G | *BNR1** | 650 | E/X | |
| | | B9J08_005216 | PEKT02000010 | 678277 | A/G | ***SSK2*** | 610 | E/G | -5.84 |
| 230 | CAS | B9J08_002676 | PEKT02000007 | 215643 | GA/G | *BNR1** | 650 | E/X | |
| | | B9J08_005216 | PEKT02000010 | 677418 | G/T | ***SSK2*** | 324 | E/* | -23.98 |
| 236 | CAS | B9J08_002676 | PEKT02000007 | 215643 | GA/G | *BNR1** | 650 | E/X | |
| | | B9J08_005216 | PEKT02000010 | 677115 | TC/T | ***SSK2*** | 223 | S/X | |
| 232 | CAS | B9J08_002676 | PEKT02000007 | 215643 | GA/G | *BNR1** | 650 | E/X | |
| | | B9J08_001824 | PEKT02000004 | 625581 | G/T | ***PBS2*** | 294 | Y/* | -23.30 |
| 237 | CAS | B9J08_002676 | PEKT02000007 | 215643 | GA/G | *BNR1** | 650 | E/X | |
| | | B9J08_001824 | PEKT02000004 | 625581 | G/T | ***PBS2*** | 294 | Y/* | -23.30 |
| 235 | CAS | B9J08_002676 | PEKT02000007 | 215643 | GA/G | *BNR1** | 650 | E/X | |
| | | B9J08_004369 | PEKT02000008 | 656997 | C/A | ***HOG1*** | 5 | G/* | -23.60 |
| 152 | MCF | B9J08_002676 | PEKT02000007 | 215643 | GA/G | *BNR1** | 650 | E/X | |
| | | B9J08_000964 | PEKT02000002 | 1006624 | G/T | ***FKS1*** | 639 | S/Y | -11.50 |

**Supplementary Table 3: $^{13}$C chemical-shift assignments of carbohydrates in intact *C. auris* cells.** Chemical shifts are reported in parts per million (ppm) for wt and *hog1Δ* cells grown under drug-free conditions or with 2 μg mL$^{-1}$ caspofungin (CAS). Assignments were obtained from two-dimensional DARR and DP-INADEQUATE spectra (**Supplementary Fig. 8**), as indicated. C1–C6 denote carbohydrate carbon positions; $CH_3$ and CO denote acetyl methyl and carbonyl carbons, respectively. Numbers following carbohydrate names distinguish separately resolved resonance sets. Sample annotations indicate whether assignments were observed in all samples or only in *hog1Δ* cells.

| | C1 | C2 | C3 | C4 | C5 | C6 | CH3 | CO | | |
|---|---|---|---|---|---|---|---|---|---|---|
| Chitin | 104.37 | 55.92 | 74.24 | 83.66 | 76.06 | 60.67 | 23.58 | 175.37 | All samples | DARR |
| | | | | | | | 22.8 | 174.1 | | |
| | | | | | | | 23.1 | 173.9 | Only hog1Δ | |
| β-1,3-glucan | 104.16 | 74.88 | 87.39 | 68.75 | 77.98 | 61.87 | | | All samples with different concentration | |
| β-1,6-glucan | 102.9 | 73.6 | 76.3 | 70.61 | 74.81 | 69.52 | | | | |
| α-1,2-mannan | 103.21 | 79.5 | 71.16 | 68.3 | 74.06 | 62.08 | | | | |
| β-1,3,6-glucan | 102.04 | 73.07 | 85.75 | 68.58 | 75.59 | 70.07 | | | | DP INADEQUATE |
| β-1,6-glucan | 104.16 | 74.47 | 76.2 | 70.24 | 75.2 | 69.35 | | | | |
| β-1,3-glucan 1 | 104.15 | 74.2 | 85.93 | 68.29 | 77.48 | 62.35 | | | | |
| β-1,3-glucan 2 | 104.12 | 74.44 | 85.6 | 68.36 | 77.2 | 62.41 | | | | |
| α-1,2-mannan 1 | 101.81 | 79.61 | 71.49 | 67.39 | 73.37 | 62.5 | | | | |
| α-1,2-mannan 2 | 100.26 | 78.95 | 71.19 | 67.98 | 74.17 | 61.95 | | | | |
| α-1,2-mannan 3 | 99.37 | 79.83 | 71.39 | 67.52 | 74.24 | 62.3 | | | | |
| α-1,6-mannan 1 | 103.26 | 70.96 | 73.34 | 68.11 | 71.18 | 67.01 | | | | |
| α-1,6-mannan 2 | 101.13 | 70.97 | 73.22 | 68.53 | 71.11 | 66.14 | | | | |
| α-1,6-mannan 3 | 100.92 | 71.58 | 74.09 | 68.58 | 71.18 | 67.56 | | | | |
| Galactose 1 | 97.32 | 75.69 | 77.04 | 70.45 | 72.56 | 62.3 | | | | |
| Galactose 2 | 97.41 | 71.94 | 77.54 | 72.74 | 74.44 | 62.24 | | | | |
| Glucose | 93.46 | 73.04 | 74.64 | 71.54 | 72.53 | 62.8 | | | | |

**Supplementary Table 4: Differential protein abundance, selected biological-process assignments and heatmap inclusion.** Protein-level results for 699 proteins showing |$\log_2$ fold change| > 1 and an adjusted P value < 0.05 in at least one comparison. $\text{Log}_2$ fold changes and Benjamini–Hochberg-adjusted P values are provided for CAS-treated versus untreated wt, CAS-treated versus untreated *hog1Δ*, *hog1Δ* versus wt under CAS exposure, and *hog1Δ* versus wt without CAS (ND = no drug). Positive $\log_2$ fold changes indicate higher abundance in the first condition named. ‘All processes’ lists assignments to the selected GO biological-process categories. Blank entries indicate no assignment to these categories and do not necessarily indicate an absence of GO annotations. ‘Included in Fig. 3e’ identifies proteins displayed in the heatmap. Blank quantitative entries indicate unavailable results, and non-finite fold-change estimates are retained as Inf or −Inf.

| Gene | Symbol | All processes | Included in Fig. 3e | wt-CAS vs wt-ND log2(FC) | wt-CAS vs wt-ND p(adj) | *hog1*Δ-CAS vs hog1Δ-ND log2(FC) | *hog1*Δ-CAS vs hog1Δ-ND p(adj) | *hog1*Δ-CAS vs wt-CAS log2(FC) | *hog1*Δ-CAS vs wt-CAS p(adj) | *hog1*Δ-ND vs wt-ND log2(FC) | *hog1*Δ-ND vs wt-ND p(adj) |
|---|---|---|---|---|---|---|---|---|---|---|---|
| B9J08_000116 | | | FALSE | 5,407 | 0,015 | 5,240 | 0,055 | -0,963 | 0,029 | -0,796 | 0,835 |
| B9J08_002829 | *DBF2* | Cytoskeleton organization; DNA replication / cell cycle | TRUE | 4,568 | 0,031 | Inf | 0,508 | -2,857 | 0,061 | -Inf | 0,627 |
| B9J08_002990 | | | FALSE | 4,556 | 0,032 | 2,411 | 0,508 | -1,906 | 0,144 | 0,239 | 0,957 |
| B9J08_004381 | *UAP1* | | FALSE | 4,365 | 0,034 | 1,477 | 0,012 | -2,644 | 0,019 | 0,244 | 0,385 |
| B9J08_002443 | *GNP2* | Transmembrane transport | TRUE | 4,239 | 0,029 | | | -Inf | 0,024 | -Inf | 0,627 |
| B9J08_000890 | | | FALSE | 3,857 | 0,040 | 0,939 | 0,356 | -1,105 | 0,068 | 1,813 | 0,561 |
| B9J08_004410 | *CRH11* | Cell wall organization / biogenesis; Carbohydrate metabolism | TRUE | 3,610 | 0,046 | | | -Inf | 0,033 | -Inf | 0,627 |
| B9J08_005294 | | | FALSE | 3,575 | 0,008 | -0,098 | 0,607 | -3,593 | 0,010 | 0,081 | 0,793 |
| B9J08_005249 | *GIN4* | Cytoskeleton organization; DNA replication / cell cycle | TRUE | 3,279 | 0,046 | -1,483 | 0,658 | -4,152 | 0,026 | 0,610 | 0,866 |
| B9J08_003467 | *CDC54* | DNA replication / cell cycle | TRUE | 3,093 | 0,005 | 1,630 | 0,138 | -2,169 | 0,008 | -0,706 | 0,630 |
| B9J08_001559 | | Transmembrane transport | TRUE | 3,004 | 0,013 | -0,202 | 0,641 | -2,670 | 0,018 | 0,536 | 0,502 |
| B9J08_005433 | | | FALSE | 2,766 | 0,014 | -2,142 | 0,053 | -4,461 | 0,012 | 0,447 | 0,495 |
| B9J08_000663 | | | FALSE | 2,694 | 0,023 | Inf | 0,508 | -5,864 | 0,010 | -Inf | 0,319 |
| B9J08_000765 | *CAS5* | Cell wall organization / biogenesis; Lipid metabolism; Response to stress | TRUE | 2,642 | 0,049 | 2,188 | 0,085 | -0,894 | 0,074 | -0,440 | 0,790 |
| B9J08_001239 | | | FALSE | 2,501 | 0,036 | 0,698 | 0,079 | -1,119 | 0,026 | 0,684 | 0,313 |
| B9J08_005476 | *MCM2* | DNA replication / cell cycle | TRUE | 2,411 | 0,031 | 1,576 | 0,126 | -1,320 | 0,028 | -0,485 | 0,721 |
| B9J08_003331 | | Translation / ribosome biogenesis | TRUE | 2,362 | 0,047 | 0,516 | 0,518 | -0,650 | 0,308 | 1,196 | 0,286 |
| B9J08_004413 | *RLP24* | | FALSE | 2,331 | 0,039 | 0,224 | 0,508 | -1,063 | 0,010 | 1,045 | 0,318 |
| B9J08_002262 | *KRE6a* | Cell wall organization / biogenesis; Carbohydrate metabolism | TRUE | 2,205 | 0,032 | 0,463 | 0,253 | -1,605 | 0,014 | 0,136 | 0,783 |
| B9J08_002379 | | Protein folding | TRUE | 2,175 | 0,028 | -0,280 | 0,845 | -3,377 | 0,018 | -0,922 | 0,554 |
| B9J08_000777 | | | FALSE | 2,111 | 0,015 | -Inf | 0,156 | -Inf | 0,018 | 0,954 | 0,483 |
| B9J08_000735 | *BCK1* | Cell wall organization / biogenesis | TRUE | 2,087 | 0,031 | 2,283 | 0,050 | -0,448 | 0,119 | -0,644 | 0,286 |
| B9J08_003486 | *PLB3* | Lipid metabolism | TRUE | 2,054 | 0,014 | -0,329 | 0,197 | -2,003 | 0,011 | 0,380 | 0,340 |
| B9J08_002497 | *CDC28* | Cytoskeleton organization; Protein catabolism; Response to stress; DNA replication / cell cycle | TRUE | 2,045 | 0,045 | 0,514 | 0,164 | -1,429 | 0,018 | 0,102 | 0,758 |
| B9J08_000878 | | Translation / ribosome biogenesis | TRUE | 2,014 | 0,034 | Inf | 0,102 | -2,628 | 0,018 | -Inf | 0,268 |
| B9J08_002555 | | | FALSE | 1,995 | 0,030 | 1,796 | 0,146 | 0,119 | 0,800 | 0,318 | 0,657 |

| Gene | Symbol | All processes | Included in Fig. 3e | wt-CAS vs wt-ND log2(FC) | wt-CAS vs wt-ND p(adj) | *hog1*Δ-CAS vs hog1Δ-ND log2(FC) | *hog1*Δ-CAS vs hog1Δ-ND p(adj) | *hog1*Δ-CAS vs wt-CAS log2(FC) | *hog1*Δ-CAS vs wt-CAS p(adj) | *hog1*Δ-ND vs wt-ND log2(FC) | *hog1*Δ-ND vs wt-ND p(adj) |
|---|---|---|---|---|---|---|---|---|---|---|---|
| B9J08_004701 | *MDJ1* | Protein folding | TRUE | 1,898 | 0,028 | 0,095 | 0,607 | -1,527 | 0,010 | 0,276 | 0,255 |
| B9J08_004973 | | | FALSE | 1,888 | 0,034 | -0,375 | 0,053 | -1,260 | 0,039 | 1,003 | 0,255 |
| B9J08_003981 | *MDR1* | Response to stress; Transmembrane transport | TRUE | 1,864 | 0,046 | -0,496 | 0,240 | -1,399 | 0,025 | 0,961 | 0,274 |
| B9J08_004661 | *SWI4* | DNA replication / cell cycle | TRUE | 1,730 | 0,022 | 1,138 | 0,062 | -0,487 | 0,098 | 0,104 | 0,863 |
| B9J08_000233 | *RPL7* | | FALSE | 1,676 | 0,037 | 0,087 | 0,654 | -0,942 | 0,025 | 0,647 | 0,286 |
| B9J08_002537 | | | FALSE | 1,629 | 0,046 | -0,443 | 0,508 | -1,908 | 0,019 | 0,164 | 0,833 |
| B9J08_005185 | *JEM1* | Protein folding | TRUE | 1,616 | 0,032 | 0,396 | 0,529 | -0,672 | 0,245 | 0,548 | 0,301 |
| B9J08_003641 | | | FALSE | 1,615 | 0,045 | 0,092 | 0,706 | -0,709 | 0,050 | 0,814 | 0,300 |
| B9J08_002033 | *REX3* | | FALSE | 1,601 | 0,042 | 0,364 | 0,140 | -0,178 | 0,301 | 1,059 | 0,235 |
| B9J08_003935 | *MRPS9* | Translation / ribosome biogenesis; Cellular respiration | TRUE | 1,572 | 0,028 | -0,079 | 0,293 | -1,302 | 0,022 | 0,350 | 0,409 |
| B9J08_001901 | *UTR2* | Cell wall organization / biogenesis; Carbohydrate metabolism | TRUE | 1,570 | 0,032 | 0,411 | 0,370 | -2,552 | 0,011 | -1,393 | 0,185 |
| B9J08_002719 | *RPL30* | | FALSE | 1,546 | 0,031 | 2,438 | 0,020 | 1,082 | 0,053 | 0,190 | 0,813 |
| B9J08_003249 | *RPL13* | Translation / ribosome biogenesis | TRUE | 1,542 | 0,040 | -0,229 | 0,313 | -0,869 | 0,030 | 0,902 | 0,246 |
| B9J08_000457 | | | FALSE | 1,528 | 0,032 | 0,316 | 0,051 | -0,979 | 0,037 | 0,233 | 0,507 |
| B9J08_001128 | | Translation / ribosome biogenesis | TRUE | 1,526 | 0,034 | -0,988 | 0,155 | -2,668 | 0,017 | -0,155 | 0,753 |
| B9J08_002961 | *RPS8A* | Translation / ribosome biogenesis | TRUE | 1,511 | 0,042 | 2,272 | 0,032 | 0,648 | 0,037 | -0,114 | 0,577 |
| B9J08_005330 | | | FALSE | 1,494 | 0,047 | 1,069 | 0,068 | 0,387 | 0,230 | 0,812 | 0,286 |
| B9J08_000700 | | | FALSE | 1,444 | 0,034 | -1,027 | 0,245 | -2,271 | 0,022 | 0,199 | 0,774 |
| B9J08_002506 | | | FALSE | 1,404 | 0,034 | -1,206 | 0,045 | -1,895 | 0,013 | 0,715 | 0,255 |
| B9J08_004640 | *SIM1* | Cell wall organization / biogenesis; DNA replication / cell cycle | TRUE | 1,401 | 0,022 | -0,290 | 0,171 | -1,595 | 0,010 | 0,095 | 0,672 |
| B9J08_002809 | | Translation / ribosome biogenesis | TRUE | 1,397 | 0,042 | -0,526 | 0,171 | -1,765 | 0,010 | 0,157 | 0,717 |
| B9J08_004083 | *INN1* | | FALSE | 1,384 | 0,029 | -1,325 | 0,076 | -2,160 | 0,006 | 0,549 | 0,330 |
| B9J08_004991 | | | FALSE | 1,374 | 0,044 | -0,427 | 0,308 | -1,222 | 0,043 | 0,579 | 0,317 |
| B9J08_001356 | *PEA2* | | FALSE | 1,362 | 0,036 | -0,497 | 0,134 | -1,480 | 0,018 | 0,379 | 0,185 |
| B9J08_002677 | *MIS12* | | FALSE | 1,358 | 0,024 | 0,084 | 0,750 | -1,820 | 0,018 | -0,546 | 0,255 |
| B9J08_002124 | *DED1* | Translation / ribosome biogenesis; Response to stress | TRUE | 1,329 | 0,031 | 0,342 | 0,046 | -0,839 | 0,021 | 0,148 | 0,422 |
| B9J08_005200 | | | FALSE | 1,296 | 0,044 | 0,244 | 0,731 | -0,515 | 0,416 | 0,537 | 0,286 |
| B9J08_000827 | | | FALSE | 1,295 | 0,047 | -1,506 | 0,203 | -2,428 | 0,010 | 0,373 | 0,647 |
| B9J08_001926 | | | FALSE | 1,294 | 0,042 | -0,871 | 0,013 | -2,075 | 0,025 | 0,090 | 0,791 |
| B9J08_004292 | *CAR1* | | FALSE | 1,263 | 0,037 | -0,094 | 0,514 | -0,918 | 0,020 | 0,440 | 0,261 |
| B9J08_002495 | *GUK1* | | FALSE | 1,243 | 0,011 | 0,338 | 0,104 | -0,527 | 0,026 | 0,379 | 0,323 |
| B9J08_002094 | | | FALSE | 1,240 | 0,032 | -0,285 | 0,106 | -0,975 | 0,012 | 0,549 | 0,235 |
| B9J08_005297 | *TIM50* | | FALSE | 1,219 | 0,041 | -0,442 | 0,037 | -1,078 | 0,025 | 0,582 | 0,266 |
| B9J08_001959 | *NIK1* | Response to stress | TRUE | 1,206 | 0,030 | -0,239 | 0,387 | -1,577 | 0,022 | -0,131 | 0,688 |
| B9J08_000267 | | | FALSE | 1,193 | 0,046 | 2,337 | 0,367 | -0,726 | 0,450 | -1,870 | 0,385 |
| B9J08_000523 | *OCT1* | Cellular respiration | TRUE | 1,184 | 0,049 | 1,395 | 0,139 | 0,015 | 0,970 | -0,197 | 0,617 |
| B9J08_005312 | *MRP17* | Translation / ribosome biogenesis | TRUE | 1,167 | 0,046 | 0,265 | 0,456 | -0,489 | 0,083 | 0,413 | 0,491 |
| B9J08_004671 | | | FALSE | 1,121 | 0,011 | 0,471 | 0,119 | 0,232 | 0,302 | 0,882 | 0,235 |
| B9J08_000771 | *RPS6A* | Translation / ribosome biogenesis | TRUE | 1,117 | 0,034 | -0,800 | 0,049 | -1,310 | 0,019 | 0,608 | 0,235 |
| B9J08_003526 | *MUC1* | | FALSE | 1,073 | 0,034 | 0,773 | 0,023 | 0,093 | 0,343 | 0,393 | 0,208 |
| B9J08_002598 | | | FALSE | 1,067 | 0,014 | 0,072 | 0,712 | -0,796 | 0,050 | 0,199 | 0,468 |
| B9J08_000439 | | | FALSE | 1,029 | 0,034 | 0,753 | 0,108 | -0,437 | 0,125 | -0,161 | 0,545 |

| Gene | Symbol | All processes | Included in Fig. 3e | wt-CAS vs wt-ND log2(FC) | wt-CAS vs wt-ND p(adj) | *hog1*Δ-CAS vs hog1Δ-ND log2(FC) | *hog1*Δ-CAS vs hog1Δ-ND p(adj) | *hog1*Δ-CAS vs wt-CAS log2(FC) | *hog1*Δ-CAS vs wt-CAS p(adj) | *hog1*Δ-ND vs wt-ND log2(FC) | *hog1*Δ-ND vs wt-ND p(adj) |
|---|---|---|---|---|---|---|---|---|---|---|---|
| B9J08_000476 | *HSP104* | Protein folding; Response to stress | TRUE | -1,020 | 0,042 | -0,434 | 0,030 | 0,318 | 0,119 | -0,268 | 0,286 |
| B9J08_002575 | | | FALSE | -1,100 | 0,034 | 0,364 | 0,159 | 1,148 | 0,045 | -0,316 | 0,143 |
| B9J08_000717 | *MSC3* | | FALSE | -1,173 | 0,034 | -0,922 | 0,021 | 0,684 | 0,046 | 0,432 | 0,208 |
| B9J08_004941 | | | FALSE | -1,174 | 0,039 | -1,140 | 0,030 | 0,232 | 0,301 | 0,198 | 0,375 |
| B9J08_003998 | *ACS1* | | FALSE | -1,513 | 0,031 | 0,012 | 0,964 | 1,093 | 0,100 | -0,432 | 0,208 |
| B9J08_000528 | *SOD2* | Response to stress | TRUE | -1,552 | 0,047 | -0,348 | 0,083 | 1,067 | 0,025 | -0,138 | 0,489 |
| B9J08_002479 | *ZTA1* | Response to stress | TRUE | -1,655 | 0,005 | -1,649 | 0,017 | -0,287 | 0,295 | -0,293 | 0,274 |
| B9J08_002683 | | | FALSE | -1,718 | 0,005 | -0,372 | 0,218 | 1,209 | 0,101 | -0,137 | 0,246 |
| B9J08_001591 | *GAD1* | | FALSE | -1,740 | 0,015 | -0,136 | 0,302 | 1,397 | 0,025 | -0,207 | 0,273 |
| B9J08_001987 | | | FALSE | -1,796 | 0,034 | -0,165 | 0,885 | 0,020 | 0,963 | -1,611 | 0,289 |
| B9J08_004003 | *DOT5* | | FALSE | -1,833 | 0,043 | -0,596 | 0,083 | 1,050 | 0,008 | -0,187 | 0,390 |
| B9J08_003314 | *RBE1* | | FALSE | -2,209 | 0,023 | -1,177 | 0,050 | 0,866 | 0,069 | -0,166 | 0,454 |
| B9J08_000483 | *HSP70* | Response to stress | TRUE | -2,229 | 0,018 | -1,748 | 0,013 | 0,096 | 0,651 | -0,385 | 0,208 |
| B9J08_004838 | | | FALSE | -2,665 | 0,039 | -1,470 | 0,106 | -0,186 | 0,727 | -1,381 | 0,161 |
| B9J08_002737 | | | FALSE | -2,743 | 0,047 | -2,079 | 0,062 | -0,254 | 0,205 | -0,918 | 0,185 |
| B9J08_003627 | *HSP21* | Response to stress | TRUE | -2,917 | 0,034 | -1,151 | 0,013 | 1,053 | 0,028 | -0,712 | 0,164 |
| B9J08_004015 | *GCA1* | Carbohydrate metabolism | TRUE | -3,007 | 0,046 | -2,084 | 0,029 | 1,146 | 0,295 | 0,224 | 0,453 |
| B9J08_004096 | | | FALSE | -3,708 | 0,042 | 0,974 | 0,049 | 4,856 | 0,010 | 0,174 | 0,665 |
| B9J08_003568 | *GCY1* | | FALSE | -3,866 | 0,034 | -Inf | 0,055 | -Inf | 0,513 | -0,322 | 0,409 |
| B9J08_001065 | *PRP45* | | FALSE | -3,928 | 0,044 | 1,515 | 0,470 | 4,101 | 0,312 | -1,342 | 0,517 |
| B9J08_001613 | *PHO100* | | FALSE | -4,344 | 0,047 | -1,433 | 0,013 | 3,289 | 0,053 | 0,378 | 0,226 |
| B9J08_004858 | | Lipid metabolism | TRUE | -Inf | 0,128 | 7,700 | 0,035 | Inf | 0,033 | 0,201 | 0,730 |
| B9J08_003902 | *EMC9* | | FALSE | 2,208 | 0,178 | 6,149 | 0,045 | 5,130 | 0,044 | 1,189 | 0,582 |
| B9J08_002856 | *CHS8* | | FALSE | 1,755 | 0,097 | 5,293 | 0,041 | 0,152 | 0,586 | -3,386 | 0,208 |
| B9J08_001340 | *RVS161* | Cell wall organization / biogenesis; Vesicle-mediated transport; Cytoskeleton organization; DNA replication / cell cycle | TRUE | 0,307 | 0,152 | 5,025 | 0,030 | 4,650 | 0,028 | -0,068 | 0,337 |
| B9J08_002140 | | | FALSE | -0,715 | 0,565 | 3,846 | 0,045 | 2,792 | 0,047 | -1,768 | 0,459 |
| B9J08_000894 | | | FALSE | 2,315 | 0,095 | 3,594 | 0,012 | 1,570 | 0,019 | 0,292 | 0,893 |
| B9J08_001079 | *WSC2* | | FALSE | 0,557 | 0,333 | 3,481 | 0,026 | 0,631 | 0,095 | -2,293 | 0,293 |
| B9J08_003727 | *SUV3* | Cellular respiration | TRUE | 0,116 | 0,863 | 3,336 | 0,030 | 1,007 | 0,058 | -2,213 | 0,339 |
| B9J08_004346 | *BUB1* | DNA replication / cell cycle | TRUE | Inf | 0,557 | 3,216 | 0,033 | 2,122 | 0,144 | Inf | 0,627 |
| B9J08_004820 | *TAC1b* | | FALSE | -0,801 | 0,663 | 2,998 | 0,045 | 2,104 | 0,141 | -1,695 | 0,425 |
| B9J08_002196 | *SAS2* | | FALSE | -2,655 | 0,156 | 2,924 | 0,043 | 4,843 | 0,052 | -0,736 | 0,627 |
| B9J08_001487 | | Transmembrane transport | TRUE | -1,018 | 0,557 | 2,881 | 0,019 | 4,469 | 0,015 | 0,570 | 0,627 |
| B9J08_002362 | | | FALSE | -0,183 | 0,890 | 2,809 | 0,022 | 2,297 | 0,058 | -0,695 | 0,598 |
| B9J08_002242 | | Transmembrane transport | TRUE | | | 2,775 | 0,045 | Inf | 0,020 | Inf | 0,513 |
| B9J08_004650 | *MNN26* | Cell wall organization / biogenesis; Carbohydrate metabolism | TRUE | 1,606 | 0,112 | 2,744 | 0,017 | 0,765 | 0,036 | -0,373 | 0,731 |
| B9J08_004987 | | | FALSE | | | 2,730 | 0,034 | Inf | 0,019 | Inf | 0,459 |
| B9J08_004150 | *CHS3* | | FALSE | 2,109 | 0,065 | 2,642 | 0,029 | -0,383 | 0,127 | -0,917 | 0,627 |
| B9J08_002643 | *RCT1* | | FALSE | 6,756 | 0,061 | 2,597 | 0,036 | -4,512 | 0,032 | -0,354 | 0,627 |
| B9J08_003545 | *TFA1* | | FALSE | -0,481 | 0,301 | 2,461 | 0,017 | 2,900 | 0,006 | -0,041 | 0,878 |
| B9J08_002399 | *SUP35* | Translation / ribosome biogenesis | TRUE | 0,442 | 0,174 | 2,441 | 0,034 | 2,066 | 0,026 | 0,067 | 0,590 |
| B9J08_003861 | *SPT5* | Response to stress | TRUE | 0,192 | 0,531 | 2,350 | 0,029 | 2,328 | 0,016 | 0,170 | 0,454 |
| B9J08_004754 | *HHT1* | | FALSE | -0,371 | 0,586 | 2,327 | 0,008 | 3,159 | 0,006 | 0,461 | 0,515 |

| Gene | Symbol | All processes | Included in Fig. 3e | wt-CAS vs wt-ND log2(FC) | wt-CAS vs wt-ND p(adj) | *hog1*Δ-CAS vs hog1Δ-ND log2(FC) | *hog1*Δ-CAS vs hog1Δ-ND p(adj) | *hog1*Δ-CAS vs wt-CAS log2(FC) | *hog1*Δ-CAS vs wt-CAS p(adj) | *hog1*Δ-ND vs wt-ND log2(FC) | *hog1*Δ-ND vs wt-ND p(adj) |
|---|---|---|---|---|---|---|---|---|---|---|---|
| B9J08_002632 | *GFA1* | | FALSE | 3,613 | 0,079 | 2,262 | 0,027 | -1,453 | 0,058 | -0,102 | 0,703 |
| B9J08_002681 | *MKC1* | DNA replication / cell cycle; Transmembrane transport | TRUE | 3,665 | 0,390 | 2,214 | 0,034 | -0,890 | 0,561 | 0,561 | 0,530 |
| B9J08_002612 | | | FALSE | 0,665 | 0,137 | 2,193 | 0,040 | 1,477 | 0,048 | -0,051 | 0,927 |
| B9J08_002670 | *NGS1* | Carbohydrate metabolism | TRUE | -0,253 | 0,841 | 2,140 | 0,009 | 2,876 | 0,028 | 0,483 | 0,536 |
| B9J08_002221 | | | FALSE | 1,250 | 0,074 | 2,076 | 0,017 | 0,968 | 0,021 | 0,142 | 0,758 |
| B9J08_001881 | | | FALSE | -0,659 | 0,452 | 2,065 | 0,008 | 2,269 | 0,028 | -0,455 | 0,598 |
| B9J08_003961 | | | FALSE | 0,614 | 0,384 | 2,045 | 0,030 | 1,763 | 0,050 | 0,332 | 0,745 |
| B9J08_000599 | | | FALSE | 1,681 | 0,192 | 2,011 | 0,031 | 0,653 | 0,188 | 0,323 | 0,759 |
| B9J08_001431 | | Vesicle-mediated transport | TRUE | 0,084 | 0,945 | 1,969 | 0,020 | 2,452 | 0,050 | 0,568 | 0,164 |
| B9J08_000579 | | | FALSE | -0,480 | 0,262 | 1,807 | 0,031 | 2,129 | 0,023 | -0,157 | 0,645 |
| B9J08_003259 | | | FALSE | -0,320 | 0,557 | 1,805 | 0,038 | 1,769 | 0,029 | -0,355 | 0,459 |
| B9J08_000011 | *ATG11* | Response to stress | TRUE | -0,378 | 0,294 | 1,792 | 0,027 | 2,553 | 0,025 | 0,383 | 0,421 |
| B9J08_000195 | | | FALSE | -0,425 | 0,339 | 1,785 | 0,039 | 1,625 | 0,022 | -0,586 | 0,385 |
| B9J08_000289 | | | FALSE | 0,366 | 0,505 | 1,783 | 0,025 | 1,640 | 0,021 | 0,222 | 0,627 |
| B9J08_000181 | *TAO3* | Cell wall organization / biogenesis; Response to stress | TRUE | -0,070 | 0,911 | 1,778 | 0,037 | 2,094 | 0,036 | 0,246 | 0,724 |
| B9J08_003957 | *MEA1* | | FALSE | -1,853 | 0,188 | 1,721 | 0,021 | 3,793 | 0,010 | 0,219 | 0,716 |
| B9J08_003755 | *RPS15* | Translation / ribosome biogenesis | TRUE | 0,230 | 0,600 | 1,720 | 0,012 | 1,458 | 0,029 | -0,032 | 0,943 |
| B9J08_001724 | *PXA2* | Transmembrane transport | TRUE | 0,784 | 0,228 | 1,695 | 0,025 | 0,907 | 0,054 | -0,004 | 0,995 |
| B9J08_002511 | | | FALSE | 1,505 | 0,111 | 1,693 | 0,044 | 0,409 | 0,245 | 0,221 | 0,808 |
| B9J08_003434 | *SPC72* | Cytoskeleton organization | TRUE | 0,214 | 0,443 | 1,631 | 0,028 | 1,501 | 0,026 | 0,084 | 0,794 |
| B9J08_001813 | | | FALSE | -0,184 | 0,644 | 1,612 | 0,018 | 1,880 | 0,012 | 0,084 | 0,797 |
| B9J08_003663 | *MDL2* | Transmembrane transport | TRUE | 0,719 | 0,249 | 1,599 | 0,041 | 1,385 | 0,025 | 0,506 | 0,286 |
| B9J08_002617 | *FLC1* | | FALSE | 0,389 | 0,470 | 1,596 | 0,031 | 1,297 | 0,045 | 0,090 | 0,875 |
| B9J08_005332 | | | FALSE | -0,126 | 0,846 | 1,585 | 0,040 | 1,265 | 0,049 | -0,446 | 0,611 |
| B9J08_005270 | | Vesicle-mediated transport | TRUE | -1,182 | 0,193 | 1,543 | 0,008 | 2,436 | 0,028 | -0,288 | 0,627 |
| B9J08_001150 | *VMA10* | Transmembrane transport | TRUE | -0,298 | 0,525 | 1,542 | 0,022 | 1,669 | 0,028 | -0,172 | 0,731 |
| B9J08_005478 | *PIM1* | Protein catabolism | TRUE | 0,490 | 0,136 | 1,532 | 0,038 | 0,644 | 0,042 | -0,398 | 0,179 |
| B9J08_002263 | *KRE6b* | Cell wall organization / biogenesis; Carbohydrate metabolism | TRUE | 1,244 | 0,119 | 1,531 | 0,042 | -0,114 | 0,554 | -0,401 | 0,634 |
| B9J08_002896 | *FUN31* | | FALSE | 0,488 | 0,612 | 1,529 | 0,039 | 1,476 | 0,010 | 0,435 | 0,721 |
| B9J08_002941 | *MED14* | | FALSE | 0,142 | 0,864 | 1,527 | 0,018 | 1,583 | 0,047 | 0,198 | 0,778 |
| B9J08_001933 | | | FALSE | 0,360 | 0,131 | 1,523 | 0,048 | 1,562 | 0,069 | 0,398 | 0,481 |
| B9J08_000371 | | | FALSE | -0,171 | 0,696 | 1,512 | 0,035 | 2,051 | 0,025 | 0,368 | 0,461 |
| PTR_B | | | FALSE | 1,033 | 0,399 | 1,505 | 0,028 | 1,500 | 0,054 | 1,028 | 0,301 |
| B9J08_003960 | *SPA2* | Cytoskeleton organization; DNA replication / cell cycle | TRUE | 0,524 | 0,021 | 1,486 | 0,013 | 0,996 | 0,022 | 0,034 | 0,669 |
| B9J08_004874 | *CGT1* | | FALSE | -0,329 | 0,320 | 1,477 | 0,017 | 1,700 | 0,013 | -0,106 | 0,483 |
| B9J08_001214 | | | FALSE | -1,371 | 0,257 | 1,473 | 0,014 | 2,187 | 0,067 | -0,657 | 0,462 |
| B9J08_001351 | | Cellular respiration | TRUE | -0,381 | 0,475 | 1,472 | 0,037 | 2,780 | 0,025 | 0,927 | 0,268 |
| B9J08_004117 | *LTV1* | | FALSE | 0,407 | 0,492 | 1,467 | 0,017 | 0,943 | 0,104 | -0,117 | 0,753 |
| B9J08_005522 | | | FALSE | 1,029 | 0,160 | 1,465 | 0,031 | 1,138 | 0,040 | 0,702 | 0,422 |
| B9J08_002923 | *SMD3* | | FALSE | 0,172 | 0,575 | 1,462 | 0,013 | 0,592 | 0,031 | -0,699 | 0,346 |
| B9J08_001897 | *CDC53* | Protein catabolism | TRUE | 0,506 | 0,337 | 1,459 | 0,026 | 1,140 | 0,028 | 0,187 | 0,735 |
| B9J08_000434 | *PET9* | Transmembrane transport | TRUE | 0,254 | 0,355 | 1,446 | 0,008 | 0,748 | 0,028 | -0,444 | 0,349 |

| Gene | Symbol | All processes | Included in Fig. 3e | wt-CAS vs wt-ND log2(FC) | wt-CAS vs wt-ND p(adj) | *hog1*Δ-CAS vs hog1Δ-ND log2(FC) | *hog1*Δ-CAS vs hog1Δ-ND p(adj) | *hog1*Δ-CAS vs wt-CAS log2(FC) | *hog1*Δ-CAS vs wt-CAS p(adj) | *hog1*Δ-ND vs wt-ND log2(FC) | *hog1*Δ-ND vs wt-ND p(adj) |
|---|---|---|---|---|---|---|---|---|---|---|---|
| B9J08_002423 | *STB3* | | FALSE | -0,969 | 0,034 | 1,443 | 0,029 | 2,493 | 0,027 | 0,081 | 0,627 |
| B9J08_005121 | *HMO1* | DNA replication / cell cycle | TRUE | 1,170 | 0,096 | 1,438 | 0,017 | 0,844 | 0,025 | 0,576 | 0,431 |
| B9J08_001178 | *MCI4* | Cellular respiration | TRUE | 0,585 | 0,257 | 1,438 | 0,040 | 0,153 | 0,609 | -0,700 | 0,354 |
| B9J08_003700 | *KOG1* | | FALSE | 0,307 | 0,448 | 1,435 | 0,022 | 1,366 | 0,020 | 0,238 | 0,410 |
| B9J08_002174 | *NAA25* | | FALSE | 0,630 | 0,235 | 1,433 | 0,021 | 0,937 | 0,021 | 0,134 | 0,827 |
| B9J08_003265 | *PDE2* | Cell wall organization / biogenesis; Carbohydrate metabolism; Response to stress | TRUE | -Inf | 0,240 | 1,427 | 0,017 | Inf | 0,022 | 0,471 | 0,622 |
| B9J08_001020 | *FKS2* | Carbohydrate metabolism | TRUE | 2,809 | 0,130 | 1,419 | 0,031 | -0,792 | 0,212 | 0,598 | 0,459 |
| B9J08_001001 | *CRL1* | | FALSE | 2,431 | 0,110 | 1,406 | 0,037 | -0,783 | 0,168 | 0,242 | 0,652 |
| B9J08_002180 | *SRD1* | | FALSE | -0,479 | 0,212 | 1,406 | 0,026 | 2,329 | 0,019 | 0,445 | 0,286 |
| B9J08_000142 | *RPS26A* | Translation / ribosome biogenesis | TRUE | 1,149 | 0,227 | 1,394 | 0,044 | -0,026 | 0,953 | -0,271 | 0,526 |
| B9J08_002485 | *LEM3* | | FALSE | -0,011 | 0,985 | 1,389 | 0,025 | 1,604 | 0,013 | 0,204 | 0,743 |
| B9J08_002796 | | | FALSE | 0,585 | 0,215 | 1,389 | 0,032 | 1,534 | 0,043 | 0,730 | 0,268 |
| B9J08_004574 | | Translation / ribosome biogenesis | TRUE | 0,503 | 0,204 | 1,385 | 0,030 | 0,137 | 0,526 | -0,745 | 0,211 |
| B9J08_004337 | *TIP20* | Vesicle-mediated transport | TRUE | -0,076 | 0,891 | 1,380 | 0,039 | 1,884 | 0,022 | 0,429 | 0,279 |
| B9J08_004971 | | | FALSE | 4,779 | 0,096 | 1,379 | 0,025 | -2,314 | 0,070 | 1,086 | 0,283 |
| B9J08_004687 | *SEC26* | Vesicle-mediated transport | TRUE | -0,325 | 0,573 | 1,373 | 0,029 | 1,497 | 0,021 | -0,200 | 0,758 |
| B9J08_003539 | | | FALSE | 0,490 | 0,345 | 1,371 | 0,027 | 1,120 | 0,057 | 0,238 | 0,680 |
| B9J08_000489 | | | FALSE | -0,990 | 0,348 | 1,366 | 0,023 | 2,078 | 0,059 | -0,278 | 0,557 |
| B9J08_004818 | | | FALSE | 0,871 | 0,074 | 1,364 | 0,025 | 0,936 | 0,012 | 0,444 | 0,355 |
| B9J08_003890 | | | FALSE | 1,424 | 0,134 | 1,360 | 0,042 | -0,090 | 0,801 | -0,025 | 0,978 |
| B9J08_003557 | *SNL1* | | FALSE | 1,086 | 0,274 | 1,354 | 0,012 | 0,679 | 0,194 | 0,411 | 0,399 |
| B9J08_001107 | *SEF1* | Response to stress | TRUE | 0,356 | 0,669 | 1,349 | 0,029 | 1,618 | 0,035 | 0,625 | 0,468 |
| B9J08_002776 | | | FALSE | -0,151 | 0,313 | 1,320 | 0,049 | 1,210 | 0,012 | -0,261 | 0,569 |
| B9J08_001149 | | | FALSE | 0,844 | 0,131 | 1,312 | 0,042 | 1,299 | 0,040 | 0,831 | 0,258 |
| B9J08_002886 | *GIT3* | Transmembrane transport | TRUE | 0,345 | 0,642 | 1,306 | 0,049 | 2,078 | 0,043 | 1,118 | 0,299 |
| B9J08_003016 | | | FALSE | 1,136 | 0,130 | 1,302 | 0,017 | 0,382 | 0,076 | 0,216 | 0,716 |
| B9J08_003720 | | Transmembrane transport | TRUE | 0,354 | 0,391 | 1,302 | 0,018 | 1,481 | 0,013 | 0,533 | 0,355 |
| B9J08_005281 | | | FALSE | 0,456 | 0,658 | 1,294 | 0,048 | 0,216 | 0,752 | -0,623 | 0,575 |
| B9J08_003281 | *RIM15* | Response to stress; DNA replication / cell cycle | TRUE | 0,308 | 0,280 | 1,290 | 0,031 | 1,610 | 0,031 | 0,628 | 0,235 |
| B9J08_003876 | *RRP6* | | FALSE | 0,538 | 0,314 | 1,284 | 0,010 | 1,269 | 0,026 | 0,523 | 0,436 |
| B9J08_003616 | *OPY2* | Cell wall organization / biogenesis | TRUE | 2,182 | 0,126 | 1,275 | 0,037 | -0,330 | 0,380 | 0,576 | 0,461 |
| B9J08_002673 | | Protein catabolism | TRUE | 1,100 | 0,132 | 1,265 | 0,038 | 0,348 | 0,174 | 0,183 | 0,806 |
| B9J08_001256 | *RPB1* | Response to stress | TRUE | 0,450 | 0,179 | 1,260 | 0,008 | 1,055 | 0,019 | 0,245 | 0,450 |
| B9J08_002252 | | | FALSE | 0,028 | 0,943 | 1,244 | 0,049 | 0,741 | 0,101 | -0,476 | 0,343 |
| B9J08_000431 | *VPS1* | | FALSE | 0,797 | 0,061 | 1,242 | 0,008 | 0,721 | 0,025 | 0,276 | 0,410 |
| B9J08_002867 | | | FALSE | 0,916 | 0,196 | 1,221 | 0,019 | 0,374 | 0,265 | 0,069 | 0,848 |
| B9J08_003715 | | | FALSE | 0,743 | 0,124 | 1,218 | 0,039 | 0,363 | 0,154 | -0,112 | 0,674 |
| B9J08_003224 | *HMG1* | Lipid metabolism | TRUE | 1,441 | 0,075 | 1,210 | 0,013 | -0,121 | 0,496 | 0,110 | 0,881 |
| B9J08_004494 | | Transmembrane transport | TRUE | 0,558 | 0,706 | 1,203 | 0,029 | 1,719 | 0,094 | 1,074 | 0,274 |
| B9J08_005178 | | | FALSE | 0,543 | 0,256 | 1,198 | 0,017 | 0,543 | 0,110 | -0,112 | 0,788 |
| B9J08_000932 | *CDC50* | | FALSE | 0,165 | 0,685 | 1,194 | 0,037 | 0,804 | 0,041 | -0,226 | 0,699 |
| B9J08_000739 | *TIM54* | | FALSE | 0,747 | 0,147 | 1,179 | 0,013 | 0,652 | 0,056 | 0,221 | 0,286 |
| B9J08_003073 | | | FALSE | 4,355 | 0,056 | 1,177 | 0,028 | -2,523 | 0,028 | 0,655 | 0,367 |

| **Gene** | **Symbol** | **All processes** | **Included in Fig. 3e** | **wt-CAS vs wt-ND log2(FC)** | **wt-CAS vs wt-ND p(adj)** | ***hog1*Δ-CAS vs hog1Δ-ND log2(FC)** | ***hog1*Δ-CAS vs hog1Δ-ND p(adj)** | ***hog1*Δ-CAS vs wt-CAS log2(FC)** | ***hog1*Δ-CAS vs wt-CAS p(adj)** | ***hog1*Δ-ND vs wt-ND log2(FC)** | ***hog1*Δ-ND vs wt-ND p(adj)** |
|---|---|---|---|---|---|---|---|---|---|---|---|
| B9J08_000462 | *GLC7* | Response to stress; DNA replication / cell cycle | TRUE | -0,628 | 0,098 | 1,176 | 0,048 | 0,908 | 0,040 | -0,896 | 0,158 |
| B9J08_001142 | *LPT1* | Lipid metabolism | TRUE | -Inf | 0,261 | 1,174 | 0,028 | Inf | 0,010 | -0,177 | 0,878 |
| B9J08_005392 | | | FALSE | -0,315 | 0,501 | 1,171 | 0,025 | 1,174 | 0,019 | -0,312 | 0,607 |
| B9J08_003934 | | Translation / ribosome biogenesis | TRUE | -0,055 | 0,903 | 1,171 | 0,025 | 0,362 | 0,331 | -0,864 | 0,320 |
| B9J08_003931 | | | FALSE | 0,434 | 0,584 | 1,163 | 0,020 | 0,894 | 0,169 | 0,165 | 0,624 |
| B9J08_005274 | | | FALSE | 0,422 | 0,476 | 1,149 | 0,048 | 1,283 | 0,061 | 0,555 | 0,462 |
| B9J08_002532 | | | FALSE | 0,869 | 0,125 | 1,147 | 0,046 | 0,127 | 0,564 | -0,151 | 0,780 |
| B9J08_002615 | *BMS1* | Translation / ribosome biogenesis | TRUE | 1,749 | 0,056 | 1,142 | 0,036 | 0,096 | 0,577 | 0,702 | 0,398 |
| B9J08_000848 | *PEX13* | Lipid metabolism; Transmembrane transport | TRUE | 0,196 | 0,371 | 1,129 | 0,027 | 1,539 | 0,022 | 0,606 | 0,355 |
| B9J08_002133 | | | FALSE | 0,311 | 0,560 | 1,129 | 0,037 | 0,906 | 0,062 | 0,089 | 0,877 |
| B9J08_001924 | *SEC27* | Vesicle-mediated transport | TRUE | -0,336 | 0,269 | 1,128 | 0,017 | 1,279 | 0,022 | -0,185 | 0,513 |
| B9J08_003166 | | | FALSE | -0,239 | 0,618 | 1,124 | 0,017 | 1,555 | 0,013 | 0,192 | 0,702 |
| B9J08_004206 | | | FALSE | 0,064 | 0,763 | 1,122 | 0,025 | 0,542 | 0,065 | -0,516 | 0,229 |
| B9J08_005452 | *MODF* | | FALSE | -0,247 | 0,296 | 1,121 | 0,034 | 0,858 | 0,041 | -0,509 | 0,254 |
| B9J08_003667 | *YPT1* | Vesicle-mediated transport | TRUE | -0,098 | 0,815 | 1,119 | 0,031 | 1,489 | 0,012 | 0,271 | 0,574 |
| B9J08_001037 | *TOM22* | | FALSE | 0,369 | 0,214 | 1,108 | 0,013 | 0,858 | 0,022 | 0,119 | 0,627 |
| B9J08_001972 | *INO80* | | FALSE | 0,257 | 0,557 | 1,092 | 0,034 | 1,045 | 0,014 | 0,210 | 0,677 |
| B9J08_002155 | | | FALSE | 0,551 | 0,098 | 1,090 | 0,028 | -0,251 | 0,096 | -0,791 | 0,274 |
| B9J08_000775 | *TPS2* | Carbohydrate metabolism; Response to stress | TRUE | -0,363 | 0,200 | 1,086 | 0,037 | 1,484 | 0,040 | 0,035 | 0,878 |
| B9J08_004178 | *SIP3* | | FALSE | -0,077 | 0,876 | 1,083 | 0,040 | 1,911 | 0,034 | 0,751 | 0,323 |
| B9J08_005214 | | | FALSE | 0,035 | 0,850 | 1,073 | 0,029 | 1,230 | 0,038 | 0,193 | 0,498 |
| B9J08_002214 | *CGR1* | | FALSE | 3,110 | 0,102 | 1,070 | 0,017 | -2,144 | 0,074 | -0,104 | 0,665 |
| B9J08_002811 | | | FALSE | 0,133 | 0,776 | 1,069 | 0,042 | 0,712 | 0,071 | -0,223 | 0,699 |
| B9J08_004909 | | | FALSE | 1,138 | 0,121 | 1,068 | 0,011 | -0,120 | 0,346 | -0,050 | 0,950 |
| B9J08_002173 | *ENA2* | Transmembrane transport | TRUE | -0,135 | 0,750 | 1,057 | 0,036 | 2,232 | 0,026 | 1,040 | 0,254 |
| B9J08_002476 | *GZF3* | Response to stress | TRUE | 0,385 | 0,225 | 1,056 | 0,040 | 0,978 | 0,026 | 0,307 | 0,254 |
| B9J08_004653 | *ARP2* | Cell wall organization / biogenesis; Vesicle-mediated transport; Cytoskeleton organization; Response to stress | TRUE | 0,434 | 0,122 | 1,045 | 0,038 | 0,638 | 0,025 | 0,027 | 0,949 |
| B9J08_005426 | *UTP18* | Translation / ribosome biogenesis | TRUE | 0,905 | 0,034 | 1,045 | 0,013 | 0,196 | 0,079 | 0,057 | 0,805 |
| B9J08_004501 | *ALO1* | Response to stress | TRUE | 2,001 | 0,089 | 1,039 | 0,018 | -0,015 | 0,949 | 0,947 | 0,226 |
| B9J08_000105 | *DCP2* | Response to stress | TRUE | 0,019 | 0,923 | 1,039 | 0,045 | 1,220 | 0,025 | 0,200 | 0,436 |
| B9J08_001446 | *SAM51* | | FALSE | 0,096 | 0,557 | 1,030 | 0,013 | 0,761 | 0,025 | -0,173 | 0,411 |
| B9J08_005023 | *TIM44* | | FALSE | 0,709 | 0,127 | 1,012 | 0,008 | 0,322 | 0,196 | 0,018 | 0,961 |
| B9J08_005213 | *MYO5* | Cell wall organization / biogenesis; Vesicle-mediated transport; Cytoskeleton organization; Response to stress; DNA replication / cell cycle | TRUE | 0,699 | 0,168 | 1,009 | 0,025 | 0,506 | 0,050 | 0,195 | 0,682 |
| B9J08_001930 | *AOX2* | Cellular respiration | TRUE | -0,631 | 0,195 | 1,009 | 0,017 | 2,209 | 0,015 | 0,569 | 0,267 |
| B9J08_000934 | *NOG2* | | FALSE | 0,948 | 0,152 | 1,004 | 0,025 | 0,526 | 0,051 | 0,470 | 0,472 |
| B9J08_000863 | *UGA1* | | FALSE | -1,206 | 0,083 | -1,005 | 0,031 | 0,118 | 0,089 | -0,083 | 0,632 |
| B9J08_003056 | | | FALSE | 0,080 | 0,786 | -1,007 | 0,031 | -1,232 | 0,045 | -0,144 | 0,668 |
| B9J08_001999 | | Translation / ribosome biogenesis | TRUE | -0,616 | 0,051 | -1,020 | 0,049 | -0,607 | 0,028 | -0,203 | 0,419 |

| Gene | Symbol | All processes | Included in Fig. 3e | wt-CAS vs wt-ND log2(FC) | wt-CAS vs wt-ND p(adj) | ***hog1*Δ-CAS vs hog1Δ-ND log2(FC)** | ***hog1*Δ-CAS vs hog1Δ-ND p(adj)** | ***hog1*Δ-CAS vs wt-CAS log2(FC)** | ***hog1*Δ-CAS vs wt-CAS p(adj)** | ***hog1*Δ-ND vs wt-ND log2(FC)** | ***hog1*Δ-ND vs wt-ND p(adj)** |
|---|---|---|---|---|---|---|---|---|---|---|---|
| B9J08_001695 | *TPS1* | Carbohydrate metabolism; Response to stress; DNA replication / cell cycle | TRUE | -0,428 | 0,255 | -1,020 | 0,042 | -0,535 | 0,248 | 0,057 | 0,763 |
| B9J08_001238 | | | FALSE | 0,835 | 0,170 | -1,022 | 0,022 | -1,182 | 0,089 | 0,676 | 0,179 |
| B9J08_002502 | *MMS2* | | FALSE | -0,144 | 0,228 | -1,026 | 0,027 | -0,652 | 0,040 | 0,230 | 0,257 |
| B9J08_005456 | *GCN3* | Translation / ribosome biogenesis | TRUE | 0,554 | 0,435 | -1,029 | 0,027 | -1,584 | 0,172 | -0,001 | 0,994 |
| B9J08_002232 | *ATM1* | Transmembrane transport | TRUE | -1,036 | 0,317 | -1,039 | 0,025 | 0,000 | 0,999 | 0,004 | 0,995 |
| B9J08_002962 | | | FALSE | -2,105 | 0,103 | -1,040 | 0,008 | 0,480 | 0,047 | -0,585 | 0,350 |
| B9J08_002542 | *SIP5* | | FALSE | 0,797 | 0,034 | -1,043 | 0,025 | -1,374 | 0,011 | 0,466 | 0,235 |
| B9J08_004483 | | | FALSE | -0,510 | 0,210 | -1,047 | 0,030 | -0,110 | 0,757 | 0,428 | 0,252 |
| B9J08_004854 | | | FALSE | -1,325 | 0,322 | -1,049 | 0,041 | 0,150 | 0,919 | -0,127 | 0,665 |
| B9J08_004800 | | | FALSE | 0,192 | 0,671 | -1,052 | 0,043 | -1,434 | 0,142 | -0,191 | 0,479 |
| B9J08_002568 | *PRE6* | Protein catabolism | TRUE | -0,430 | 0,421 | -1,059 | 0,029 | -0,893 | 0,183 | -0,264 | 0,627 |
| B9J08_005449 | *LSM5* | | FALSE | -0,252 | 0,472 | -1,062 | 0,030 | -0,350 | 0,373 | 0,460 | 0,252 |
| B9J08_004720 | | | FALSE | -0,869 | 0,455 | -1,063 | 0,029 | 0,255 | 0,811 | 0,449 | 0,495 |
| B9J08_003939 | | | FALSE | -0,269 | 0,391 | -1,069 | 0,018 | -0,513 | 0,198 | 0,287 | 0,320 |
| B9J08_001740 | *POX1-3* | Lipid metabolism | TRUE | -0,283 | 0,196 | -1,075 | 0,030 | -1,086 | 0,057 | -0,294 | 0,244 |
| B9J08_002303 | *ARG5,6* | | FALSE | -0,298 | 0,482 | -1,082 | 0,031 | -1,413 | 0,124 | -0,630 | 0,320 |
| B9J08_002043 | *ARO1* | | FALSE | -0,207 | 0,493 | -1,085 | 0,029 | -0,704 | 0,051 | 0,173 | 0,598 |
| B9J08_002273 | *PHB1* | | FALSE | 0,152 | 0,339 | -1,092 | 0,049 | -1,027 | 0,066 | 0,217 | 0,459 |
| B9J08_001733 | | | FALSE | -0,576 | 0,232 | -1,095 | 0,029 | -0,922 | 0,201 | -0,402 | 0,320 |
| B9J08_002992 | | | FALSE | -1,553 | 0,183 | -1,099 | 0,042 | -0,192 | 0,565 | -0,646 | 0,447 |
| B9J08_002878 | | | FALSE | -0,534 | 0,223 | -1,100 | 0,036 | -0,715 | 0,191 | -0,149 | 0,307 |
| B9J08_001631 | *CDC37* | Protein folding | TRUE | -0,314 | 0,446 | -1,102 | 0,023 | -0,886 | 0,158 | -0,098 | 0,759 |
| B9J08_003246 | | | FALSE | -0,849 | 0,227 | -1,103 | 0,038 | -0,772 | 0,308 | -0,519 | 0,454 |
| B9J08_003986 | | | FALSE | 0,102 | 0,753 | -1,111 | 0,045 | -1,699 | 0,088 | -0,486 | 0,250 |
| B9J08_001350 | | | FALSE | -3,869 | 0,112 | -1,114 | 0,019 | 2,940 | 0,046 | 0,186 | 0,694 |
| B9J08_005134 | *SOL3* | Carbohydrate metabolism | TRUE | -0,297 | 0,149 | -1,119 | 0,030 | -0,828 | 0,051 | -0,006 | 0,970 |
| B9J08_001646 | | | FALSE | 0,103 | 0,414 | -1,122 | 0,031 | -1,606 | 0,006 | -0,381 | 0,249 |
| B9J08_000076 | *SCP1* | Cytoskeleton organization; DNA replication / cell cycle | TRUE | 0,027 | 0,949 | -1,127 | 0,041 | -0,534 | 0,263 | 0,621 | 0,274 |
| B9J08_001271 | *CCT8* | Response to stress | TRUE | -0,011 | 0,902 | -1,130 | 0,012 | -1,094 | 0,024 | 0,025 | 0,746 |
| B9J08_002982 | *KTI12* | | FALSE | -0,430 | 0,233 | -1,131 | 0,042 | -0,831 | 0,112 | -0,130 | 0,585 |
| B9J08_003645 | *POB3* | Response to stress | TRUE | 0,314 | 0,212 | -1,135 | 0,023 | -1,486 | 0,042 | -0,037 | 0,863 |
| B9J08_002611 | *KIN2* | Cytoskeleton organization; DNA replication / cell cycle | TRUE | 0,367 | 0,377 | -1,144 | 0,013 | -1,045 | 0,063 | 0,466 | 0,406 |
| B9J08_004694 | | | FALSE | 0,230 | 0,432 | -1,149 | 0,008 | -0,856 | 0,106 | 0,523 | 0,158 |
| B9J08_001771 | | | FALSE | -0,082 | 0,902 | -1,154 | 0,038 | -0,948 | 0,341 | 0,124 | 0,605 |
| B9J08_003357 | *RPN13* | | FALSE | 0,111 | 0,650 | -1,178 | 0,028 | -1,587 | 0,010 | -0,297 | 0,469 |
| B9J08_001730 | *SEN1* | | FALSE | 0,891 | 0,321 | -1,189 | 0,027 | -1,758 | 0,158 | 0,322 | 0,268 |
| B9J08_005227 | | | FALSE | -1,321 | 0,096 | -1,192 | 0,013 | -0,318 | 0,147 | -0,447 | 0,324 |
| B9J08_005145 | | | FALSE | 0,016 | 0,947 | -1,198 | 0,028 | -1,196 | 0,017 | 0,018 | 0,957 |
| B9J08_002112 | *TIM9* | | FALSE | 0,360 | 0,194 | -1,201 | 0,030 | -1,394 | 0,010 | 0,167 | 0,560 |
| B9J08_004763 | *MET8* | | FALSE | -0,231 | 0,440 | -1,204 | 0,041 | -0,429 | 0,068 | 0,544 | 0,266 |
| B9J08_003608 | | | FALSE | 0,268 | 0,389 | -1,222 | 0,008 | -1,066 | 0,088 | 0,424 | 0,258 |
| B9J08_001679 | *CPH2* | Response to stress | TRUE | -1,094 | 0,221 | -1,238 | 0,008 | 0,399 | 0,315 | 0,542 | 0,405 |
| B9J08_001919 | | | FALSE | -0,114 | 0,668 | -1,238 | 0,031 | -1,485 | 0,060 | -0,361 | 0,301 |

| Gene | Symbol | All processes | Included in Fig. 3e | wt-CAS vs wt-ND log2(FC) | wt-CAS vs wt-ND p(adj) | *hog1*Δ-CAS vs hog1Δ-ND log2(FC) | *hog1*Δ-CAS vs hog1Δ-ND p(adj) | *hog1*Δ-CAS vs wt-CAS log2(FC) | *hog1*Δ-CAS vs wt-CAS p(adj) | *hog1*Δ-ND vs wt-ND log2(FC) | *hog1*Δ-ND vs wt-ND p(adj) |
|---|---|---|---|---|---|---|---|---|---|---|---|
| B9J08_002528 | | | FALSE | 0,320 | 0,557 | -1,239 | 0,029 | -1,261 | 0,172 | 0,298 | 0,286 |
| B9J08_002845 | | Response to stress; DNA replication / cell cycle | TRUE | 0,691 | 0,133 | -1,241 | 0,029 | -2,031 | 0,032 | -0,100 | 0,608 |
| B9J08_004583 | | | FALSE | -0,441 | 0,093 | -1,243 | 0,033 | -0,871 | 0,057 | -0,069 | 0,627 |
| B9J08_002315 | | | FALSE | -1,777 | 0,246 | -1,245 | 0,023 | 0,780 | 0,232 | 0,248 | 0,720 |
| B9J08_002762 | *INO1* | Lipid metabolism | TRUE | -1,575 | 0,252 | -1,276 | 0,025 | 0,390 | 0,302 | 0,090 | 0,895 |
| B9J08_002137 | | DNA replication / cell cycle | TRUE | 0,513 | 0,256 | -1,279 | 0,037 | -1,236 | 0,062 | 0,557 | 0,346 |
| B9J08_000916 | | | FALSE | 0,130 | 0,750 | -1,285 | 0,009 | -1,027 | 0,028 | 0,388 | 0,475 |
| B9J08_000204 | | Vesicle-mediated transport | TRUE | -0,359 | 0,205 | -1,293 | 0,013 | -0,642 | 0,085 | 0,292 | 0,270 |
| B9J08_003811 | | | FALSE | 0,331 | 0,480 | -1,305 | 0,027 | -1,355 | 0,128 | 0,281 | 0,324 |
| B9J08_001183 | *ZRC1* | Response to stress; Transmembrane transport | TRUE | 0,248 | 0,557 | -1,311 | 0,029 | -1,189 | 0,096 | 0,370 | 0,458 |
| B9J08_003285 | *ACB1* | | FALSE | 0,276 | 0,592 | -1,323 | 0,018 | -1,844 | 0,068 | -0,245 | 0,726 |
| B9J08_000396 | | | FALSE | -0,978 | 0,096 | -1,338 | 0,013 | -0,146 | 0,406 | 0,214 | 0,454 |
| B9J08_002527 | | | FALSE | 0,047 | 0,921 | -1,342 | 0,040 | -1,025 | 0,205 | 0,365 | 0,286 |
| B9J08_001759 | *SCS7* | Lipid metabolism | TRUE | 1,432 | 0,112 | -1,342 | 0,031 | -1,828 | 0,071 | 0,946 | 0,268 |
| B9J08_005093 | | | FALSE | -0,931 | 0,147 | -1,344 | 0,029 | -0,433 | 0,301 | -0,020 | 0,960 |
| B9J08_001493 | *AKL1* | | FALSE | 0,423 | 0,109 | -1,348 | 0,021 | -1,383 | 0,019 | 0,388 | 0,249 |
| B9J08_001963 | *PEX14* | Transmembrane transport | TRUE | -0,206 | 0,470 | -1,348 | 0,030 | -0,964 | 0,087 | 0,178 | 0,422 |
| B9J08_000618 | | | FALSE | 0,494 | 0,426 | -1,355 | 0,042 | -1,158 | 0,163 | 0,690 | 0,263 |
| B9J08_002146 | *YCF1* | Response to stress; Transmembrane transport | TRUE | -0,102 | 0,807 | -1,359 | 0,025 | -0,972 | 0,022 | 0,285 | 0,585 |
| B9J08_001842 | *RPT4* | | FALSE | 0,064 | 0,286 | -1,366 | 0,036 | -1,360 | 0,014 | 0,070 | 0,621 |
| B9J08_000038 | | Translation / ribosome biogenesis | TRUE | 0,742 | 0,635 | -1,369 | 0,031 | -1,984 | 0,388 | 0,127 | 0,854 |
| B9J08_005298 | *HAP43* | | FALSE | -0,240 | 0,708 | -1,386 | 0,034 | 0,059 | 0,923 | 1,206 | 0,143 |
| B9J08_003236 | | | FALSE | -0,404 | 0,325 | -1,390 | 0,019 | -0,542 | 0,130 | 0,443 | 0,326 |
| B9J08_003721 | | Response to stress | TRUE | 0,521 | 0,496 | -1,394 | 0,049 | -1,261 | 0,227 | 0,655 | 0,300 |
| B9J08_002792 | *RCK2* | Cell wall organization / biogenesis; Response to stress; DNA replication / cell cycle | TRUE | 0,966 | 0,147 | -1,409 | 0,027 | -2,921 | 0,059 | -0,547 | 0,196 |
| B9J08_001502 | | | FALSE | 0,723 | 0,089 | -1,418 | 0,017 | -1,712 | 0,010 | 0,429 | 0,293 |
| B9J08_001590 | | | FALSE | -3,010 | 0,247 | -1,433 | 0,045 | 1,289 | 0,209 | -0,289 | 0,763 |
| B9J08_005383 | *FMP27* | | FALSE | -0,403 | 0,521 | -1,443 | 0,043 | -0,513 | 0,069 | 0,527 | 0,432 |
| B9J08_001081 | *RPL23A* | Translation / ribosome biogenesis | TRUE | 0,377 | 0,431 | -1,444 | 0,013 | -1,620 | 0,037 | 0,201 | 0,707 |
| B9J08_002009 | | | FALSE | 1,136 | 0,084 | -1,458 | 0,020 | -1,836 | 0,057 | 0,758 | 0,240 |
| B9J08_003886 | *SLP2* | Lipid metabolism | TRUE | 0,090 | 0,615 | -1,470 | 0,012 | -2,002 | 0,028 | -0,443 | 0,308 |
| B9J08_004072 | *QCR2* | | FALSE | -1,265 | 0,075 | -1,480 | 0,033 | -0,679 | 0,176 | -0,464 | 0,307 |
| B9J08_000253 | | | FALSE | -0,752 | 0,073 | -1,488 | 0,030 | -0,788 | 0,018 | -0,051 | 0,763 |
| B9J08_003725 | *TIF33* | | FALSE | -0,077 | 0,840 | -1,494 | 0,030 | -1,457 | 0,114 | -0,040 | 0,903 |
| B9J08_001806 | | | FALSE | -0,079 | 0,786 | -1,509 | 0,036 | -1,194 | 0,042 | 0,236 | 0,510 |
| B9J08_002760 | | | FALSE | -1,902 | 0,204 | -1,523 | 0,035 | 0,253 | 0,887 | -0,126 | 0,739 |
| B9J08_005220 | | Translation / ribosome biogenesis | TRUE | -0,043 | 0,927 | -1,524 | 0,013 | -1,249 | 0,087 | 0,232 | 0,627 |
| B9J08_000365 | *SGA1* | Carbohydrate metabolism | TRUE | -1,657 | 0,182 | -1,524 | 0,048 | -0,100 | 0,946 | -0,232 | 0,657 |
| B9J08_005050 | | | FALSE | -0,366 | 0,334 | -1,539 | 0,030 | -0,987 | 0,100 | 0,186 | 0,577 |
| B9J08_005119 | *BRF1* | | FALSE | -0,167 | 0,660 | -1,546 | 0,027 | -1,083 | 0,052 | 0,296 | 0,514 |
| B9J08_000299 | *FZO1* | | FALSE | 0,189 | 0,416 | -1,562 | 0,025 | -1,682 | 0,040 | 0,069 | 0,763 |
| B9J08_002181 | *FAB1* | Lipid metabolism; Response to stress | TRUE | 0,642 | 0,212 | -1,572 | 0,027 | -1,540 | 0,060 | 0,673 | 0,270 |

| Gene | Symbol | All processes | Included in Fig. 3e | wt-CAS vs wt-ND log2(FC) | wt-CAS vs wt-ND p(adj) | *hog1*Δ-CAS vs hog1Δ-ND log2(FC) | *hog1*Δ-CAS vs hog1Δ-ND p(adj) | *hog1*Δ-CAS vs wt-CAS log2(FC) | *hog1*Δ-CAS vs wt-CAS p(adj) | *hog1*Δ-ND vs wt-ND log2(FC) | *hog1*Δ-ND vs wt-ND p(adj) |
|---|---|---|---|---|---|---|---|---|---|---|---|
| B9J08_005275 | | | FALSE | -0,838 | 0,061 | -1,573 | 0,039 | -0,448 | 0,053 | 0,287 | 0,338 |
| B9J08_000498 | *IPP1* | | FALSE | -1,080 | 0,309 | -1,576 | 0,048 | -1,134 | 0,296 | -0,639 | 0,525 |
| B9J08_001297 | *VIP1* | Cytoskeleton organization; DNA replication / cell cycle | TRUE | 0,352 | 0,162 | -1,576 | 0,030 | -1,423 | 0,037 | 0,505 | 0,235 |
| B9J08_005144 | *HTA2* | Response to stress; DNA replication / cell cycle | TRUE | 0,220 | 0,326 | -1,583 | 0,012 | -1,710 | 0,029 | 0,094 | 0,627 |
| B9J08_001118 | *CIC1* | | FALSE | 1,106 | 0,241 | -1,593 | 0,046 | -2,426 | 0,125 | 0,273 | 0,550 |
| B9J08_000234 | *RFX1* | | FALSE | 1,168 | 0,086 | -1,616 | 0,030 | -2,265 | 0,037 | 0,519 | 0,268 |
| B9J08_004953 | *TSR2* | | FALSE | -0,036 | 0,857 | -1,616 | 0,018 | -1,791 | 0,050 | -0,211 | 0,313 |
| B9J08_004272 | | | FALSE | 0,907 | 0,196 | -1,633 | 0,008 | -1,780 | 0,104 | 0,760 | 0,266 |
| B9J08_005110 | | | FALSE | 0,326 | 0,730 | -1,641 | 0,025 | -1,531 | 0,311 | 0,436 | 0,347 |
| B9J08_000099 | | | FALSE | 0,379 | 0,440 | -1,641 | 0,008 | -1,996 | 0,106 | 0,024 | 0,854 |
| B9J08_002072 | | Response to stress; DNA replication / cell cycle | TRUE | 0,516 | 0,183 | -1,651 | 0,031 | -2,400 | 0,046 | -0,233 | 0,590 |
| B9J08_003550 | *YWP1* | | FALSE | -0,576 | 0,367 | -1,695 | 0,026 | -2,043 | 0,101 | -0,924 | 0,393 |
| B9J08_001867 | | | FALSE | -0,020 | 0,970 | -1,696 | 0,034 | -1,362 | 0,163 | 0,314 | 0,466 |
| B9J08_005087 | | | FALSE | 0,169 | 0,759 | -1,703 | 0,024 | -1,640 | 0,029 | 0,232 | 0,722 |
| B9J08_003251 | *XOG1* | Cell wall organization / biogenesis; Carbohydrate metabolism | TRUE | 0,441 | 0,092 | -1,739 | 0,029 | -3,771 | 0,019 | -1,592 | 0,211 |
| B9J08_000654 | *SVF1* | Response to stress | TRUE | -1,467 | 0,250 | -1,747 | 0,022 | -0,907 | 0,609 | -0,627 | 0,478 |
| B9J08_003394 | *MMD1* | | FALSE | -1,047 | 0,275 | -1,764 | 0,008 | -0,884 | 0,496 | -0,167 | 0,254 |
| B9J08_005510 | *SNU13* | Translation / ribosome biogenesis | TRUE | -1,217 | 0,449 | -1,776 | 0,049 | -1,110 | 0,658 | -0,550 | 0,627 |
| B9J08_005245 | *PGA4* | Cell wall organization / biogenesis; Carbohydrate metabolism; DNA replication / cell cycle | TRUE | -0,412 | 0,671 | -1,781 | 0,034 | -1,401 | 0,426 | -0,033 | 0,933 |
| B9J08_003089 | *RPS24* | Translation / ribosome biogenesis | TRUE | 1,370 | 0,122 | -1,782 | 0,035 | -2,288 | 0,068 | 0,864 | 0,185 |
| B9J08_004505 | | | FALSE | 0,081 | 0,774 | -1,782 | 0,021 | -1,755 | 0,029 | 0,108 | 0,731 |
| B9J08_003075 | *NCB2* | | FALSE | 0,309 | 0,523 | -1,783 | 0,028 | -1,615 | 0,095 | 0,476 | 0,307 |
| B9J08_002195 | *DPB4* | Response to stress | TRUE | 0,275 | 0,493 | -1,786 | 0,019 | -1,768 | 0,074 | 0,293 | 0,375 |
| B9J08_003511 | | | FALSE | 0,443 | 0,629 | -1,794 | 0,038 | -2,209 | 0,245 | 0,028 | 0,911 |
| B9J08_001169 | | | FALSE | -0,550 | 0,286 | -1,827 | 0,013 | -1,276 | 0,174 | 0,001 | 0,995 |
| B9J08_003438 | | | FALSE | 0,113 | 0,762 | -1,842 | 0,012 | -1,736 | 0,107 | 0,220 | 0,373 |
| B9J08_004868 | *LIG1* | | FALSE | 0,397 | 0,377 | -1,853 | 0,029 | -1,791 | 0,059 | 0,459 | 0,346 |
| B9J08_001645 | | | FALSE | -0,674 | 0,534 | -1,856 | 0,041 | -1,199 | 0,508 | -0,017 | 0,967 |
| B9J08_001800 | | Response to stress | TRUE | 0,298 | 0,328 | -1,862 | 0,029 | -2,017 | 0,047 | 0,143 | 0,627 |
| B9J08_004068 | *VPS4* | Vesicle-mediated transport; Protein catabolism; DNA replication / cell cycle | TRUE | 0,105 | 0,476 | -1,870 | 0,025 | -1,630 | 0,037 | 0,345 | 0,235 |
| B9J08_001227 | *TDH3* | Carbohydrate metabolism; Cellular respiration; Response to stress | TRUE | -0,065 | 0,705 | -1,892 | 0,002 | -2,233 | 0,040 | -0,406 | 0,274 |
| B9J08_004766 | *DRS2* | | FALSE | 0,299 | 0,552 | -1,913 | 0,029 | -2,039 | 0,101 | 0,174 | 0,660 |
| B9J08_000882 | | | FALSE | -1,866 | 0,085 | -1,919 | 0,027 | -0,498 | 0,647 | -0,445 | 0,387 |
| B9J08_004565 | *ICE2* | Lipid metabolism | TRUE | -Inf | 0,183 | -1,930 | 0,025 | Inf | 0,143 | -0,451 | 0,627 |
| B9J08_000925 | *ATG13* | Response to stress | TRUE | 2,227 | 0,301 | -1,988 | 0,025 | -3,726 | 0,205 | 0,489 | 0,270 |
| B9J08_002777 | | | FALSE | -1,296 | 0,222 | -2,008 | 0,031 | -0,451 | 0,714 | 0,261 | 0,472 |
| B9J08_005314 | | Carbohydrate metabolism | TRUE | 0,019 | 0,899 | -2,011 | 0,046 | -2,106 | 0,041 | -0,076 | 0,696 |
| B9J08_001382 | | | FALSE | -1,913 | 0,126 | -2,013 | 0,048 | -0,609 | 0,242 | -0,509 | 0,433 |
| B9J08_000518 | | Vesicle-mediated transport | TRUE | -0,106 | 0,584 | -2,051 | 0,013 | -1,911 | 0,025 | 0,033 | 0,846 |

| Gene | Symbol | All processes | Included in Fig. 3e | wt-CAS vs wt-ND log2(FC) | wt-CAS vs wt-ND p(adj) | *hog1*Δ-CAS vs hog1Δ-ND log2(FC) | *hog1*Δ-CAS vs hog1Δ-ND p(adj) | *hog1*Δ-CAS vs wt-CAS log2(FC) | *hog1*Δ-CAS vs wt-CAS p(adj) | *hog1*Δ-ND vs wt-ND log2(FC) | *hog1*Δ-ND vs wt-ND p(adj) |
|---|---|---|---|---|---|---|---|---|---|---|---|
| B9J08_003799 | *MP65* | Carbohydrate metabolism; Response to stress | TRUE | 0,314 | 0,272 | -2,055 | 0,013 | -3,007 | 0,014 | -0,638 | 0,307 |
| B9J08_004018 | | | FALSE | -0,933 | 0,402 | -2,060 | 0,021 | -0,923 | 0,538 | 0,203 | 0,721 |
| B9J08_000850 | | | FALSE | -1,169 | 0,092 | -2,080 | 0,020 | -0,957 | 0,210 | -0,046 | 0,779 |
| B9J08_001379 | | | FALSE | -0,778 | 0,240 | -2,189 | 0,029 | -1,285 | 0,230 | 0,126 | 0,740 |
| B9J08_005336 | *RPL12* | Translation / ribosome biogenesis | TRUE | 0,464 | 0,551 | -2,219 | 0,013 | -2,160 | 0,168 | 0,523 | 0,409 |
| B9J08_002761 | *CHT3* | Cell wall organization / biogenesis; Carbohydrate metabolism; DNA replication / cell cycle | TRUE | -1,948 | 0,085 | -2,231 | 0,017 | -0,268 | 0,512 | 0,014 | 0,964 |
| B9J08_002374 | *HOC1* | | FALSE | -0,045 | 0,970 | -2,236 | 0,012 | -2,112 | 0,380 | 0,079 | 0,857 |
| B9J08_000715 | | | FALSE | -1,789 | 0,215 | -2,239 | 0,037 | -0,667 | 0,749 | -0,217 | 0,524 |
| B9J08_004739 | | | FALSE | -1,779 | 0,291 | -2,255 | 0,017 | -0,229 | 0,924 | 0,246 | 0,330 |
| B9J08_004907 | *PRP8* | | FALSE | 0,369 | 0,594 | -2,255 | 0,016 | -2,285 | 0,118 | 0,339 | 0,627 |
| B9J08_002688 | *FDH1* | | FALSE | -3,987 | 0,142 | -2,268 | 0,012 | 1,169 | 0,157 | -0,550 | 0,472 |
| B9J08_002393 | | | FALSE | -0,118 | 0,877 | -2,286 | 0,021 | -1,477 | 0,106 | 0,691 | 0,443 |
| B9J08_000437 | *UBC13* | Response to stress | TRUE | -0,998 | 0,186 | -2,297 | 0,020 | -0,337 | 0,682 | 0,962 | 0,158 |
| B9J08_004806 | *TFC4* | | FALSE | 0,003 | 0,992 | -2,351 | 0,021 | -1,656 | 0,054 | 0,698 | 0,185 |
| B9J08_002011 | | | FALSE | 0,046 | 0,943 | -2,421 | 0,031 | -1,925 | 0,134 | 0,542 | 0,436 |
| B9J08_001552 | | | FALSE | 0,939 | 0,148 | -2,424 | 0,039 | -3,371 | 0,040 | -0,008 | 0,970 |
| B9J08_001402 | | | FALSE | 0,467 | 0,215 | -2,532 | 0,030 | -3,666 | 0,031 | -0,668 | 0,208 |
| B9J08_004046 | *MCD4* | Lipid metabolism | TRUE | 0,305 | 0,833 | -2,535 | 0,025 | -2,005 | 0,388 | 0,836 | 0,431 |
| B9J08_001841 | *GCD1* | Translation / ribosome biogenesis | TRUE | 0,114 | 0,473 | -2,538 | 0,043 | -2,868 | 0,007 | -0,216 | 0,479 |
| B9J08_004029 | | | FALSE | -0,145 | 0,908 | -2,544 | 0,031 | -1,753 | 0,408 | 0,646 | 0,447 |
| B9J08_000430 | | | FALSE | -1,460 | 0,195 | -2,584 | 0,009 | -0,817 | 0,573 | 0,307 | 0,271 |
| B9J08_005266 | | | FALSE | -0,837 | 0,236 | -2,596 | 0,031 | -1,262 | 0,028 | 0,497 | 0,352 |
| B9J08_003002 | | Transmembrane transport | TRUE | -Inf | 0,198 | -2,675 | 0,048 | Inf | 0,316 | 1,423 | 0,254 |
| B9J08_003740 | *RAD1* | Response to stress; DNA replication / cell cycle | TRUE | 1,066 | 0,157 | -2,723 | 0,025 | -2,828 | 0,055 | 0,961 | 0,235 |
| B9J08_003912 | | | FALSE | -4,711 | 0,070 | -2,805 | 0,022 | 1,775 | 0,427 | -0,131 | 0,683 |
| B9J08_003477 | *PRP40* | | FALSE | 0,835 | 0,058 | -2,893 | 0,029 | -3,364 | 0,006 | 0,364 | 0,320 |
| B9J08_002798 | *OLE2* | Lipid metabolism | TRUE | 0,538 | 0,769 | -2,927 | 0,042 | -2,325 | 0,299 | 1,141 | 0,531 |
| B9J08_004159 | | | FALSE | 0,428 | 0,267 | -2,944 | 0,025 | -2,465 | 0,057 | 0,907 | 0,208 |
| B9J08_002788 | *TPK2* | Cell wall organization / biogenesis; Carbohydrate metabolism; Response to stress | TRUE | 0,212 | 0,806 | -2,967 | 0,025 | -3,027 | 0,203 | 0,151 | 0,790 |
| B9J08_001046 | | Translation / ribosome biogenesis | TRUE | -0,582 | 0,590 | -2,990 | 0,035 | -2,121 | 0,402 | 0,287 | 0,431 |
| B9J08_005546 | | | FALSE | -Inf | 0,225 | -3,108 | 0,027 | Inf | 0,400 | 0,920 | 0,345 |
| B9J08_004873 | | Protein folding | TRUE | 0,002 | 0,979 | -3,142 | 0,031 | -2,768 | 0,010 | 0,376 | 0,286 |
| B9J08_000049 | *KRE9* | Cell wall organization / biogenesis; Carbohydrate metabolism | TRUE | -0,018 | 0,976 | -3,194 | 0,013 | -4,076 | 0,139 | -0,900 | 0,257 |
| B9J08_004804 | *HPD1* | Lipid metabolism | TRUE | -1,480 | 0,095 | -3,341 | 0,030 | -2,183 | 0,230 | -0,323 | 0,421 |
| B9J08_001121 | *MAC1* | Cellular respiration | TRUE | -0,002 | 0,996 | -3,591 | 0,012 | -3,096 | 0,022 | 0,494 | 0,309 |
| B9J08_001704 | | | FALSE | 0,994 | 0,351 | -4,020 | 0,040 | -4,610 | 0,140 | 0,404 | 0,459 |
| B9J08_004617 | *PTR3* | | FALSE | -1,149 | 0,351 | -4,029 | 0,027 | -2,223 | 0,436 | 0,656 | 0,352 |
| B9J08_000282 | *SPP1* | | FALSE | 0,435 | 0,493 | -4,052 | 0,039 | -4,818 | 0,094 | -0,330 | 0,627 |
| B9J08_004295 | *PSD2* | Lipid metabolism; Response to stress | TRUE | 1,488 | 0,124 | -4,077 | 0,034 | -5,163 | 0,065 | 0,403 | 0,390 |
| B9J08_000550 | *SWD3* | | FALSE | 0,085 | 0,929 | -4,237 | 0,018 | -4,248 | 0,239 | 0,074 | 0,822 |

| Gene | Symbol | All processes | Included in Fig. 3e | wt-CAS vs wt-ND log2(FC) | wt-CAS vs wt-ND p(adj) | *hog1*Δ-CAS vs hog1Δ-ND log2(FC) | *hog1*Δ-CAS vs hog1Δ-ND p(adj) | *hog1*Δ-CAS vs wt-CAS log2(FC) | *hog1*Δ-CAS vs wt-CAS p(adj) | *hog1*Δ-ND vs wt-ND log2(FC) | *hog1*Δ-ND vs wt-ND p(adj) |
|---|---|---|---|---|---|---|---|---|---|---|---|
| B9J08_001282 | *VPS75* | | FALSE | -0,819 | 0,073 | -4,289 | 0,038 | -3,907 | 0,051 | -0,437 | 0,246 |
| B9J08_003179 | | | FALSE | 0,142 | 0,954 | -4,578 | 0,013 | -4,220 | 0,533 | 0,499 | 0,274 |
| B9J08_002618 | | Protein catabolism | TRUE | -0,100 | 0,788 | -4,637 | 0,044 | -4,172 | 0,071 | 0,365 | 0,409 |
| B9J08_002120 | *ENP2* | | FALSE | 0,220 | 0,588 | -4,643 | 0,020 | -4,422 | 0,079 | 0,441 | 0,266 |
| B9J08_003948 | *KIS2* | Carbohydrate metabolism | TRUE | -0,904 | 0,405 | -4,671 | 0,032 | -4,101 | 0,326 | -0,335 | 0,535 |
| B9J08_001844 | *NIP100* | Cytoskeleton organization; DNA replication / cell cycle | TRUE | 1,204 | 0,577 | -4,791 | 0,020 | -4,070 | 0,312 | 1,926 | 0,373 |
| B9J08_002364 | *PAM17* | | FALSE | -0,436 | 0,525 | -5,229 | 0,033 | -3,573 | 0,053 | 1,220 | 0,233 |
| B9J08_003736 | *SSN3* | | FALSE | 2,362 | 0,129 | -6,012 | 0,045 | -6,375 | 0,096 | 1,999 | 0,257 |
| B9J08_003065 | | | FALSE | -0,328 | 0,698 | -6,589 | 0,045 | -6,287 | 0,168 | -0,026 | 0,972 |
| B9J08_001540 | | | FALSE | -5,229 | 0,099 | 0,399 | 0,124 | 6,496 | 0,036 | 0,868 | 0,211 |
| B9J08_001450 | *CFL4* | | FALSE | -2,071 | 0,262 | 0,340 | 0,080 | 5,246 | 0,006 | 2,835 | 0,143 |
| B9J08_002110 | *SIT1* | Transmembrane transport | TRUE | -3,690 | 0,084 | 0,778 | 0,070 | 4,555 | 0,019 | 0,087 | 0,822 |
| B9J08_003527 | | Transmembrane transport | TRUE | -2,040 | 0,202 | 2,589 | 0,056 | 4,408 | 0,037 | -0,221 | 0,700 |
| B9J08_004637 | | | FALSE | -3,418 | 0,067 | 1,020 | 0,062 | 4,398 | 0,022 | -0,041 | 0,952 |
| B9J08_002171 | *HPA2* | | FALSE | -4,058 | 0,108 | 0,664 | 0,075 | 4,317 | 0,040 | -0,405 | 0,461 |
| B9J08_005127 | | | FALSE | -3,514 | 0,268 | 0,970 | 0,102 | 4,244 | 0,032 | -0,240 | 0,826 |
| B9J08_005404 | | | FALSE | -4,011 | 0,280 | -0,008 | 0,974 | 4,180 | 0,027 | 0,178 | 0,864 |
| B9J08_003231 | *CTN1* | Cellular respiration | TRUE | -1,744 | 0,123 | 1,831 | 0,059 | 3,900 | 0,049 | 0,325 | 0,459 |
| B9J08_001069 | | | FALSE | -3,529 | 0,084 | 0,308 | 0,172 | 3,882 | 0,018 | 0,045 | 0,922 |
| B9J08_002930 | *ZCF13* | Carbohydrate metabolism; Response to stress | TRUE | -3,515 | 0,253 | -0,120 | 0,660 | 3,880 | 0,025 | 0,485 | 0,608 |
| B9J08_002965 | | Transmembrane transport | TRUE | -3,041 | 0,149 | 1,674 | 0,112 | 3,808 | 0,019 | -0,907 | 0,524 |
| B9J08_005075 | *UCF1* | | FALSE | -2,769 | 0,074 | 0,219 | 0,380 | 3,609 | 0,036 | 0,622 | 0,310 |
| B9J08_001948 | | | FALSE | -0,255 | 0,925 | 2,197 | 0,080 | 3,562 | 0,043 | 1,110 | 0,513 |
| B9J08_001292 | *SOD4* | Response to stress | TRUE | -2,746 | 0,428 | -1,528 | 0,227 | 3,504 | 0,043 | 2,285 | 0,338 |
| B9J08_004458 | *ZCF27* | | FALSE | -2,002 | 0,228 | 0,626 | 0,110 | 3,462 | 0,025 | 0,834 | 0,324 |
| B9J08_000347 | *HGT18* | Transmembrane transport | TRUE | -3,535 | 0,086 | 0,138 | 0,370 | 3,457 | 0,027 | -0,216 | 0,638 |
| B9J08_001285 | | | FALSE | -1,663 | 0,341 | 1,304 | 0,079 | 3,414 | 0,036 | 0,447 | 0,576 |
| B9J08_000240 | | | FALSE | -0,342 | 0,751 | 2,638 | 0,069 | 3,180 | 0,043 | 0,199 | 0,555 |
| B9J08_003316 | | | FALSE | -2,485 | 0,145 | 0,408 | 0,353 | 3,157 | 0,048 | 0,264 | 0,681 |
| B9J08_003391 | | | FALSE | -2,157 | 0,126 | -0,136 | 0,481 | 2,855 | 0,040 | 0,834 | 0,246 |
| B9J08_003570 | | Vesicle-mediated transport | TRUE | -1,942 | 0,166 | 0,541 | 0,119 | 2,625 | 0,032 | 0,143 | 0,793 |
| B9J08_005375 | | Transmembrane transport | TRUE | -0,766 | 0,552 | 0,876 | 0,142 | 2,551 | 0,040 | 0,910 | 0,461 |
| B9J08_005565 | | | FALSE | -1,404 | 0,089 | 0,813 | 0,024 | 2,533 | 0,015 | 0,316 | 0,385 |
| B9J08_000902 | *TEA1* | | FALSE | -0,043 | 0,951 | 1,394 | 0,053 | 2,486 | 0,010 | 1,050 | 0,388 |
| B9J08_000693 | *CRM1* | | FALSE | -0,218 | 0,730 | 2,230 | 0,052 | 2,433 | 0,044 | -0,016 | 0,982 |
| B9J08_004878 | *MEP2* | Response to stress; Transmembrane transport | TRUE | -3,051 | 0,136 | -1,052 | 0,121 | 2,396 | 0,010 | 0,397 | 0,488 |
| B9J08_002534 | *SCT2* | | FALSE | -1,978 | 0,241 | -0,026 | 0,920 | 2,395 | 0,049 | 0,443 | 0,571 |
| B9J08_002108 | *FTR1* | Lipid metabolism; Response to stress | TRUE | -0,492 | 0,594 | -0,016 | 0,948 | 2,379 | 0,050 | 1,903 | 0,185 |
| B9J08_005552 | | | FALSE | -1,624 | 0,115 | 0,917 | 0,038 | 2,360 | 0,024 | -0,181 | 0,657 |
| B9J08_001834 | *TPO4* | Transmembrane transport | TRUE | -1,637 | 0,238 | -0,472 | 0,083 | 2,330 | 0,042 | 1,165 | 0,255 |
| B9J08_000180 | *MET18* | | FALSE | -0,508 | 0,688 | 1,348 | 0,088 | 2,303 | 0,050 | 0,446 | 0,559 |
| B9J08_005351 | *CWH43* | | FALSE | -1,710 | 0,136 | 0,237 | 0,109 | 2,287 | 0,011 | 0,340 | 0,468 |
| B9J08_000494 | *PHO91* | Transmembrane transport | TRUE | -1,308 | 0,125 | 0,969 | 0,066 | 2,284 | 0,042 | 0,007 | 0,986 |

| Gene | Symbol | All processes | Included in Fig. 3e | wt-CAS vs wt-ND log2(FC) | wt-CAS vs wt-ND p(adj) | *hog1*Δ-CAS vs hog1Δ-ND log2(FC) | *hog1*Δ-CAS vs hog1Δ-ND p(adj) | *hog1*Δ-CAS vs wt-CAS log2(FC) | *hog1*Δ-CAS vs wt-CAS p(adj) | *hog1*Δ-ND vs wt-ND log2(FC) | *hog1*Δ-ND vs wt-ND p(adj) |
|---|---|---|---|---|---|---|---|---|---|---|---|
| B9J08_001914 | | | FALSE | -1,993 | 0,173 | 0,397 | 0,025 | 2,248 | 0,008 | -0,143 | 0,787 |
| B9J08_005509 | *CUP5* | Transmembrane transport | TRUE | -1,539 | 0,286 | 0,565 | 0,014 | 2,148 | 0,010 | 0,045 | 0,957 |
| B9J08_002852 | *ERG4* | Lipid metabolism | TRUE | -1,646 | 0,296 | 0,500 | 0,211 | 2,118 | 0,046 | -0,028 | 0,974 |
| B9J08_003993 | *HRT2* | | FALSE | -0,465 | 0,322 | 1,470 | 0,068 | 2,085 | 0,050 | 0,149 | 0,680 |
| B9J08_003191 | | | FALSE | -0,680 | 0,146 | 0,981 | 0,031 | 2,045 | 0,037 | 0,384 | 0,286 |
| B9J08_002624 | | | FALSE | -0,427 | 0,316 | 0,396 | 0,164 | 2,036 | 0,010 | 1,213 | 0,249 |
| B9J08_002736 | | Protein catabolism; Response to stress | TRUE | -0,407 | 0,541 | 1,127 | 0,112 | 2,021 | 0,014 | 0,487 | 0,627 |
| B9J08_001514 | | | FALSE | -1,644 | 0,057 | 0,076 | 0,360 | 1,982 | 0,006 | 0,262 | 0,293 |
| B9J08_003657 | *ZRT2* | Transmembrane transport | TRUE | -2,841 | 0,194 | -0,895 | 0,085 | 1,957 | 0,013 | 0,010 | 0,988 |
| B9J08_003924 | *IRA2* | | FALSE | -0,607 | 0,381 | 0,672 | 0,040 | 1,930 | 0,017 | 0,652 | 0,345 |
| B9J08_002799 | | | FALSE | 0,190 | 0,774 | 1,639 | 0,065 | 1,902 | 0,028 | 0,454 | 0,208 |
| B9J08_005198 | *RGS2* | | FALSE | -1,174 | 0,095 | 0,725 | 0,058 | 1,898 | 0,029 | -0,001 | 0,996 |
| B9J08_000657 | *ADP1* | | FALSE | -1,022 | 0,185 | 1,260 | 0,063 | 1,896 | 0,043 | -0,386 | 0,483 |
| B9J08_000962 | | Vesicle-mediated transport | TRUE | -1,394 | 0,089 | 0,121 | 0,212 | 1,870 | 0,046 | 0,355 | 0,333 |
| B9J08_001899 | *MLT1* | Response to stress; Transmembrane transport | TRUE | -1,345 | 0,144 | 0,896 | 0,057 | 1,854 | 0,040 | -0,387 | 0,459 |
| B9J08_001650 | | | FALSE | -0,535 | 0,405 | 0,851 | 0,091 | 1,829 | 0,035 | 0,442 | 0,318 |
| B9J08_002524 | *GAT1* | | FALSE | -0,180 | 0,658 | 0,895 | 0,079 | 1,811 | 0,036 | 0,737 | 0,208 |
| B9J08_005018 | *CDC21* | | FALSE | -0,746 | 0,200 | 0,229 | 0,266 | 1,806 | 0,019 | 0,831 | 0,240 |
| B9J08_004642 | *YBP1* | Response to stress | TRUE | -0,679 | 0,391 | 0,879 | 0,060 | 1,806 | 0,024 | 0,248 | 0,672 |
| B9J08_005106 | *NOC2* | Response to stress | TRUE | -0,523 | 0,557 | 0,741 | 0,026 | 1,805 | 0,021 | 0,541 | 0,501 |
| B9J08_002986 | *GLC3* | Carbohydrate metabolism | TRUE | -1,679 | 0,092 | -0,149 | 0,343 | 1,785 | 0,025 | 0,255 | 0,459 |
| B9J08_004758 | | | FALSE | -0,721 | 0,060 | 0,841 | 0,029 | 1,763 | 0,019 | 0,201 | 0,352 |
| B9J08_004847 | *VTC3* | | FALSE | -0,825 | 0,147 | 0,483 | 0,059 | 1,752 | 0,018 | 0,445 | 0,309 |
| B9J08_005234 | | | FALSE | -1,310 | 0,233 | 0,141 | 0,539 | 1,746 | 0,040 | 0,294 | 0,627 |
| B9J08_003598 | *CSH3* | Response to stress | TRUE | -0,983 | 0,408 | 0,659 | 0,134 | 1,739 | 0,037 | 0,098 | 0,916 |
| B9J08_005380 | *SUR7* | Cell wall organization / biogenesis; Vesicle-mediated transport; Cytoskeleton organization; Lipid metabolism; Response to stress | TRUE | -1,873 | 0,241 | -0,280 | 0,240 | 1,738 | 0,038 | 0,145 | 0,834 |
| B9J08_005439 | | | FALSE | -0,838 | 0,271 | 0,442 | 0,236 | 1,722 | 0,044 | 0,442 | 0,472 |
| B9J08_003360 | | DNA replication / cell cycle | TRUE | -1,613 | 0,290 | 0,067 | 0,757 | 1,719 | 0,049 | 0,039 | 0,966 |
| B9J08_000397 | *PEX11* | Lipid metabolism | TRUE | -0,832 | 0,263 | 0,996 | 0,045 | 1,706 | 0,018 | -0,122 | 0,813 |
| B9J08_005396 | *VPS15* | Vesicle-mediated transport | TRUE | -0,508 | 0,481 | 0,671 | 0,091 | 1,702 | 0,049 | 0,523 | 0,459 |
| B9J08_002245 | | | FALSE | -2,024 | 0,054 | -0,170 | 0,355 | 1,698 | 0,038 | -0,157 | 0,479 |
| B9J08_003925 | *IRA2* | | FALSE | -0,466 | 0,562 | 1,146 | 0,067 | 1,695 | 0,033 | 0,082 | 0,924 |
| B9J08_005062 | | | FALSE | -2,760 | 0,126 | -0,533 | 0,092 | 1,693 | 0,045 | -0,535 | 0,431 |
| B9J08_005143 | | | FALSE | -0,339 | 0,279 | 0,856 | 0,094 | 1,685 | 0,018 | 0,491 | 0,512 |
| B9J08_004065 | | | FALSE | -1,911 | 0,088 | -0,517 | 0,112 | 1,680 | 0,028 | 0,286 | 0,461 |
| B9J08_000007 | | | FALSE | -1,185 | 0,285 | 0,427 | 0,110 | 1,679 | 0,036 | 0,067 | 0,926 |
| B9J08_001788 | | | FALSE | -0,626 | 0,273 | 0,783 | 0,029 | 1,658 | 0,013 | 0,249 | 0,585 |
| B9J08_001532 | | | FALSE | -1,429 | 0,083 | 0,288 | 0,068 | 1,648 | 0,020 | -0,069 | 0,736 |
| B9J08_001592 | *VRG4* | Transmembrane transport | TRUE | -0,394 | 0,672 | 1,143 | 0,077 | 1,621 | 0,043 | 0,084 | 0,937 |
| B9J08_000862 | *TBF1* | | FALSE | -0,426 | 0,584 | 0,891 | 0,029 | 1,614 | 0,038 | 0,297 | 0,652 |
| B9J08_002104 | *MDG1* | | FALSE | -1,147 | 0,075 | 0,106 | 0,302 | 1,613 | 0,017 | 0,360 | 0,286 |
| B9J08_001937 | *NIT3* | | FALSE | -0,959 | 0,177 | 0,583 | 0,138 | 1,606 | 0,046 | 0,063 | 0,906 |

| Gene | Symbol | All processes | Included in Fig. 3e | wt-CAS vs wt-ND log2(FC) | wt-CAS vs wt-ND p(adj) | *hog1*Δ-CAS vs hog1Δ-ND log2(FC) | *hog1*Δ-CAS vs hog1Δ-ND p(adj) | *hog1*Δ-CAS vs wt-CAS log2(FC) | *hog1*Δ-CAS vs wt-CAS p(adj) | *hog1*Δ-ND vs wt-ND log2(FC) | *hog1*Δ-ND vs wt-ND p(adj) |
|---|---|---|---|---|---|---|---|---|---|---|---|
| B9J08_003729 | | | FALSE | -0,800 | 0,409 | 0,495 | 0,140 | 1,587 | 0,028 | 0,292 | 0,697 |
| B9J08_002454 | *HYM1* | Cell wall organization / biogenesis; Response to stress | TRUE | 0,033 | 0,951 | 0,888 | 0,098 | 1,566 | 0,038 | 0,710 | 0,271 |
| B9J08_001660 | *AMO2* | | FALSE | -3,190 | 0,060 | -0,630 | 0,057 | 1,551 | 0,044 | -1,009 | 0,233 |
| B9J08_004833 | *GLG2* | | FALSE | -1,209 | 0,108 | 0,163 | 0,296 | 1,543 | 0,028 | 0,171 | 0,556 |
| B9J08_004515 | | | FALSE | -1,257 | 0,081 | 0,863 | 0,031 | 1,537 | 0,022 | -0,583 | 0,268 |
| B9J08_004856 | *RAD23* | Response to stress | TRUE | -0,866 | 0,146 | 0,942 | 0,051 | 1,520 | 0,028 | -0,288 | 0,476 |
| B9J08_001741 | *PXP2* | Lipid metabolism | TRUE | -2,265 | 0,168 | -1,015 | 0,090 | 1,519 | 0,040 | 0,268 | 0,633 |
| B9J08_004976 | | | FALSE | 0,215 | 0,693 | 1,087 | 0,084 | 1,509 | 0,039 | 0,637 | 0,270 |
| B9J08_000273 | | | FALSE | -0,554 | 0,293 | 0,668 | 0,058 | 1,507 | 0,022 | 0,285 | 0,550 |
| B9J08_005175 | | | FALSE | -0,905 | 0,401 | 0,304 | 0,186 | 1,493 | 0,022 | 0,283 | 0,709 |
| B9J08_001811 | *YVC1* | Transmembrane transport | TRUE | -0,554 | 0,470 | 0,020 | 0,919 | 1,475 | 0,036 | 0,901 | 0,286 |
| B9J08_003338 | *RFC5* | Response to stress; DNA replication / cell cycle | TRUE | 0,079 | 0,783 | 1,129 | 0,065 | 1,473 | 0,030 | 0,423 | 0,164 |
| B9J08_000901 | | | FALSE | -1,081 | 0,074 | 0,692 | 0,019 | 1,458 | 0,012 | -0,315 | 0,373 |
| B9J08_001111 | | | FALSE | -0,304 | 0,507 | 0,598 | 0,100 | 1,435 | 0,050 | 0,532 | 0,345 |
| B9J08_000493 | | | FALSE | -0,533 | 0,375 | 0,547 | 0,138 | 1,416 | 0,025 | 0,336 | 0,598 |
| B9J08_003206 | | Cytoskeleton organization; Translation / ribosome biogenesis | TRUE | -0,798 | 0,200 | -0,091 | 0,659 | 1,410 | 0,038 | 0,703 | 0,268 |
| B9J08_005228 | *TAL1* | Carbohydrate metabolism | TRUE | -1,758 | 0,113 | 0,392 | 0,140 | 1,408 | 0,043 | -0,741 | 0,373 |
| B9J08_005348 | | | FALSE | -0,414 | 0,685 | 0,701 | 0,047 | 1,404 | 0,022 | 0,290 | 0,765 |
| B9J08_004840 | *YCP4* | Response to stress | TRUE | -1,938 | 0,160 | -0,182 | 0,418 | 1,370 | 0,019 | -0,386 | 0,540 |
| B9J08_005155 | | | FALSE | -0,355 | 0,630 | 0,977 | 0,066 | 1,365 | 0,024 | 0,033 | 0,970 |
| B9J08_005322 | | Transmembrane transport | TRUE | -0,738 | 0,051 | 0,122 | 0,529 | 1,364 | 0,023 | 0,504 | 0,321 |
| B9J08_001025 | *GSY1* | Carbohydrate metabolism | TRUE | -1,006 | 0,097 | -0,242 | 0,187 | 1,361 | 0,028 | 0,597 | 0,235 |
| B9J08_003025 | | | FALSE | -0,395 | 0,365 | 0,429 | 0,054 | 1,358 | 0,025 | 0,534 | 0,307 |
| B9J08_001391 | | | FALSE | -0,409 | 0,477 | 0,695 | 0,065 | 1,338 | 0,049 | 0,234 | 0,468 |
| B9J08_005339 | | Carbohydrate metabolism | TRUE | -1,649 | 0,073 | 0,062 | 0,603 | 1,325 | 0,033 | -0,386 | 0,268 |
| B9J08_005441 | | Transmembrane transport | TRUE | -0,182 | 0,762 | 0,895 | 0,066 | 1,325 | 0,040 | 0,248 | 0,695 |
| B9J08_002547 | | | FALSE | -0,162 | 0,789 | 0,887 | 0,051 | 1,324 | 0,041 | 0,274 | 0,673 |
| B9J08_001803 | *NOP1* | | FALSE | 0,089 | 0,560 | 0,910 | 0,032 | 1,323 | 0,014 | 0,502 | 0,143 |
| B9J08_003390 | *PCL7* | | FALSE | -0,296 | 0,239 | 0,824 | 0,048 | 1,321 | 0,028 | 0,200 | 0,406 |
| B9J08_003892 | *ORC4* | DNA replication / cell cycle | TRUE | 0,323 | 0,416 | 0,919 | 0,031 | 1,318 | 0,021 | 0,722 | 0,279 |
| B9J08_003572 | *VAN1* | Cell wall organization / biogenesis; Carbohydrate metabolism | TRUE | -0,550 | 0,427 | 0,800 | 0,097 | 1,318 | 0,046 | -0,032 | 0,962 |
| B9J08_002234 | | | FALSE | -1,927 | 0,186 | -0,258 | 0,451 | 1,315 | 0,018 | -0,354 | 0,627 |
| B9J08_001728 | | | FALSE | -0,554 | 0,557 | 0,710 | 0,045 | 1,315 | 0,022 | 0,050 | 0,959 |
| B9J08_000052 | | | FALSE | -0,219 | 0,552 | 1,171 | 0,054 | 1,304 | 0,028 | -0,086 | 0,860 |
| B9J08_005007 | *ERG7* | Lipid metabolism | TRUE | -0,040 | 0,927 | 0,981 | 0,073 | 1,298 | 0,046 | 0,277 | 0,598 |
| B9J08_003276 | *GPH1* | Carbohydrate metabolism | TRUE | -0,635 | 0,194 | 0,599 | 0,060 | 1,293 | 0,017 | 0,059 | 0,850 |
| B9J08_002490 | *SNU66* | | FALSE | -0,317 | 0,327 | 0,958 | 0,049 | 1,291 | 0,032 | 0,016 | 0,957 |
| B9J08_005201 | *PYC2* | | FALSE | -0,490 | 0,098 | 0,764 | 0,084 | 1,288 | 0,049 | 0,035 | 0,822 |
| B9J08_003167 | | | FALSE | -1,090 | 0,095 | 0,337 | 0,114 | 1,284 | 0,032 | -0,144 | 0,558 |
| B9J08_003428 | *TRI1* | | FALSE | -0,803 | 0,191 | 0,008 | 0,957 | 1,283 | 0,042 | 0,472 | 0,307 |
| B9J08_005451 | *DOS2* | | FALSE | -0,885 | 0,184 | 0,501 | 0,092 | 1,271 | 0,019 | -0,115 | 0,792 |
| B9J08_003702 | *CDG1* | | FALSE | -2,157 | 0,105 | -0,727 | 0,031 | 1,263 | 0,043 | -0,167 | 0,633 |

| Gene | Symbol | All processes | Included in Fig. 3e | wt-CAS vs wt-ND log2(FC) | wt-CAS vs wt-ND p(adj) | ***hog1*Δ-CAS vs hog1Δ-ND log2(FC)** | ***hog1*Δ-CAS vs hog1Δ-ND p(adj)** | ***hog1*Δ-CAS vs wt-CAS log2(FC)** | ***hog1*Δ-CAS vs wt-CAS p(adj)** | ***hog1*Δ-ND vs wt-ND log2(FC)** | ***hog1*Δ-ND vs wt-ND p(adj)** |
|---|---|---|---|---|---|---|---|---|---|---|---|
| B9J08_000393 | | | FALSE | -1,064 | 0,074 | -0,169 | 0,170 | 1,263 | 0,012 | 0,368 | 0,268 |
| B9J08_003173 | | | FALSE | -1,135 | 0,267 | -0,033 | 0,820 | 1,262 | 0,015 | 0,161 | 0,777 |
| B9J08_002687 | *ZCF29* | Response to stress | TRUE | -0,699 | 0,361 | 0,454 | 0,091 | 1,252 | 0,022 | 0,100 | 0,874 |
| B9J08_002662 | | | FALSE | -0,811 | 0,073 | 0,781 | 0,050 | 1,252 | 0,032 | -0,340 | 0,320 |
| B9J08_004825 | | Transmembrane transport | TRUE | -1,420 | 0,227 | -0,124 | 0,361 | 1,247 | 0,026 | -0,049 | 0,942 |
| B9J08_000474 | | Lipid metabolism | TRUE | -0,610 | 0,392 | 0,496 | 0,066 | 1,246 | 0,022 | 0,140 | 0,801 |
| B9J08_004827 | *ATG9* | | FALSE | -0,133 | 0,819 | 0,137 | 0,546 | 1,245 | 0,028 | 0,976 | 0,268 |
| B9J08_000779 | | | FALSE | -0,685 | 0,230 | 0,211 | 0,269 | 1,240 | 0,037 | 0,345 | 0,439 |
| B9J08_004667 | | | FALSE | -1,797 | 0,096 | -0,817 | 0,079 | 1,238 | 0,019 | 0,258 | 0,454 |
| B9J08_002816 | *YMC1* | Transmembrane transport | TRUE | -0,294 | 0,600 | 0,856 | 0,027 | 1,229 | 0,025 | 0,080 | 0,892 |
| B9J08_004289 | | | FALSE | -0,351 | 0,299 | 0,630 | 0,052 | 1,222 | 0,022 | 0,241 | 0,286 |
| B9J08_002000 | | | FALSE | -0,403 | 0,618 | 0,814 | 0,061 | 1,200 | 0,042 | -0,017 | 0,985 |
| B9J08_004309 | *PFK1* | Carbohydrate metabolism; Cellular respiration | TRUE | -0,505 | 0,125 | 0,942 | 0,012 | 1,199 | 0,015 | -0,248 | 0,352 |
| B9J08_000845 | *NIT2* | | FALSE | -1,324 | 0,061 | 0,222 | 0,064 | 1,192 | 0,014 | -0,354 | 0,299 |
| B9J08_002082 | | | FALSE | -0,909 | 0,075 | -0,038 | 0,543 | 1,183 | 0,049 | 0,311 | 0,278 |
| B9J08_003095 | | | FALSE | -0,369 | 0,266 | 0,533 | 0,059 | 1,181 | 0,024 | 0,279 | 0,361 |
| B9J08_000729 | *SCH9* | Response to stress | TRUE | -0,599 | 0,092 | 0,370 | 0,106 | 1,177 | 0,047 | 0,208 | 0,421 |
| B9J08_000889 | | | FALSE | -1,097 | 0,073 | 0,164 | 0,543 | 1,170 | 0,031 | -0,090 | 0,813 |
| B9J08_003669 | *RPN3* | Protein catabolism | TRUE | -0,469 | 0,397 | 0,561 | 0,065 | 1,167 | 0,019 | 0,137 | 0,764 |
| B9J08_005455 | *DOA4* | | FALSE | 0,237 | 0,557 | 0,645 | 0,036 | 1,165 | 0,021 | 0,756 | 0,255 |
| B9J08_000080 | *AVT1* | | FALSE | -0,013 | 0,980 | 0,753 | 0,037 | 1,164 | 0,043 | 0,397 | 0,497 |
| B9J08_004062 | | | FALSE | -0,523 | 0,071 | 0,406 | 0,108 | 1,163 | 0,013 | 0,234 | 0,445 |
| B9J08_004603 | | | FALSE | -0,978 | 0,345 | 0,516 | 0,574 | 1,159 | 0,043 | -0,335 | 0,816 |
| B9J08_002340 | | | FALSE | -0,069 | 0,853 | 0,872 | 0,064 | 1,140 | 0,050 | 0,199 | 0,633 |
| B9J08_001956 | *SPS20* | | FALSE | -0,738 | 0,183 | 0,466 | 0,141 | 1,133 | 0,041 | -0,070 | 0,840 |
| B9J08_003531 | *HIS5* | | FALSE | -0,293 | 0,202 | 0,682 | 0,020 | 1,133 | 0,013 | 0,158 | 0,422 |
| B9J08_003732 | *RIM11* | Response to stress | TRUE | -0,328 | 0,557 | 0,386 | 0,087 | 1,130 | 0,049 | 0,416 | 0,454 |
| B9J08_001489 | *PHO113* | | FALSE | -3,193 | 0,175 | -2,286 | 0,062 | 1,124 | 0,010 | 0,218 | 0,717 |
| B9J08_000164 | *CDR1* | Response to stress; Transmembrane transport | TRUE | -0,312 | 0,513 | 0,337 | 0,114 | 1,118 | 0,025 | 0,469 | 0,409 |
| B9J08_004554 | *SEC24* | Vesicle-mediated transport | TRUE | -0,554 | 0,331 | 0,645 | 0,013 | 1,117 | 0,025 | -0,082 | 0,862 |
| B9J08_000501 | *PHM7* | | FALSE | -0,202 | 0,700 | 0,608 | 0,047 | 1,107 | 0,025 | 0,297 | 0,609 |
| B9J08_000517 | *FET31* | | FALSE | 0,292 | 0,177 | 0,130 | 0,525 | 1,098 | 0,019 | 1,260 | 0,255 |
| B9J08_001144 | *MED4* | | FALSE | -0,257 | 0,341 | 0,557 | 0,085 | 1,090 | 0,031 | 0,277 | 0,252 |
| B9J08_003048 | *NIC96* | | FALSE | 0,083 | 0,747 | 0,868 | 0,029 | 1,089 | 0,028 | 0,305 | 0,431 |
| B9J08_002477 | *PGM2* | Carbohydrate metabolism | TRUE | -0,698 | 0,156 | 0,584 | 0,058 | 1,088 | 0,037 | -0,194 | 0,523 |
| B9J08_003806 | | | FALSE | -1,038 | 0,096 | 0,317 | 0,337 | 1,084 | 0,047 | -0,271 | 0,540 |
| B9J08_003451 | | | FALSE | -0,326 | 0,442 | 0,287 | 0,243 | 1,082 | 0,037 | 0,469 | 0,387 |
| B9J08_000750 | | | FALSE | 0,232 | 0,164 | 0,781 | 0,058 | 1,079 | 0,045 | 0,530 | 0,196 |
| B9J08_002692 | | | FALSE | 0,014 | 0,716 | 0,579 | 0,062 | 1,077 | 0,050 | 0,512 | 0,244 |
| B9J08_001627 | *VPS9* | Vesicle-mediated transport | TRUE | -0,024 | 0,810 | 0,476 | 0,044 | 1,063 | 0,004 | 0,562 | 0,187 |
| B9J08_001748 | *TCC1* | | FALSE | -0,497 | 0,093 | 0,737 | 0,045 | 1,061 | 0,032 | -0,173 | 0,454 |
| B9J08_003992 | | | FALSE | -1,356 | 0,085 | -0,615 | 0,270 | 1,056 | 0,033 | 0,315 | 0,608 |
| B9J08_000686 | *ARG1* | | FALSE | -0,334 | 0,098 | 0,272 | 0,072 | 1,054 | 0,008 | 0,448 | 0,208 |
| B9J08_000513 | *TOM1* | | FALSE | -0,641 | 0,425 | 0,322 | 0,119 | 1,049 | 0,049 | 0,086 | 0,901 |

| Gene | Symbol | All processes | Included in Fig. 3e | wt-CAS vs wt-ND log2(FC) | wt-CAS vs wt-ND p(adj) | *hog1*Δ-CAS vs hog1Δ-ND log2(FC) | *hog1*Δ-CAS vs hog1Δ-ND p(adj) | *hog1*Δ-CAS vs wt-CAS log2(FC) | *hog1*Δ-CAS vs wt-CAS p(adj) | *hog1*Δ-ND vs wt-ND log2(FC) | *hog1*Δ-ND vs wt-ND p(adj) |
|---|---|---|---|---|---|---|---|---|---|---|---|
| B9J08_002166 | | | FALSE | -1,721 | 0,085 | -0,622 | 0,079 | 1,031 | 0,029 | -0,068 | 0,822 |
| B9J08_000162 | *RAD7* | Response to stress | TRUE | -0,501 | 0,194 | 0,647 | 0,070 | 1,029 | 0,043 | -0,118 | 0,645 |
| B9J08_003898 | | | FALSE | -1,585 | 0,075 | -0,262 | 0,378 | 1,026 | 0,028 | -0,297 | 0,466 |
| B9J08_002898 | *TAF60* | | FALSE | -0,275 | 0,538 | 0,114 | 0,660 | 1,026 | 0,022 | 0,637 | 0,365 |
| B9J08_004540 | *CYR1* | Response to stress | TRUE | 0,076 | 0,773 | 0,818 | 0,050 | 1,021 | 0,028 | 0,279 | 0,454 |
| B9J08_003871 | | Transmembrane transport | TRUE | -0,290 | 0,648 | 0,184 | 0,376 | 1,019 | 0,049 | 0,545 | 0,459 |
| B9J08_003042 | | | FALSE | 0,502 | 0,339 | 0,766 | 0,046 | 1,016 | 0,043 | 0,753 | 0,313 |
| B9J08_004756 | | | FALSE | -0,962 | 0,101 | -0,036 | 0,803 | 1,008 | 0,043 | 0,083 | 0,582 |
| B9J08_002844 | *ARP9* | | FALSE | -0,489 | 0,316 | 0,436 | 0,018 | 1,004 | 0,008 | 0,078 | 0,829 |
| B9J08_002192 | *POM152* | | FALSE | 0,736 | 0,092 | -0,254 | 0,213 | -1,002 | 0,015 | -0,011 | 0,973 |
| B9J08_005334 | *STE50* | | FALSE | 0,928 | 0,064 | -0,231 | 0,149 | -1,003 | 0,013 | 0,156 | 0,539 |
| B9J08_002916 | | | FALSE | 1,149 | 0,067 | -0,270 | 0,400 | -1,006 | 0,042 | 0,413 | 0,255 |
| B9J08_003204 | | | FALSE | 0,273 | 0,401 | -0,345 | 0,415 | -1,025 | 0,019 | -0,407 | 0,479 |
| B9J08_001191 | | | FALSE | 0,522 | 0,210 | -0,789 | 0,068 | -1,036 | 0,031 | 0,274 | 0,539 |
| B9J08_000013 | *ILV3* | | FALSE | 0,160 | 0,472 | -0,430 | 0,122 | -1,051 | 0,025 | -0,461 | 0,323 |
| B9J08_001910 | *MSI3* | Protein folding; Protein catabolism; Response to stress | TRUE | 0,178 | 0,295 | -0,809 | 0,030 | -1,059 | 0,040 | -0,071 | 0,665 |
| B9J08_003466 | *MDH1-3* | Cellular respiration | TRUE | 0,131 | 0,280 | -0,961 | 0,020 | -1,067 | 0,025 | 0,025 | 0,803 |
| B9J08_000640 | | | FALSE | 0,324 | 0,506 | -0,492 | 0,220 | -1,069 | 0,012 | -0,253 | 0,711 |
| B9J08_000692 | *YRB1* | Cytoskeleton organization | TRUE | 0,057 | 0,685 | -0,820 | 0,131 | -1,077 | 0,020 | -0,199 | 0,598 |
| B9J08_005354 | *ORC6* | DNA replication / cell cycle | TRUE | 0,411 | 0,321 | -0,998 | 0,171 | -1,083 | 0,028 | 0,326 | 0,627 |
| B9J08_005147 | | | FALSE | 0,892 | 0,042 | -0,569 | 0,099 | -1,092 | 0,018 | 0,369 | 0,353 |
| B9J08_004146 | *RPL14* | Translation / ribosome biogenesis | TRUE | 0,793 | 0,124 | -0,232 | 0,155 | -1,096 | 0,022 | -0,071 | 0,881 |
| B9J08_004595 | | | FALSE | 0,832 | 0,008 | -0,518 | 0,106 | -1,096 | 0,045 | 0,254 | 0,250 |
| B9J08_004648 | | | FALSE | -0,266 | 0,156 | -0,745 | 0,054 | -1,116 | 0,020 | -0,636 | 0,196 |
| B9J08_004192 | *GDH2* | | FALSE | 1,313 | 0,055 | 0,143 | 0,378 | -1,118 | 0,034 | 0,051 | 0,845 |
| B9J08_005362 | | | FALSE | 1,130 | 0,083 | -0,168 | 0,154 | -1,120 | 0,012 | 0,177 | 0,627 |
| B9J08_002329 | | | FALSE | 0,379 | 0,240 | -0,631 | 0,081 | -1,122 | 0,040 | -0,112 | 0,717 |
| B9J08_001908 | *MNT2* | Cell wall organization / biogenesis | TRUE | 1,036 | 0,073 | -0,103 | 0,662 | -1,132 | 0,039 | 0,007 | 0,985 |
| B9J08_003609 | *GCV2* | | FALSE | -0,309 | 0,200 | -0,808 | 0,085 | -1,149 | 0,043 | -0,649 | 0,161 |
| B9J08_004301 | *HYS2* | Response to stress; DNA replication / cell cycle | TRUE | 0,594 | 0,138 | 0,064 | 0,828 | -1,155 | 0,025 | -0,625 | 0,351 |
| B9J08_002013 | | | FALSE | -0,279 | 0,081 | -0,853 | 0,038 | -1,157 | 0,025 | -0,583 | 0,143 |
| B9J08_002473 | | Translation / ribosome biogenesis | TRUE | 1,419 | 0,124 | -0,116 | 0,508 | -1,167 | 0,043 | 0,368 | 0,692 |
| B9J08_004516 | *SPT20* | | FALSE | 0,349 | 0,287 | -1,285 | 0,084 | -1,177 | 0,018 | 0,457 | 0,409 |
| B9J08_001202 | *NUO2* | DNA replication / cell cycle | TRUE | 0,440 | 0,097 | -0,494 | 0,094 | -1,185 | 0,010 | -0,251 | 0,345 |
| B9J08_002669 | *PCK1* | Carbohydrate metabolism | TRUE | 1,505 | 0,148 | 0,350 | 0,244 | -1,187 | 0,014 | -0,032 | 0,978 |
| B9J08_003707 | *PRD1* | | FALSE | 0,669 | 0,108 | -0,461 | 0,163 | -1,188 | 0,022 | -0,058 | 0,867 |
| B9J08_005140 | | | FALSE | 2,838 | 0,051 | 0,591 | 0,508 | -1,199 | 0,046 | 1,048 | 0,633 |
| B9J08_001633 | | | FALSE | 0,039 | 0,878 | -0,712 | 0,073 | -1,204 | 0,026 | -0,453 | 0,397 |
| B9J08_000075 | *IMH3* | Response to stress | TRUE | 0,468 | 0,078 | -0,651 | 0,075 | -1,209 | 0,022 | -0,091 | 0,680 |
| B9J08_001302 | *BRE1* | Response to stress | TRUE | 0,189 | 0,313 | -1,308 | 0,057 | -1,217 | 0,028 | 0,279 | 0,436 |
| B9J08_000650 | *LSC1* | | FALSE | 0,254 | 0,096 | -0,242 | 0,214 | -1,225 | 0,012 | -0,730 | 0,143 |
| B9J08_001744 | | | FALSE | 0,565 | 0,113 | -0,446 | 0,097 | -1,226 | 0,037 | -0,215 | 0,476 |
| B9J08_001475 | | Carbohydrate metabolism | TRUE | 1,356 | 0,056 | -0,283 | 0,370 | -1,250 | 0,025 | 0,389 | 0,459 |
| B9J08_003553 | *DLD1* | | FALSE | 0,216 | 0,450 | -0,539 | 0,187 | -1,256 | 0,034 | -0,500 | 0,341 |

| Gene | Symbol | All processes | Included in Fig. 3e | wt-CAS vs wt-ND log2(FC) | wt-CAS vs wt-ND p(adj) | ***hog1*Δ-CAS vs hog1Δ-ND log2(FC)** | ***hog1*Δ-CAS vs hog1Δ-ND p(adj)** | ***hog1*Δ-CAS vs wt-CAS log2(FC)** | ***hog1*Δ-CAS vs wt-CAS p(adj)** | ***hog1*Δ-ND vs wt-ND log2(FC)** | ***hog1*Δ-ND vs wt-ND p(adj)** |
|---|---|---|---|---|---|---|---|---|---|---|---|
| B9J08_002301 | *MSS116* | | FALSE | 0,873 | 0,095 | -0,168 | 0,425 | -1,256 | 0,049 | -0,214 | 0,627 |
| B9J08_001743 | *PAL1* | | FALSE | 0,954 | 0,108 | -0,369 | 0,134 | -1,260 | 0,040 | 0,063 | 0,912 |
| B9J08_003356 | *SAK1* | | FALSE | 1,050 | 0,056 | -0,426 | 0,138 | -1,270 | 0,020 | 0,206 | 0,383 |
| B9J08_002422 | | | FALSE | 1,444 | 0,092 | -0,609 | 0,243 | -1,277 | 0,013 | 0,775 | 0,387 |
| B9J08_000619 | *COQ4* | | FALSE | 0,288 | 0,377 | -0,695 | 0,053 | -1,287 | 0,019 | -0,304 | 0,517 |
| B9J08_002876 | *CMD1* | Cytoskeleton organization; Lipid metabolism; DNA replication / cell cycle | TRUE | 1,140 | 0,060 | 0,211 | 0,189 | -1,291 | 0,049 | -0,362 | 0,472 |
| B9J08_005296 | *FGR39* | Response to stress | TRUE | 1,134 | 0,085 | Inf | 0,066 | -1,295 | 0,024 | -Inf | 0,257 |
| B9J08_003110 | | | FALSE | 2,415 | 0,071 | 2,038 | 0,067 | -1,315 | 0,022 | -0,938 | 0,669 |
| B9J08_005044 | *YME1* | | FALSE | 0,923 | 0,056 | -0,457 | 0,120 | -1,324 | 0,017 | 0,056 | 0,646 |
| B9J08_003126 | *QCR8* | | FALSE | -0,077 | 0,826 | -0,438 | 0,289 | -1,325 | 0,017 | -0,964 | 0,310 |
| B9J08_004528 | | | FALSE | 1,720 | 0,050 | -0,154 | 0,662 | -1,330 | 0,029 | 0,544 | 0,268 |
| B9J08_003610 | *TEF1* | Translation / ribosome biogenesis | TRUE | 0,834 | 0,074 | -0,581 | 0,041 | -1,336 | 0,027 | 0,079 | 0,801 |
| B9J08_002307 | | Translation / ribosome biogenesis | TRUE | 0,007 | 0,973 | -1,015 | 0,186 | -1,337 | 0,021 | -0,315 | 0,564 |
| B9J08_003092 | *MNN21* | Cell wall organization / biogenesis; Carbohydrate metabolism | TRUE | 0,563 | 0,343 | -1,348 | 0,084 | -1,343 | 0,044 | 0,569 | 0,459 |
| B9J08_000601 | | | FALSE | 1,025 | 0,113 | -0,719 | 0,139 | -1,351 | 0,028 | 0,393 | 0,531 |
| B9J08_000120 | *TSR1* | Translation / ribosome biogenesis | TRUE | 0,735 | 0,116 | -0,577 | 0,416 | -1,354 | 0,025 | -0,042 | 0,961 |
| B9J08_000613 | *GLT1* | | FALSE | -0,301 | 0,287 | -0,760 | 0,010 | -1,368 | 0,037 | -0,910 | 0,262 |
| B9J08_004356 | | Translation / ribosome biogenesis | TRUE | 0,512 | 0,208 | -1,306 | 0,123 | -1,374 | 0,048 | 0,443 | 0,495 |
| B9J08_004737 | *LAT1* | | FALSE | -0,679 | 0,214 | -0,972 | 0,100 | -1,379 | 0,045 | -1,086 | 0,271 |
| B9J08_002188 | *DFG5* | Cell wall organization / biogenesis; Carbohydrate metabolism | TRUE | 1,685 | 0,067 | 2,278 | 0,107 | -1,385 | 0,038 | -1,978 | 0,341 |
| B9J08_002860 | *FKH2* | Response to stress | TRUE | 0,917 | 0,084 | -0,717 | 0,119 | -1,386 | 0,031 | 0,249 | 0,550 |
| B9J08_002768 | *AHA1* | | FALSE | 0,867 | 0,084 | -0,823 | 0,104 | -1,388 | 0,025 | 0,303 | 0,222 |
| B9J08_001231 | | | FALSE | 0,603 | 0,084 | -0,673 | 0,094 | -1,389 | 0,013 | -0,113 | 0,722 |
| B9J08_004148 | *OP4* | | FALSE | 0,977 | 0,079 | -0,226 | 0,538 | -1,392 | 0,044 | -0,189 | 0,701 |
| B9J08_001143 | *HEM15* | | FALSE | 0,799 | 0,112 | 0,630 | 0,039 | -1,394 | 0,044 | -1,226 | 0,307 |
| B9J08_002693 | | | FALSE | 1,377 | 0,067 | -0,398 | 0,107 | -1,416 | 0,028 | 0,359 | 0,254 |
| B9J08_001632 | *NAB3* | | FALSE | 0,735 | 0,065 | -1,363 | 0,085 | -1,419 | 0,018 | 0,679 | 0,310 |
| B9J08_002738 | *WSC1* | | FALSE | Inf | 0,061 | Inf | 0,015 | -1,429 | 0,047 | | |
| B9J08_002764 | | | FALSE | 1,158 | 0,075 | -0,910 | 0,021 | -1,441 | 0,024 | 0,627 | 0,330 |
| B9J08_001245 | | | FALSE | 0,222 | 0,447 | 0,208 | 0,913 | -1,456 | 0,040 | -1,441 | 0,479 |
| B9J08_001992 | *KAP120* | | FALSE | 1,759 | 0,064 | -1,359 | 0,131 | -1,478 | 0,013 | 1,639 | 0,268 |
| B9J08_001224 | *ECM17* | | FALSE | 1,387 | 0,069 | -0,592 | 0,063 | -1,481 | 0,021 | 0,498 | 0,161 |
| B9J08_000755 | | | FALSE | 0,382 | 0,102 | -0,913 | 0,010 | -1,505 | 0,019 | -0,211 | 0,420 |
| B9J08_001889 | *CRC1* | | FALSE | 1,445 | 0,054 | 0,148 | 0,358 | -1,538 | 0,048 | -0,241 | 0,582 |
| B9J08_005428 | | | FALSE | 0,571 | 0,051 | -0,560 | 0,024 | -1,539 | 0,012 | -0,408 | 0,158 |
| B9J08_003829 | *RAD53* | Response to stress; DNA replication / cell cycle | TRUE | 1,131 | 0,078 | -0,720 | 0,263 | -1,540 | 0,041 | 0,311 | 0,627 |
| B9J08_000408 | *KRE5* | Cell wall organization / biogenesis; Carbohydrate metabolism; Response to stress | TRUE | 0,598 | 0,179 | -0,931 | 0,048 | -1,542 | 0,025 | -0,014 | 0,976 |
| B9J08_000858 | | | FALSE | 0,606 | 0,049 | -0,938 | 0,072 | -1,575 | 0,018 | -0,031 | 0,894 |
| B9J08_003975 | | Translation / ribosome biogenesis | TRUE | 1,182 | 0,063 | -0,654 | 0,066 | -1,582 | 0,014 | 0,255 | 0,585 |
| B9J08_004189 | *YFH1* | Response to stress | TRUE | -0,080 | 0,709 | -1,223 | 0,111 | -1,584 | 0,045 | -0,441 | 0,381 |
| B9J08_002712 | *PDI1* | | FALSE | 0,898 | 0,067 | 0,080 | 0,768 | -1,595 | 0,019 | -0,778 | 0,286 |

| Gene | Symbol | All processes | Included in Fig. 3e | wt-CAS vs wt-ND log2(FC) | wt-CAS vs wt-ND p(adj) | *hog1*Δ-CAS vs hog1Δ-ND log2(FC) | *hog1*Δ-CAS vs hog1Δ-ND p(adj) | *hog1*Δ-CAS vs wt-CAS log2(FC) | *hog1*Δ-CAS vs wt-CAS p(adj) | *hog1*Δ-ND vs wt-ND log2(FC) | *hog1*Δ-ND vs wt-ND p(adj) |
|---|---|---|---|---|---|---|---|---|---|---|---|
| B9J08_000384 | | Cell wall organization / biogenesis; Carbohydrate metabolism | TRUE | 0,610 | 0,147 | -0,397 | 0,039 | -1,596 | 0,042 | -0,589 | 0,404 |
| B9J08_000338 | *SEC3* | Vesicle-mediated transport | TRUE | 1,262 | 0,058 | -0,999 | 0,156 | -1,605 | 0,037 | 0,656 | 0,399 |
| B9J08_000672 | | | FALSE | 0,989 | 0,092 | Inf | 0,034 | -1,611 | 0,043 | -Inf | 0,262 |
| B9J08_002741 | | | FALSE | 0,440 | 0,356 | -1,116 | 0,063 | -1,622 | 0,035 | -0,067 | 0,916 |
| B9J08_000469 | *KGD2* | Cellular respiration | TRUE | -0,338 | 0,398 | -1,053 | 0,079 | -1,624 | 0,022 | -0,909 | 0,290 |
| B9J08_005410 | | | FALSE | 0,932 | 0,046 | -0,767 | 0,103 | -1,647 | 0,022 | 0,052 | 0,875 |
| B9J08_001369 | | | FALSE | 1,318 | 0,092 | -0,788 | 0,075 | -1,651 | 0,019 | 0,455 | 0,462 |
| B9J08_001708 | | Translation / ribosome biogenesis | TRUE | 1,217 | 0,120 | -0,737 | 0,059 | -1,747 | 0,012 | 0,207 | 0,721 |
| B9J08_004814 | *RIB2* | | FALSE | 1,226 | 0,071 | -1,198 | 0,089 | -1,753 | 0,041 | 0,671 | 0,325 |
| B9J08_000966 | | | FALSE | 1,911 | 0,051 | -0,330 | 0,336 | -1,791 | 0,021 | 0,450 | 0,363 |
| B9J08_001939 | *RIX7* | | FALSE | -0,050 | 0,807 | -2,055 | 0,080 | -1,879 | 0,020 | 0,126 | 0,773 |
| B9J08_003450 | *SET2* | | FALSE | 1,248 | 0,091 | -0,829 | 0,085 | -1,917 | 0,046 | 0,159 | 0,748 |
| B9J08_005098 | *LPI9* | | FALSE | 0,586 | 0,179 | -1,802 | 0,056 | -1,971 | 0,043 | 0,417 | 0,423 |
| B9J08_000659 | | | FALSE | 1,472 | 0,095 | -0,832 | 0,197 | -2,010 | 0,037 | 0,294 | 0,627 |
| B9J08_000293 | | Translation / ribosome biogenesis | TRUE | 0,514 | 0,124 | -1,674 | 0,167 | -2,017 | 0,028 | 0,171 | 0,794 |
| B9J08_000057 | *SNO1* | | FALSE | 1,299 | 0,056 | -0,384 | 0,411 | -2,021 | 0,032 | -0,337 | 0,453 |
| B9J08_001022 | | | FALSE | -0,135 | 0,522 | -2,026 | 0,084 | -2,041 | 0,031 | -0,149 | 0,700 |
| B9J08_002150 | *SLN1* | Response to stress | TRUE | 1,917 | 0,059 | -0,164 | 0,760 | -2,049 | 0,016 | 0,032 | 0,970 |
| B9J08_000527 | | | FALSE | -0,139 | 0,788 | -2,765 | 0,111 | -2,175 | 0,047 | 0,450 | 0,584 |
| B9J08_003021 | *UTP15* | Translation / ribosome biogenesis | TRUE | 0,344 | 0,250 | -2,046 | 0,190 | -2,180 | 0,042 | 0,210 | 0,809 |
| B9J08_003556 | | | FALSE | 2,256 | 0,051 | -0,337 | 0,220 | -2,187 | 0,041 | 0,406 | 0,625 |
| B9J08_002321 | *ARX1* | | FALSE | -1,041 | 0,067 | -2,822 | 0,050 | -2,244 | 0,032 | -0,463 | 0,299 |
| B9J08_001325 | | | FALSE | 5,766 | 0,056 | 4,330 | 0,111 | -2,272 | 0,013 | -0,836 | 0,274 |
| B9J08_000776 | *EHT1* | | FALSE | 1,019 | 0,065 | -0,660 | 0,800 | -2,304 | 0,020 | -0,625 | 0,786 |
| B9J08_002136 | *WOR2* | | FALSE | 0,206 | 0,867 | -0,510 | 0,841 | -2,341 | 0,032 | -1,625 | 0,617 |
| B9J08_003047 | | | FALSE | 1,749 | 0,103 | 0,274 | 0,898 | -2,350 | 0,049 | -0,875 | 0,690 |
| B9J08_005236 | *TOA2* | | FALSE | 0,434 | 0,159 | -1,847 | 0,050 | -2,383 | 0,024 | -0,103 | 0,738 |
| B9J08_001812 | | | FALSE | 2,685 | 0,051 | -1,450 | 0,139 | -2,498 | 0,028 | 1,637 | 0,269 |
| B9J08_004601 | *RNR1* | Response to stress; DNA replication / cell cycle | TRUE | 2,923 | 0,056 | 0,213 | 0,485 | -2,564 | 0,031 | 0,146 | 0,764 |
| B9J08_001463 | *MYO1* | DNA replication / cell cycle | TRUE | 0,919 | 0,056 | -0,957 | 0,604 | -2,629 | 0,045 | -0,753 | 0,627 |
| B9J08_001195 | | | FALSE | 2,360 | 0,059 | Inf | 0,303 | -2,786 | 0,024 | -Inf | 0,286 |
| B9J08_002533 | | | FALSE | 0,816 | 0,434 | -2,443 | 0,446 | -2,973 | 0,018 | 0,286 | 0,908 |
| B9J08_003680 | *DBR1* | | FALSE | 1,601 | 0,059 | -1,554 | 0,443 | -3,004 | 0,022 | 0,151 | 0,934 |
| B9J08_003505 | *SAS3* | | FALSE | 1,008 | 0,126 | -2,849 | 0,069 | -3,064 | 0,034 | 0,793 | 0,377 |
| B9J08_001375 | | | FALSE | 0,853 | 0,238 | -0,490 | 0,862 | -3,205 | 0,037 | -1,862 | 0,423 |
| B9J08_004586 | | | FALSE | 2,194 | 0,073 | -1,368 | 0,082 | -3,246 | 0,021 | 0,316 | 0,412 |
| B9J08_003519 | | | FALSE | 0,564 | 0,095 | -3,084 | 0,325 | -3,420 | 0,012 | 0,229 | 0,890 |
| B9J08_004999 | *RPP2A* | Translation / ribosome biogenesis | TRUE | 0,095 | 0,897 | -2,415 | 0,097 | -3,526 | 0,045 | -1,016 | 0,495 |
| B9J08_000017 | | | FALSE | 0,281 | 0,557 | -3,290 | 0,416 | -3,554 | 0,025 | 0,017 | 0,993 |
| B9J08_001131 | *FRK1* | DNA replication / cell cycle | TRUE | Inf | 0,056 | Inf | 0,508 | -3,826 | 0,019 | | |
| B9J08_005132 | *NMD3* | | FALSE | 1,062 | 0,127 | Inf | 0,386 | -4,697 | 0,045 | -Inf | 0,293 |
| B9J08_002563 | | | FALSE | Inf | 0,051 | Inf | 0,508 | -5,171 | 0,010 | | |
| B9J08_000087 | *CDC46* | DNA replication / cell cycle | TRUE | 1,194 | 0,441 | Inf | 0,322 | -5,387 | 0,040 | -Inf | 0,619 |
| B9J08_000374 | *SMI1B* | Cell wall organization / biogenesis | TRUE | 2,088 | 0,074 | Inf | 0,508 | -5,564 | 0,032 | -Inf | 0,226 |

| Gene | Symbol | All processes | Included in Fig. 3e | wt-CAS vs wt-ND log2(FC) | wt-CAS vs wt-ND p(adj) | *hog1*Δ-CAS vs hog1Δ-ND log2(FC) | *hog1*Δ-CAS vs hog1Δ-ND p(adj) | *hog1*Δ-CAS vs wt-CAS log2(FC) | *hog1*Δ-CAS vs wt-CAS p(adj) | *hog1*Δ-ND vs wt-ND log2(FC) | *hog1*Δ-ND vs wt-ND p(adj) |
|---|---|---|---|---|---|---|---|---|---|---|---|
| B9J08_000414 | *ZCF1* | | FALSE | 0,451 | 0,405 | -5,390 | 0,096 | -5,853 | 0,049 | -0,012 | 0,988 |

**Supplementary Table 5: Epidemiological HOG/CWI pathway variants identified in *C. auris*.** Catalogue of unique epidemiological variants in *SLN1, NIK1, SSK1, SSK2, PBS2, HOG1, MKC1* and *CEK1* derived from the genomic dataset of Gifford et al. (2026)[6]. Variants were identified relative to the *C. auris* B8441 reference genome (clade I; GCA_002759435.3). 'Variant' reports the reference sequence/contig and genomic coordinate corresponding to the variant annotation in the Gifford et al. dataset, while DNA change indicates the reference and alternative nucleotide alleles. Protein position and change indicate the corresponding amino-acid alteration. Clade occurrence is reported as the number of isolates carrying the variant relative to the number of isolates assessed within that clade (n/N), together with the corresponding percentage. ESM scores provide a sequence-based estimate of predicted functional impact, with more negative scores indicating lower compatibility with the sequence constraints of the respective protein. Missense substitutions were classified as predicted high impact when their ESM score fell below the protein-specific threshold derived from a two-component Gaussian mixture model. For stop-gained variants, ESM scores were assigned using the lowest score preceding the introduced stop codon. An ESM score of 0 indicates variants for which reconstruction resulted in no change to the encoded protein sequence. ESM scores are left blank for frameshift variants. Complex/multiresidue and in-frame insertion/deletion variants were excluded from the lollipop analysis (**Fig. 4c**) to restrict the comparison to consistently interpretable missense, stop-gained and frameshift variants. Variants occurring in ≥95% of the epidemiological isolates within a clade were also excluded, as these were considered clade-associated background variation rather than within-clade epidemiological variation.

| Gene | Variant | DNA change | Protein position (a.a.) | Protein change | Consequence | Clade occurrence (n/N; %) | ESM score | Predicted high impact | Lollipop inclusion |
|---|---|---|---|---|---|---|---|---|---|
| *SLN1* | 40:450,185 | TTTC>CTTT | 29 | Complex | Multiresidue | V: 3/3 (100.00%) | 0 | | No – complex/in-frame |
| *SLN1* | 40:450,226 | AT>A | 43 | Del-Frameshift-n=1_43 | Frameshift | IV: 1/1576 (0.06%) | | | Yes |
| *SLN1* | 40:450,232 | C>G | 45 | T45S | Missense | V: 3/3 (100.00%) | -2.54323283 | | No – ≥95% within clade |
| *SLN1* | 40:450,262 | T>C | 55 | L55S | Missense | IV: 1/1576 (0.06%) | -9.51491182 | High impact | Yes |
| *SLN1* | 40:450,278 | AATA>GATT | 60 | Complex | Multiresidue | V: 3/3 (100.00%) | 0 | | No – complex/in-frame |
| *SLN1* | 40:450,297 | GAGCAAA>CAACAGT | 66 | Complex | Multiresidue | V: 3/3 (100.00%) | 0.46166992 | | No – complex/in-frame |
| *SLN1* | 40:450,308 | GCAA>CCAG | 70 | Complex | Multiresidue | V: 3/3 (100.00%) | 0 | | No – complex/in-frame |
| *SLN1* | 40:450,326 | GACGTAT>AACATAC | 76 | Complex | Multiresidue | V: 3/3 (100.00%) | 0 | | No – complex/in-frame |
| *SLN1* | 40:450,353 | TATTG>CATCA | 85 | Complex | Multiresidue | V: 3/3 (100.00%) | -0.15014648 | | No – complex/in-frame |
| *SLN1* | 40:450,365 | TCTCTCGAATT>CCTTTCAAATC | 89 | Complex | Multiresidue | V: 3/3 (100.00%) | -2.4179077 | | No – complex/in-frame |
| *SLN1* | 40:450,397 | A>G | 100 | K100R | Missense | V: 3/3 (100.00%) | -3.58500981 | | No – ≥95% within clade |
| *SLN1* | 40:450,402 | G>A | 102 | V102I | Missense | V: 3/3 (100.00%) | -3.76632157 | | No – ≥95% within clade |
| *SLN1* | 40:450,410 | CGACGCC>AGAAGCA | 104 | Complex | Multiresidue | V: 3/3 (100.00%) | -0.07489014 | | No – complex/in-frame |
| *SLN1* | 40:450,497 | CGCC>GGCT | 133 | Complex | Multiresidue | V: 3/3 (100.00%) | 0 | | No – complex/in-frame |
| *SLN1* | 40:450,506 | TCG>ACA | 136 | Complex | Multiresidue | V: 3/3 (100.00%) | -0.7761841 | | No – complex/in-frame |
| *SLN1* | 40:450,524 | GGTT>CGTC | 142 | Complex | Multiresidue | V: 3/3 (100.00%) | 0.67471313 | | No – complex/in-frame |
| *SLN1* | 40:450,536 | TTTGGAG>CTTAGAT | 146 | Complex | Multiresidue | V: 2/3 (66.67%) | 0.3840332 | | No – complex/in-frame |
| *SLN1* | 40:450,547 | C>T | 150 | A150V | Missense | V: 2/3 (66.67%) | -3.20631164 | | Yes |
| *SLN1* | 40:450,561 | A>G | 155 | T155A | Missense | V: 2/3 (66.67%) | -3.47786692 | | Yes |
| *SLN1* | 40:450,661 | C>T | 188 | P188L | Missense | IV: 13/1576 (0.82%) | -2.03972661 | | Yes |
| *SLN1* | 40:450,674 | CAA>TAC | 192 | Complex | Multiresidue | VI: 2/6 (33.33%) | 0.22998047 | | No – complex/in-frame |

| Gene | Variant | DNA change | Protein position (a.a.) | Protein change | Consequence | Clade occurrence (n/N; %) | ESM score | Predicted high impact | Lollipop inclusion |
|---|---|---|---|---|---|---|---|---|---|
| *SLN1* | 40:450,676 | A>C | 193 | K193T | Missense | II: 40/40 (100.00%); III: 5502/5568 (98.81%); IV: 1576/1576 (100.00%); V: 3/3 (100.00%); VI: 4/6 (66.67%) | -1.86589253 | | Yes |
| *SLN1* | 40:450,719 | AAAT>GAAC | 207 | Complex | Multiresidue | V: 3/3 (100.00%) | 0 | | No – complex/in-frame |
| *SLN1* | 40:450,777 | G>A | 227 | D227N | Missense | V: 3/3 (100.00%) | -2.64209458 | | No – ≥95% within clade |
| *SLN1* | 40:450,782 | TGA>CGG | 228 | Complex | Multiresidue | V: 3/3 (100.00%) | -1.7591248 | | No – complex/in-frame |
| *SLN1* | 40:450,791 | TAAACTA>CAAAACG | 231 | Complex | Multiresidue | V: 3/3 (100.00%) | 2.2997437 | | No – complex/in-frame |
| *SLN1* | 40:450,809 | TTTCGG>CTACGA | 237 | Complex | Multiresidue | V: 3/3 (100.00%) | -0.522583 | | No – complex/in-frame |
| *SLN1* | 40:450,813 | G>A | 239 | G239R | Missense | I: 3/5174 (0.06%) | -4.5725645 | | Yes |
| *SLN1* | 40:450,837 | T>C | 247 | S247P | Missense | IV: 2/1576 (0.13%) | -6.16838886 | | Yes |
| *SLN1* | 40:450,842 | CTCTCTC>GTCCCTT | 248 | Complex | Multiresidue | V: 3/3 (100.00%) | 0 | | No – complex/in-frame |
| *SLN1* | 40:450,855 | G>A | 253 | A253T | Missense | I: 1/5174 (0.02%) | -3.58217476 | | Yes |
| *SLN1* | 40:450,870 | G>T | 258 | A258S | Missense | V: 3/3 (100.00%) | -3.30084851 | | No – ≥95% within clade |
| *SLN1* | 40:450,881 | AGGC>TGGA | 261 | Complex | Multiresidue | V: 3/3 (100.00%) | 0 | | No – complex/in-frame |
| *SLN1* | 40:450,899 | TACTGGA>AACCGGC | 267 | Complex | Multiresidue | V: 3/3 (100.00%) | 0 | | No – complex/in-frame |
| *SLN1* | 40:450,915 | TTGTTCCCA>CTATTTCCC | 272 | Complex | Multiresidue | V: 3/3 (100.00%) | 0 | | No – complex/in-frame |
| *SLN1* | 40:450,937 | A>ATCG | 280 | Ins-Non-frameshift-n=3_280 | In-frame indel | V: 3/3 (100.00%) | -0.44369507 | | No – complex/in-frame |
| *SLN1* | 40:450,942 | ATT>GTA | 281 | Complex | Multiresidue | V: 3/3 (100.00%) | 0.06378174 | | No – complex/in-frame |
| *SLN1* | 40:450,965 | AGAC>TGAT | 289 | Complex | Multiresidue | V: 3/3 (100.00%) | 0 | | No – complex/in-frame |
| *SLN1* | 40:450,980 | AGTT>CATA | 294 | Complex | Multiresidue | V: 3/3 (100.00%) | -0.04125977 | | No – complex/in-frame |
| *SLN1* | 40:451,064 | TACT>AACC | 322 | Complex | Multiresidue | V: 3/3 (100.00%) | 0 | | No – complex/in-frame |
| *SLN1* | 40:451,073 | T>A | 325 | D325E | Missense | V: 3/3 (100.00%) | -3.67095332 | | No – ≥95% within clade |
| *SLN1* | 40:451,080 | TT>CC | 327 | Complex | Multiresidue | V: 3/3 (100.00%) | -2.4849854 | | No – complex/in-frame |
| *SLN1* | 40:451,114 | AG>GA | 339 | Complex | Multiresidue | V: 3/3 (100.00%) | -0.49554443 | | No – complex/in-frame |
| *SLN1* | 40:451,127 | CCAGA>TCAAG | 343 | Complex | Multiresidue | V: 3/3 (100.00%) | -0.15692139 | | No – complex/in-frame |
| *SLN1* | 40:451,165 | G>A | 356 | G356E | Missense | I: 1/5174 (0.02%) | -11.97682927 | High impact | Yes |
| *SLN1* | 40:451,205 | ATTCCCT>TTTTCCC | 369 | Complex | Multiresidue | V: 3/3 (100.00%) | 0 | | No – complex/in-frame |
| *SLN1* | 40:451,229 | CGCTCCC>TGCACCT | 377 | Complex | Multiresidue | V: 3/3 (100.00%) | 0 | | No – complex/in-frame |
| *SLN1* | 40:451,256 | GACTGAG>TACGGAA | 386 | Complex | Multiresidue | V: 3/3 (100.00%) | 0 | | No – complex/in-frame |
| *SLN1* | 40:451,294 | A>G | 399 | K399R | Missense | V: 3/3 (100.00%) | -2.4347611 | | No – ≥95% within clade |
| *SLN1* | 40:451,303 | GCTTTT>ATCTAC | 402 | Complex | Multiresidue | V: 3/3 (100.00%) | -0.31604004 | | No – complex/in-frame |
| *SLN1* | 40:451,321 | G>A | 408 | G408E | Missense | II: 40/40 (100.00%) | -4.47598056 | | No – ≥95% within clade |
| *SLN1* | 40:451,324 | GTCACGA>CTTACGG | 409 | Complex | Multiresidue | V: 3/3 (100.00%) | -0.17098999 | | No – complex/in-frame |
| *SLN1* | 40:451,366 | T>C | 423 | V423A | Missense | V: 3/3 (100.00%) | -2.63683546 | | No – ≥95% within clade |
| *SLN1* | 40:451,429 | AC>GT | 444 | Complex | Multiresidue | V: 3/3 (100.00%) | 0.380188 | | No – complex/in-frame |
| *SLN1* | 40:451,493 | TCTC>ACTT | 465 | Complex | Multiresidue | V: 3/3 (100.00%) | 0 | | No – complex/in-frame |
| *SLN1* | 40:451,518 | A>T | 474 | K474* | Stop-gained | I: 1/5174 (0.02%) | -25.97406521 | | Yes |
| *SLN1* | 40:451,544 | AGAACTT>TGAGCTC | 482 | Complex | Multiresidue | V: 3/3 (100.00%) | 0 | | No – complex/in-frame |
| *SLN1* | 40:451,586 | GAAG>AAAA | 496 | Complex | Multiresidue | V: 3/3 (100.00%) | 0 | | No – complex/in-frame |
| *SLN1* | 40:451,619 | GC>AA | 507 | Complex | Multiresidue | V: 3/3 (100.00%) | 0.7112732 | | No – complex/in-frame |
| *SLN1* | 40:451,667 | AAATGAGTCA>GAACGAATCG | 523 | Complex | Multiresidue | V: 3/3 (100.00%) | 0 | | No – complex/in-frame |

| Gene | Variant | DNA change | Protein position (a.a.) | Protein change | Consequence | Clade occurrence (n/N; %) | ESM score | Predicted high impact | Lollipop inclusion |
|---|---|---|---|---|---|---|---|---|---|
| *SLN1* | 40:451,682 | TATA>CATT | 528 | Complex | Multiresidue | V: 3/3 (100.00%) | 0 | | No – complex/in-frame |
| *SLN1* | 40:451,727 | TGGCATC>CGGTATT | 543 | Complex | Multiresidue | V: 3/3 (100.00%) | 0 | | No – complex/in-frame |
| *SLN1* | 40:451,784 | T>G | 562 | H562Q | Missense | I: 3/5174 (0.06%) | 0.92033052 | | Yes |
| *SLN1* | 40:451,853 | CTCC>TTCT | 585 | Complex | Multiresidue | V: 3/3 (100.00%) | 0 | | No – complex/in-frame |
| *SLN1* | 40:451,859 | G>C | 587 | K587N | Missense | III: 1/5568 (0.02%) | -12.99181532 | High impact | Yes |
| *SLN1* | 40:451,967 | T>G | 623 | N623K | Missense | V: 3/3 (100.00%) | -4.43710354 | | No – ≥95% within clade |
| *SLN1* | 40:452,051 | CCTC>TCTT | 651 | Complex | Multiresidue | V: 3/3 (100.00%) | 0 | | No – complex/in-frame |
| *SLN1* | 40:452,176 | A>G | 693 | K693R | Missense | II: 40/40 (100.00%); III: 5506/5568 (98.89%); IV: 1574/1576 (99.87%); V: 3/3 (100.00%); VI: 6/6 (100.00%) | -4.16863711 | | No – ≥95% within clade |
| *SLN1* | 40:452,204 | AA>GT | 702 | Complex | Multiresidue | V: 3/3 (100.00%) | 0.5670471 | | No – complex/in-frame |
| *SLN1* | 40:452,225 | AT>GC | 709 | Complex | Multiresidue | V: 3/3 (100.00%) | -0.00323486 | | No – complex/in-frame |
| *SLN1* | 40:452,226 | T>C | 710 | Y710H | Missense | II: 40/40 (100.00%); III: 5506/5568 (98.89%); IV: 1576/1576 (100.00%); VI: 6/6 (100.00%) | -3.3594816 | | No – ≥95% within clade |
| *SLN1* | 40:452,317 | CAAG>TAGT | 740 | Complex | Multiresidue | V: 3/3 (100.00%) | 0.79870605 | | No – complex/in-frame |
| *SLN1* | 40:452,325 | G>A | 743 | D743N | Missense | V: 3/3 (100.00%) | -3.06232044 | | No – ≥95% within clade |
| *SLN1* | 40:452,332 | A>C | 745 | D745A | Missense | V: 3/3 (100.00%) | -3.12797141 | | No – ≥95% within clade |
| *SLN1* | 40:452,337 | T>C | 747 | S747P | Missense | V: 3/3 (100.00%) | -4.5143862 | | No – ≥95% within clade |
| *SLN1* | 40:452,347 | G>A | 750 | S750N | Missense | I: 5/5174 (0.10%) | -3.69065621 | | Yes |
| *SLN1* | 40:452,415 | A>AT | 772 | Ins-Frameshift-n=1_772 | Frameshift | III: 1/5568 (0.02%) | | | Yes |
| *SLN1* | 40:452,440 | ATAAGTCGT>A | 781 | Del-Frameshift-n=8_781 | Frameshift | IV: 1/1576 (0.06%) | | | Yes |
| *SLN1* | 40:452,457 | T>A | 787 | L787I | Missense | IV: 1573/1576 (99.81%); V: 3/3 (100.00%); VI: 6/6 (100.00%) | -3.687029 | | No – ≥95% within clade |
| *SLN1* | 40:452,478 | T>G | 794 | L794V | Missense | I: 5/5174 (0.10%) | -4.26494728 | | Yes |
| *SLN1* | 40:452,537 | CGCC>TGCT | 813 | Complex | Multiresidue | V: 3/3 (100.00%) | 0 | | No – complex/in-frame |
| *SLN1* | 40:452,539 | C>T | 814 | A814V | Missense | II: 40/40 (100.00%) | -3.61580775 | | No – ≥95% within clade |
| *SLN1* | 40:452,545 | T>A | 816 | V816D | Missense | IV: 1575/1576 (99.94%); V: 3/3 (100.00%); VI: 6/6 (100.00%) | -3.22533051 | | No – ≥95% within clade |
| *SLN1* | 40:452,554 | A>C | 819 | N819T | Missense | V: 3/3 (100.00%) | -3.28294381 | | No – ≥95% within clade |
| *SLN1* | 40:452,554 | A>T | 819 | N819I | Missense | IV: 1572/1576 (99.75%); VI: 6/6 (100.00%) | -3.91378475 | | No – ≥95% within clade |
| *SLN1* | 40:452,595 | A>G | 833 | T833A | Missense | II: 40/40 (100.00%) | -3.12192962 | | No – ≥95% within clade |
| *SLN1* | 40:452,624 | CA>TT | 842 | Complex | Multiresidue | IV: 1573/1576 (99.81%); VI: 6/6 (100.00%) | -0.9225464 | | No – complex/in-frame |
| *SLN1* | 40:452,629 | C>T | 844 | T844I | Missense | V: 3/3 (100.00%) | -3.69820133 | | No – ≥95% within clade |
| *SLN1* | 40:452,645 | TACT>AAAC | 849 | Complex | Multiresidue | V: 3/3 (100.00%) | 0.23275757 | | No – complex/in-frame |
| *SLN1* | 40:452,673 | G>A | 859 | A859T | Missense | V: 3/3 (100.00%) | -3.05891509 | | No – ≥95% within clade |
| *SLN1* | 40:452,686 | A>T | 863 | Q863L | Missense | III: 5496/5568 (98.71%) | -2.72994806 | | No – ≥95% within clade |
| *SLN1* | 40:452,735 | GTCGACA>G | 879 | Del-Non-frameshift-n=6_879 | In-frame indel | V: 3/3 (100.00%) | -0.1513977 | | No – complex/in-frame |
| *SLN1* | 40:452,856 | G>A | 920 | G920S | Missense | IV: 78/1576 (4.95%) | -12.30746254 | High impact | Yes |
| *SLN1* | 40:452,992 | TG>T | 965 | Del-Frameshift-n=1_965 | Frameshift | I: 2/5174 (0.04%) | | | Yes |
| *SLN1* | 40:453,000 | AAG>C | 967 | Del-Frameshift-n=2_967 | Frameshift | IV: 1/1576 (0.06%) | | | Yes |

| Gene | Variant | DNA change | Protein position (a.a.) | Protein change | Consequence | Clade occurrence (n/N; %) | ESM score | Predicted high impact | Lollipop inclusion |
|---|---|---|---|---|---|---|---|---|---|
| *SLN1* | 40:453,066 | C>G | 990 | L990V | Missense | V: 3/3 (100.00%) | -3.20883496 | | No – ≥95% within clade |
| *SLN1* | 40:453,092 | G>C | 998 | E998D | Missense | IV: 1572/1576 (99.75%); VI: 6/6 (100.00%) | -3.80031559 | | No – ≥95% within clade |
| *SLN1* | 40:453,096 | TGCA>T | 999 | Del-Non-frameshift-n=3_999 | In-frame indel | III: 2/5568 (0.04%) | -0.3947754 | | No – complex/in-frame |
| *SLN1* | 40:453,119 | G>T | 1007 | E1007D | Missense | V: 3/3 (100.00%) | -3.16284096 | | No – ≥95% within clade |
| *SLN1* | 40:453,150 | G>T | 1018 | V1018L | Missense | V: 3/3 (100.00%) | -3.14164723 | | No – ≥95% within clade |
| *SLN1* | 40:453,166 | A>C | 1023 | H1023P | Missense | V: 3/3 (100.00%) | -1.82962853 | | No – ≥95% within clade |
| *SLN1* | 40:453,169 | C>T | 1024 | S1024L | Missense | I: 5126/5174 (99.07%) | -4.95138512 | | No – ≥95% within clade |
| *SLN1* | 40:453,181 | C>T | 1028 | S1028L | Missense | V: 3/3 (100.00%) | -6.11760366 | | No – ≥95% within clade |
| *SLN1* | 40:453,258 | GCCTCTCCG>TCCTCTCCA | 1053 | Complex | Multiresidue | IV: 1/1576 (0.06%) | 0.15927124 | | No – complex/in-frame |
| *SLN1* | 40:453,258 | G>T | 1054 | A1054S | Missense | IV: 1574/1576 (99.87%); V: 3/3 (100.00%); VI: 6/6 (100.00%) | -2.58032969 | | No – ≥95% within clade |
| *SLN1* | 40:453,382 | A>G | 1095 | K1095R | Missense | V: 3/3 (100.00%) | -4.33765959 | | No – ≥95% within clade |
| *SLN1* | 40:453,549 | GTGC>T | 1150 | Del-Non-frameshift-n=3_1150 | In-frame indel | IV: 26/1576 (1.65%) | 21.605042 | | No – complex/in-frame |
| *SLN1* | 40:453,657 | G>C | 1187 | V1187L | Missense | II: 13/40 (32.50%) | -8.1493374 | High impact | Yes |
| *SLN1* | 40:453,703 | C>G | 1202 | P1202R | Missense | III: 1/5568 (0.02%) | -17.98300064 | High impact | Yes |
| *SLN1* | 40:453,706 | T>C | 1203 | I1203T | Missense | I: 1/5174 (0.02%) | -8.36134717 | High impact | Yes |
| *NIK1* | 40:39,960 | C>T | 75 | S75L | Missense | II: 13/40 (32.50%) | -10.10753283 | High impact | Yes |
| *NIK1* | 40:40,473 | G>A | 246 | R246K | Missense | IV: 1/1576 (0.06%) | -9.57542673 | High impact | Yes |
| *NIK1* | 40:40,514 | CTATCC>TTGTCT | 259 | Complex | Multiresidue | V: 3/3 (100.00%) | 0 | | No – complex/in-frame |
| *NIK1* | 40:40,539 | G>A | 268 | C268Y | Missense | III: 1/5568 (0.02%) | -8.83899402 | | Yes |
| *NIK1* | 40:40,632 | AGGTC>A | 299 | Del-Frameshift-n=4_299 | Frameshift | I: 6/5174 (0.12%) | | | Yes |
| *NIK1* | 40:40,752 | C>A | 339 | S339Y | Missense | III: 1/5568 (0.02%) | -7.31325307 | | Yes |
| *NIK1* | 40:41,067 | T>C | 444 | F444S | Missense | IV: 1/1576 (0.06%) | -12.00377122 | High impact | Yes |
| *NIK1* | 40:41,108 | G>A | 458 | A458T | Missense | IV: 1/1576 (0.06%) | -3.44632477 | | Yes |
| *NIK1* | 40:41,130 | A>T | 465 | Q465L | Missense | III: 1/5568 (0.02%) | -7.55933167 | | Yes |
| *NIK1* | 40:41,135 | G>A | 467 | V467M | Missense | III: 1/5568 (0.02%) | -8.38809311 | | Yes |
| *NIK1* | 40:41,266 | TGGA>CGGG | 510 | Complex | Multiresidue | V: 3/3 (100.00%) | 0 | | No – complex/in-frame |
| *NIK1* | 40:41,358 | C>A | 541 | T541K | Missense | III: 2/5568 (0.04%) | -9.37008018 | High impact | Yes |
| *NIK1* | 40:41,664 | A>C | 643 | K643T | Missense | III: 2/5568 (0.04%) | -3.57049941 | | Yes |
| *NIK1* | 40:41,798 | T>C | 688 | S688P | Missense | IV: 1/1576 (0.06%) | -10.7629187 | High impact | Yes |
| *NIK1* | 40:41,951 | C>T | 739 | H739Y | Missense | IV: 1/1576 (0.06%) | -5.0167965 | | Yes |
| *NIK1* | 40:42,006 | G>A | 757 | R757K | Missense | IV: 5/1576 (0.32%) | -2.72945686 | | Yes |
| *NIK1* | 40:42,207 | G>T | 824 | C824F | Missense | IV: 1/1576 (0.06%) | -6.17392555 | | Yes |
| *NIK1* | 40:42,384 | T>A | 883 | I883N | Missense | III: 2/5568 (0.04%) | -10.56647775 | High impact | Yes |
| *NIK1* | 40:42,887 | G>A | 1051 | A1051T | Missense | IV: 1/1576 (0.06%) | -2.73140091 | | Yes |
| *NIK1* | 40:42,953 | C>G | 1073 | R1073G | Missense | I: 1/5174 (0.02%) | -3.41381577 | | Yes |
| *SSK1* | 41:1,232,427 | G>A | 7 | Q7* | Stop-gained | I: 1/5174 (0.02%) | -24.056969 | | Yes |
| *SSK1* | 41:1,232,324 | TTACGGACCCATA>T | 41 | Del-Non-frameshift-n=12_41 | In-frame indel | IV: 1/1576 (0.06%) | 0.4230652 | | No – complex/in-frame |
| *SSK1* | 41:1,232,260 | CAATT>C | 62 | Del-Frameshift-n=4_62 | Frameshift | I: 1/5174 (0.02%) | | | Yes |
| *SSK1* | 41:1,232,185 | AAT>TAC | 87 | Complex | Multiresidue | V: 3/3 (100.00%) | -0.06561279 | | No – complex/in-frame |
| *SSK1* | 41:1,232,084 | G>A | 121 | P121L | Missense | I: 1/5174 (0.02%) | -3.967769 | | Yes |

| Gene | Variant | DNA change | Protein position (a.a.) | Protein change | Consequence | Clade occurrence (n/N; %) | ESM score | Predicted high impact | Lollipop inclusion |
|---|---|---|---|---|---|---|---|---|---|
| *SSK1* | 41:1,232,066 | C>T | 127 | S127N | Missense | IV: 1575/1576 (99.94%); VI: 6/6 (100.00%) | -4.262076 | | No – ≥95% within clade |
| *SSK1* | 41:1,231,914 | C>T | 178 | G178S | Missense | IV: 1573/1576 (99.81%); VI: 6/6 (100.00%) | -2.4097664 | | No – ≥95% within clade |
| *SSK1* | 41:1,231,830 | T>G | 206 | N206H | Missense | V: 3/3 (100.00%) | -3.34463 | | No – ≥95% within clade |
| *SSK1* | 41:1,231,821 | T>C | 209 | T209A | Missense | V: 3/3 (100.00%) | -3.365625 | | No – ≥95% within clade |
| *SSK1* | 41:1,231,775 | G>T | 224 | T224N | Missense | IV: 1574/1576 (99.87%); V: 3/3 (100.00%); VI: 6/6 (100.00%) | -3.118393 | | No – ≥95% within clade |
| *SSK1* | 41:1,231,734 | GGT>AGC | 237 | Complex | Multiresidue | V: 3/3 (100.00%) | 0.14506531 | | No – complex/in-frame |
| *SSK1* | 41:1,231,731 | TTGGGTTCATC>T | 238 | Del-Frameshift-n=10_238 | Frameshift | IV: 1/1576 (0.06%) | | | Yes |
| *SSK1* | 41:1,231,703 | G>T | 248 | S248* | Stop-gained | I: 1/5174 (0.02%) | -24.056969 | | Yes |
| *SSK1* | 41:1,231,666 | GCCGTTG>CCCGTTA | 260 | Complex | Multiresidue | V: 1/3 (33.33%) | 0 | | No – complex/in-frame |
| *SSK1* | 41:1,231,603 | GATA>AATG | 281 | Complex | Multiresidue | V: 3/3 (100.00%) | 0 | | No – complex/in-frame |
| *SSK1* | 41:1,231,414 | GT>AG | 344 | Complex | Multiresidue | V: 3/3 (100.00%) | -0.11047363 | | No – complex/in-frame |
| *SSK1* | 41:1,231,276 | G>T | 390 | S390R | Missense | III: 5487/5568 (98.55%) | -3.383934 | | No – ≥95% within clade |
| *SSK1* | 41:1,231,270 | GC>CG | 392 | Complex | Multiresidue | V: 3/3 (100.00%) | -0.0710907 | | No – complex/in-frame |
| *SSK1* | 41:1,231,198 | ATCAC>GTCAT | 416 | Complex | Multiresidue | V: 3/3 (100.00%) | -0.00019836 | | No – complex/in-frame |
| *SSK1* | 41:1,231,182 | C>T | 422 | V422M | Missense | II: 40/40 (100.00%) | -4.793709 | | No – ≥95% within clade |
| *SSK1* | 41:1,231,142 | A>C | 435 | L435R | Missense | I: 1/5174 (0.02%) | -12.56689 | High impact | Yes |
| *SSK1* | 41:1,231,130 | G>A | 439 | S439L | Missense | V: 3/3 (100.00%) | -10.925125 | High impact | No – ≥95% within clade |
| *SSK1* | 41:1,231,068 | GTT>G | 459 | Del-Frameshift-n=2_459 | Frameshift | III: 2/5568 (0.04%) | | | Yes |
| *SSK1* | 41:1,230,845 | C>T | 534 | R534K | Missense | II: 40/40 (100.00%); IV: 1575/1576 (99.94%); V: 3/3 (100.00%); VI: 6/6 (100.00%) | -1.9217405 | | No – ≥95% within clade |
| *SSK1* | 41:1,230,825 | G>A | 541 | R541C | Missense | IV: 8/1576 (0.51%) | -8.382664 | High impact | Yes |
| *SSK1* | 41:1,230,817 | CGGA>TGGT | 543 | Complex | Multiresidue | V: 3/3 (100.00%) | 0 | | No – complex/in-frame |
| *SSK1* | 41:1,230,806 | A>G | 547 | V547A | Missense | IV: 1/1576 (0.06%) | -11.759764 | High impact | Yes |
| *SSK1* | 41:1,230,681 | A>AC | 588 | Ins-Frameshift-n=1_588 | Frameshift | III: 1/5568 (0.02%) | | | Yes |
| *SSK1* | 41:1,230,668 | A>G | 593 | L593P | Missense | IV: 1/1576 (0.06%) | -13.535643 | High impact | Yes |
| *SSK1* | 41:1,230,628 | G>C | 606 | N606K | Missense | II: 40/40 (100.00%); IV: 1575/1576 (99.94%); V: 3/3 (100.00%); VI: 6/6 (100.00%) | -3.5441828 | | No – ≥95% within clade |
| *SSK1* | 41:1,230,578 | A>G | 623 | I623T | Missense | V: 3/3 (100.00%) | -3.1464236 | | No – ≥95% within clade |
| *SSK1* | 41:1,230,564 | G>A | 628 | P628S | Missense | V: 3/3 (100.00%) | -2.6287313 | | No – ≥95% within clade |
| *SSK2* | 41:676,914 | A>T | 44 | S44C | Missense | I: 1/5174 (0.02%) | -10.52990747 | High impact | Yes |
| *SSK2* | 41:676,942 | A>C | 53 | N53T | Missense | II: 1/40 (2.50%) | -2.86008888 | | Yes |
| *SSK2* | 41:677,131 | GG>AA | 116 | Complex | Multiresidue | IV: 1572/1576 (99.75%); VI: 6/6 (100.00%) | 0.00018311 | | No – complex/in-frame |
| *SSK2* | 41:677,246 | CGAT>C | 154 | Del-Non-frameshift-n=3_154 | In-frame indel | I: 38/5174 (0.73%) | 1.0057373 | | No – complex/in-frame |
| *SSK2* | 41:677,410 | A>G | 209 | D209G | Missense | IV: 1/1576 (0.06%) | -5.54380633 | | Yes |
| *SSK2* | 41:677,419 | T>A | 212 | V212D | Missense | III: 2/5568 (0.04%) | -3.29905921 | | Yes |
| *SSK2* | 41:677,618 | G>T | 278 | K278N | Missense | II: 2/40 (5.00%) | -3.50458807 | | Yes |
| *SSK2* | 41:677,622 | A>C | 280 | I280L | Missense | III: 1/5568 (0.02%) | -2.75947529 | | Yes |
| *SSK2* | 41:678,113 | G>A | 443 | W443* | Stop-gained | I: 1/5174 (0.02%) | -24.02511266 | | Yes |

| Gene | Variant | DNA change | Protein position (a.a.) | Protein change | Consequence | Clade occurrence (n/N; %) | ESM score | Predicted high impact | Lollipop inclusion |
|---|---|---|---|---|---|---|---|---|---|
| *SSK2* | 41:678,119 | CAAGTCAAAGGAAACTT>C | 445 | Del-Frameshift-n=16_445 | Frameshift | III: 1/5568 (0.02%) | | | Yes |
| *SSK2* | 41:678,142 | C>A | 453 | T453K | Missense | IV: 1572/1576 (99.75%); V: 3/3 (100.00%); VI: 6/6 (100.00%) | -2.26643947 | | No – ≥95% within clade |
| *SSK2* | 41:678,480 | C>T | 566 | R566* | Stop-gained | III: 1/5568 (0.02%) | -24.02511266 | | Yes |
| *SSK2* | 41:678,571 | A>AT | 596 | Ins-Frameshift-n=1_596 | Frameshift | I: 1/5174 (0.02%) | | | Yes |
| *SSK2* | 41:678,834 | A>T | 684 | N684Y | Missense | I: 21/5174 (0.41%) | -3.97849265 | | Yes |
| *SSK2* | 41:678,903 | G>A | 707 | G707R | Missense | IV: 1/1576 (0.06%) | -2.41648909 | | Yes |
| *SSK2* | 41:679,166 | CAATATAAAGGAAGTGCCG>C | 794 | Del-Non-frameshift-n=18_794 | In-frame indel | III: 2/5568 (0.04%) | -3.2091675 | | No – complex/in-frame |
| *SSK2* | 41:679,458 | C>T | 892 | Q892* | Stop-gained | III: 1/5568 (0.02%) | -24.02511266 | | Yes |
| *SSK2* | 41:679,489 | C>T | 902 | A902V | Missense | I: 2/5174 (0.04%) | -5.2936299 | | Yes |
| *SSK2* | 41:679,636 | G>A | 951 | C951Y | Missense | III: 1/5568 (0.02%) | -7.78592434 | High impact | Yes |
| *SSK2* | 41:679,766 | CA>C | 994 | Del-Frameshift-n=1_994 | Frameshift | III: 3/5568 (0.05%) | | | Yes |
| *SSK2* | 41:679,776 | G>A | 998 | A998T | Missense | I: 5/5174 (0.10%) | -4.93999991 | | Yes |
| *SSK2* | 41:679,788 | G>A | 1002 | E1002K | Missense | I: 5/5174 (0.10%) | -6.19895829 | | Yes |
| *SSK2* | 41:679,969 | T>C | 1062 | V1062A | Missense | I: 1/5174 (0.02%) | -2.67942357 | | Yes |
| *SSK2* | 41:680,059 | G>A | 1092 | G1092E | Missense | I: 2/5174 (0.04%) | -5.35734153 | | Yes |
| *SSK2* | 41:680,090 | TA>T | 1102 | Del-Frameshift-n=1_1102 | Frameshift | I: 1/5174 (0.02%) | | | Yes |
| *SSK2* | 41:680,345 | T>A | 1187 | Y1187* | Stop-gained | I: 4/5174 (0.08%) | -24.02511266 | | Yes |
| *SSK2* | 41:680,457 | G>A | 1225 | D1225N | Missense | I: 2/5174 (0.04%) | -11.93070364 | High impact | Yes |
| *SSK2* | 41:680,469 | G>C | 1229 | A1229P | Missense | III: 2/5568 (0.04%) | -17.17819774 | High impact | Yes |
| *SSK2* | 41:680,554 | AT>A | 1257 | Del-Frameshift-n=1_1257 | Frameshift | III: 3/5568 (0.05%) | | | Yes |
| *SSK2* | 41:680,631 | G>C | 1283 | D1283H | Missense | IV: 1/1576 (0.06%) | -16.93025255 | High impact | Yes |
| *SSK2* | 41:680,693 | TT>CG | 1303 | Complex | Multiresidue | IV: 1/1576 (0.06%) | -1.5760193 | | No – complex/in-frame |
| *SSK2* | 41:680,818 | C>T | 1345 | A1345V | Missense | IV: 1573/1576 (99.81%); V: 3/3 (100.00%); VI: 6/6 (100.00%) | -4.45569469 | | No – ≥95% within clade |
| *SSK2* | 41:680,871 | C>G | 1363 | R1363G | Missense | IV: 2/1576 (0.13%) | -6.44129445 | | Yes |
| *SSK2* | 41:680,875 | AAG>A | 1364 | Del-Frameshift-n=2_1364 | Frameshift | IV: 1/1576 (0.06%) | | | Yes |
| *SSK2* | 41:680,881 | C>T | 1366 | A1366V | Missense | I: 1/5174 (0.02%) | -3.78788263 | | Yes |
| *PBS2* | 38:626,683 | G>C | 42 | D42E | Missense | IV: 1575/1576 (99.94%); V: 3/3 (100.00%); VI: 6/6 (100.00%) | -3.658356 | | No – ≥95% within clade |
| *PBS2* | 38:626,669 | A>G | 47 | I47T | Missense | VI: 6/6 (100.00%) | -2.2900991 | | No – ≥95% within clade |
| *PBS2* | 38:626,471 | G>A | 113 | P113L | Missense | IV: 1/1576 (0.06%) | -6.5203695 | | Yes |
| *PBS2* | 38:626,137 | C>A | 224 | K224N | Missense | IV: 1/1576 (0.06%) | -15.081397 | High impact | Yes |
| *PBS2* | 38:626,046 | C>T | 255 | D255N | Missense | III: 1/5568 (0.02%) | -8.580128 | | Yes |
| *PBS2* | 38:626,019 | C>A | 264 | G264C | Missense | III: 38/5568 (0.68%) | -10.536579 | High impact | Yes |
| *PBS2* | 38:625,986 | C>A | 275 | G275* | Stop-gained | I: 2/5174 (0.04%) | -23.299772 | | Yes |
| *PBS2* | 38:625,863 | C>T | 316 | D316N | Missense | IV: 1/1576 (0.06%) | -14.23076 | High impact | Yes |
| *PBS2* | 38:625,766 | G>T | 348 | T348N | Missense | III: 4/5568 (0.07%) | -10.606321 | High impact | Yes |
| *PBS2* | 38:625,748 | G>A | 354 | S354F | Missense | I: 2/5174 (0.04%) | -14.061874 | High impact | Yes |
| *PBS2* | 38:625,736 | G>A | 358 | P358L | Missense | I: 1/5174 (0.02%) | -12.491153 | High impact | Yes |
| *PBS2* | 38:625,619 | T>A | 397 | E397V | Missense | III: 1/5568 (0.02%) | -9.574102 | High impact | Yes |
| *PBS2* | 38:625,530 | G>A | 427 | Q427* | Stop-gained | III: 1/5568 (0.02%) | -18.522865 | | Yes |

| Gene | Variant | DNA change | Protein position (a.a.) | Protein change | Consequence | Clade occurrence (n/N; %) | ESM score | Predicted high impact | Lollipop inclusion |
|---|---|---|---|---|---|---|---|---|---|
| *PBS2* | 38:625,497 | G>T | 438 | P438T | Missense | III: 1/5568 (0.02%) | -8.206857 | | Yes |
| *PBS2* | 38:625,481 | G>T | 443 | S443* | Stop-gained | III: 1/5568 (0.02%) | -18.469265 | | Yes |
| *PBS2* | 38:625,394 | C>T | 472 | R472K | Missense | IV: 1575/1576 (99.94%); V: 3/3 (100.00%); VI: 6/6 (100.00%) | -2.5289192 | | No – ≥95% within clade |
| *PBS2* | 38:625,353 | A>C | 486 | S486A | Missense | V: 3/3 (100.00%) | -3.6446276 | | No – ≥95% within clade |
| *HOG1* | 43:657,401 | T>C | 1 | M1V | Missense | III: 3/5568 (0.05%) | -10.091508 | High impact | Yes |
| *HOG1* | 43:657,197 | G>A | 69 | R69* | Stop-gained | IV: 1/1576 (0.06%) | -24.39212 | | Yes |
| *HOG1* | 43:656,763 | TCCT>CCCC | 213 | Complex | Multiresidue | IV: 1576/1576 (100.00%); VI: 6/6 (100.00%) | 0 | | No – complex/in-frame |
| *HOG1* | 43:656,534 | G>A | 290 | P290S | Missense | I: 1/5174 (0.02%) | -9.128215 | | Yes |
| *MKC1* | 38:1,177,422 | G>C | 4 | N4K | Missense | II: 25/40 (62.50%) | -3.6051784 | | Yes |
| *MKC1* | 38:1,177,408 | T>C | 9 | N9S | Missense | IV: 1044/1576 (66.24%) | -1.228461 | | Yes |
| *MKC1* | 38:1,176,620 | A>G | 272 | S272P | Missense | III: 1/5568 (0.02%) | -14.099575 | High impact | Yes |
| *MKC1* | 38:1,176,589 | G>A | 282 | S282F | Missense | II: 40/40 (100.00%) | -7.4421167 | | No – ≥95% within clade |
| *MKC1* | 38:1,176,256 | C>T | 393 | G393E | Missense | V: 3/3 (100.00%) | -3.4500372 | | No – ≥95% within clade |
| *MKC1* | 38:1,176,186 | C>A | 416 | E416D | Missense | II: 40/40 (100.00%) | -2.6109354 | | No – ≥95% within clade |
| *MKC1* | 38:1,176,175 | T>C | 420 | D420G | Missense | I: 4/5174 (0.08%) | -3.9448948 | | Yes |
| *MKC1* | 38:1,176,053 | C>A | 461 | A461S | Missense | V: 3/3 (100.00%) | -2.8860083 | | No – ≥95% within clade |
| *MKC1* | 38:1,175,980 | C>T | 485 | R485H | Missense | II: 1/40 (2.50%) | -3.3234544 | | Yes |
| *CEK1* | 39:972,666 | T>C | 10 | Y10H | Missense | IV: 13/1576 (0.82%) | -3.3561924 | | Yes |
| *CEK1* | 39:972,711 | CACCAGAACCAGA>C | 24 | Del-Non-frameshift-n=12_24 | In-frame indel | III: 29/5568 (0.52%) | 0.34292603 | | No – complex/in-frame |
| *CEK1* | 39:972,719 | CCAGAACCAGAACCAGAACCAGAACCAGAACCAGAACCAGAACCAGAGCCAGAGTCAGC>TCAGAACCAGAACCAGA | 27 | Del-Non-frameshift-n=42_27 | In-frame indel | V: 1/3 (33.33%) | -2.1158524 | | No – complex/in-frame |
| *CEK1* | 39:972,729 | AACCAGAACCAGAACCAGAACCAGAACCAGAACCAGAGCCAGAGTCAGC>A | 30 | Del-Non-frameshift-n=48_30 | In-frame indel | IV: 1520/1576 (96.45%); VI: 5/6 (83.33%) | -2.1766052 | | No – complex/in-frame |
| *CEK1* | 39:972,736 | A>G | 33 | N33S | Missense | II: 1/40 (2.50%) | -7.906288 | | Yes |
| *CEK1* | 39:972,743 | CCAGAACCAGAACCAGAACCAGAGCCAGAGT>C | 35 | Del-Non-frameshift-n=30_35 | In-frame indel | II: 1/40 (2.50%) | -1.6202011 | | No – complex/in-frame |
| *CEK1* | 39:972,747 | AACCAGAACCAGAACCAGAGCCAGAGTCAGC>A | 36 | Del-Non-frameshift-n=30_36 | In-frame indel | III: 1314/5568 (23.60%) | -2.4072723 | | No – complex/in-frame |
| *CEK1* | 39:972,761 | CCAGAGCCAGAGT>C | 41 | Del-Non-frameshift-n=12_41 | In-frame indel | II: 4/40 (10.00%) | -0.99279785 | | No – complex/in-frame |
| *CEK1* | 39:973,078 | C>T | 147 | T147M | Missense | V: 3/3 (100.00%) | -6.070587 | | No – ≥95% within clade |
| *CEK1* | 39:973,143 | C>T | 169 | Q169* | Stop-gained | I: 1/5174 (0.02%) | -24.153805 | | Yes |
| *CEK1* | 39:973,181 | C>G | 181 | Y181* | Stop-gained | III: 1/5568 (0.02%) | -24.153805 | | Yes |
| *CEK1* | 39:973,438 | T>A | 267 | I267K | Missense | I: 1/5174 (0.02%) | -19.213427 | High impact | Yes |
| *CEK1* | 39:973,497 | C>A | 287 | L287I | Missense | I: 5/5174 (0.10%) | -7.427143 | | Yes |
| *CEK1* | 39:973,653 | A>C | 339 | M339L | Missense | I: 21/5174 (0.41%) | -2.0541043 | | Yes |
| *CEK1* | 39:973,689 | C>T | 351 | P351S | Missense | I: 2/5174 (0.04%) | -9.322148 | High impact | Yes |

**Supplementary Table 6: Oligonucleotide inventory and purpose annotations.** Oligonucleotides are grouped by target gene or plasmid backbone. Purpose labels are defined as follows: PCR, amplification; Seq, sequencing; US_cloning and DS_cloning, amplification of upstream and downstream regions for cloning into the pSFS2 backbone; TopGuide and BottomGuide, complementary oligonucleotides used for guide insertion into the pJMR19 backbone after duplexing; dDNA, amplification of donor DNA from the corresponding experimentally evolved isolates; verify deletion, verification of gene deletion by colony PCR; crRNA, CRISPR RNA duplexed with trans-activating CRISPR RNA (tracrRNA) for ribonucleoprotein (RNP) complex assembly; and Deep-seq, amplification of the *FKS1* HS1/HS3 and HS2 regions for amplicon deep sequencing. Combined labels indicate oligonucleotides used for multiple purposes. Primer-specific $T_m$ was calculated with https://tmcalculator.neb.com/.

| Target | Purpose | Oligo name | Sequence |
|---|---|---|---|
| | US_cloning | Cau_sln1_gib_us_rev | TAGAAAGTATAGGAACTTCCTATCGACAATGGAACAGG |
| | US_cloning | Cau_sln1_gib_us_rev | GCGAATTGGAGCTCCACCGCGGTCTCATCATCTGCCTTGAG |
| | DS_cloning | Cau_sln1_gib_ds_fw | CAAAAGCTGGGTACCGGGCCCAACAACATCCTTATGCAC |
| | DS_cloning | Cau_sln1_gib_ds_fw | AGATCCACTAGTTCTAGAGCACTCCCTTTGCCACATCATC |
| *SLN1* | PCR/Seq | Cau_SLN1_DS_check | GCGTTTATGAGGAATCCAGCA |
| | PCR/Seq | Cau_SLN1_DS_nch | GACGGCCACTAAGCTCATACGA |
| | PCR/Seq | Cau_SLN1_US_check | ATCCATGTAATGGCAATGACGCC |
| | PCR/Seq | Cau_SLN1_US_nch | CAGGGCAAACATCTCGGAAGAGG |
| | crRNA | SLN1_crRNA | AACAGCTCCAAAGCCGTCTT |
| | US_cloning | Cau_nik1_gib_us_rev | TAGAAAGTATAGGAACTTCCGGGATGATCTAACCGTGG |
| | US_cloning | Cau_nik1_gib_us_fw | CAAAAGCTGGGTACCGGGCCCGTGAAAGCCCTGAAATTAG |
| | DS_cloning | Cau_nik1_gib_ds_rev | GCGAATTGGAGCTCCACCGCGGAGCCCATACTCGGAGAAAAG |
| | DS_cloning | Cau_nik1_gib_ds_fw | AGATCCACTAGTTCTAGAGCTTCTCTTGAAGACAAGACAAAG |
| *NIK1* | PCR/Seq | Cau_NIK1_DS_check | GGACCCGCTCTATGTATTGG |
| | PCR/Seq | Cau_NIK1_DS_nch | CCGAGCACGAGGCAAAGA |
| | PCR/Seq | Cau_NIK1_US_check | TCAATTCTTCTCGGTGAGCCAATC |
| | PCR/Seq | Cau_NIK1_US_nch | CGCCTGAAGCTGGTCCATCATT |
| | crRNA | NIK1_crRNA | ATTGTCTGCATCAACAACTT |
| | US_cloning | Cau_ssk1_gib_us_rev | TAGAAAGTATAGGAACTTCCAAAATGTGGATGATTGGTG |
| | US_cloning | Cau_ssk1_gib_us_fw | CAAAAGCTGGGTACCGGGCCCACTCTCTTTGGCCAATTG |
| | DS_cloning | Cau_ssk1_gib_ds_rev | GCGAATTGGAGCTCCACCGCGGGAACTCTCCATATGTTAGTCCATG |
| | DS_cloning | Cau_ssk1_gib_ds_fw | AGATCCACTAGTTCTAGAGCAATCAGGGACCTGAACGG |
| *SSK1* | PCR/Seq | Cau_SSK1_DS_check | TCTTCATTGGTTGCCAGGC |
| | PCR/Seq | Cau_SSK1_DS_nch | GTGATCATCGTGGCGCTTACAGC |
| | PCR/Seq | Cau_SSK1_US_check | CGACGATGAAGACGATGATGTGG |
| | PCR/Seq | Cau_SSK1_US_nch | CCTCCTGCTCCAGCACTAGGC |
| | crRNA | SSK1_crRNA | GCATTCACGACAACGCTCGT |
| | US_cloning | Cau_ssk2_gib_us_rev | TAGAAAGTATAGGAACTTCCACTGTAGCCTATTGATTTC |
| | US_cloning | Cau_ssk2_gib_us_fw | CAAAAGCTGGGTACCGGGCCCTCACCTTAGTAGCTTTGATTC |
| | DS_cloning | Cau_ssk2_gib_ds_rev | GCGAATTGGAGCTCCACCGCGGAGAGAAAGAATAAGTCGAGAAAG |
| | DS_cloning | Cau_ssk2_gib_ds_fw | AGATCCACTAGTTCTAGAGCAGTTAAGGAGGGTCATCTAAAAC |
| *SSK2* | PCR/Seq | Cau_SSK2_DS_check | GGTTTGACGTGGTTAAGTCATACAC |
| | PCR/Seq | Cau_SSK2_DS_nch | CTACTGATCAATTGAGCGAAGCAGGC |
| | PCR/Seq | Cau_SSK2_US_check | TATTGCAGTGCGCGGTGAAC |
| | PCR/Seq | Cau_SSK2_US_nch | CTTAAGTATGCCCTCTCTTGAGC |
| | crRNA | SSK2_crRNA | AGACGACCGGCATTATCCAG |
| | US_cloning | Cau_pbs2_gib_us_rev | CAAAAGCTGGGTACCGGGCCCTACTTGCTGCCCACAGTG |
| | US_cloning | Cau_pbs2_gib_us_fw | TAGAAAGTATAGGAACTTCCATGCAATTGGCTGTGAGG |
| | DS_cloning | Cau_pbs2_gib_ds_rev | AGATCCACTAGTTCTAGAGCAGCATACGCTATTTCTACTC |
| | DS_cloning | Cau_pbs2_gib_ds_fw | GCGAATTGGAGCTCCACCGCGGTGCAGAGATCTCGTTAGG |
| | PCR/Seq | Cau_PBS2_DS_check | CACTTGGACAGCTTGTTTCCT |
| | PCR/Seq | Cau_PBS2_DS_nch | CCGATCTCCGCCCATCATATGACG |
| *PBS2* | PCR/Seq | Cau_PBS2_US_check | CACTCACACAAGCTGTCGGCA |
| | PCR/Seq | Cau_PBS2_US_nch | CTGGAATCTGATCTGGCGTG |
| | crRNA | PBS2_crRNA | GGTCTGACGTGAGGTAACGG |
| | TopGuide | Cau_PBS2_pbs2*427Q_top | CCATTTGTGCCTCCTTAGAAAAA |
| | BottomGuide | Cau_PBS2_pbs2*427Q_bottom | AACTTTTTCTAAGGAGGCACAAA |

| Target | Purpose | Oligo name | Sequence |
|---|---|---|---|
| | dDNA | Cau_PBS2_in_F | TACATTGTGGACTTTTACGG |
| | dDNA | Cau_PBS2_in_R | TCAATGGAGCGCTCTTTTTG |
| | PCR/Seq | Cau_PBS2_out_F | GGTACGTCTCGAGTTAGAGG |
| | PCR/Seq | Cau_PBS2_out_R | CTCACTGTACTCGGTGAAAC |
| *HOG1* | US_cloning | Cau_hog1_gib_us_rev | CAAAAGCTGGGTACCGGGCCCGTAGTCTCCACCAATTGTC |
| | US_cloning | Cau_hog1_gib_us_fw | TAGAAAGTATAGGAACTTCCCGAGGAAGTAACGAAGCAG |
| | DS_cloning | Cau_hog1_gib_ds_rev | AGATCCACTAGTTCTAGAGCTGCGTTTGCAGGAGTTTG |
| | DS_cloning | Cau_hog1_gib_ds_fw | GCGAATTGGAGCTCCACCGCGGATTATGGTGGGGTTGATTTG |
| | PCR/Seq | Cau_HOG1_DS_check_v2 | TCAATCTGTGGCCCAAGG |
| | PCR/Seq | Cau_HOG1_DS_nch_v2 | AAGAAGCGAATCTCTGCCG |
| | PCR/Seq | Cau_HOG1_US_nch | CCTGGAGCTCGGTGACA |
| | PCR/Seq | Cau_HOG1_US_check | GGACCCAGCAGAGGGCTTAA |
| | PCR/Seq | Cau_HOG1_US_check_v3 | GGACCCAGCGGAGGGCTTAA |
| | crRNA | HOG1_crRNA | GACCTCAACCCGGTGGGCAT |
| *MKC1* | US_cloning | Cau_mkc1_gib_us_rev | CTAACGATGCATACGACTACATC |
| | US_cloning | Cau_mkc1_gib_us_fw | CCCCAGGCTTTACACTTTATG |
| | DS_cloning | Cau_mkc1_gib_ds_rev | GCAAGGCGATTAAGTTGGGTAAC |
| | DS_cloning | Cau_mkc1_gib_ds_fw | ACATATGTGAAGTGTGAAGGGGG |
| | PCR/Seq | Cau_MKC1_DS_check_v2 | GACGAGGAACTTCCTTCAATCC |
| | PCR/Seq | Cau_MKC1_DS_nch | CCATACATGTCTGTTTGGCATGACC |
| | PCR/Seq | Cau_MKC1_US_check | TTACTTATCGCCGTACGCATGG |
| | PCR/Seq | Cau_MKC1_US_nch | GGGACAATGTCCAAGTCGTACAA |
| | crRNA | MKC1_crRNA | ATCGTCAAGGAGATTGGTCA |
| *CEK1* | US_cloning | Cau_cek1_gib_us_fw | CAAAAGCTGGGTACCGGGCCCATATCGAGACCCAAAAGATCCAAG |
| | US_cloning | Cau_cek1_gib_us_rev | TAGAAAGTATAGGAACTTCCCTGCTGGGAAAACTGCGTG |
| | DS_cloning | Cau_cek1_gib_ds_fw | AGATCCACTAGTTCTAGAGCACCACGGGCCTTGC |
| | DS_cloning | Cau_cek1_gib_ds_rev | GCGAATTGGAGCTCCACCGCGGCATTAGGAGAAATGATGTAGAAGTCAATG |
| | PCR/Seq | Cau_CEK1_DS_check | GAAGCAGTACTTGTATCTGCCG |
| | PCR/Seq | Cau_CEK1_DS_nch | GAAGGTGCCATTTGCAGAGC |
| | PCR/Seq | Cau_CEK1_US_check | CCAGCCCGACCTGATAATACCAC |
| | PCR/Seq | Cau_CEK1_US_nch | CTGGTTCTGGTGCTGGATCTGC |
| | crRNA | CEK1_crRNA | CAAGCCGTGTTTCAGGTTAG |
| pSFS2 | PCR/Seq/verify deletion | pSAT1_US-cPCR_F | CTAACGATGCATACGACTACATC |
| | PCR | pSAT1_US-cPCR_R | CCCCAGGCTTTACACTTTATG |
| | PCR | pSAT1_DS-cPCR_R | GCAAGGCGATTAAGTTGGGTAAC |
| | PCR/Seq/verify deletion | pSAT1_DS-cPCR_F | ACATATGTGAAGTGTGAAGGGGG |
| pJMR19 | Seq | seq_gRNA_pJMR19 | GGGTGTCGGTTGGGTTGTG |
| *FKS1* HS1/HS3 | Deep-seq | CLOP372-A6_fw1 | TCGTCGGCAGCGTCAGATGTGTATAAGAGACAGNTTGGGTCACTGTGTTTGCTG |
| | Deep-seq | CLOP372-B6_fw2 | TCGTCGGCAGCGTCAGATGTGTATAAGAGACAGNNTTGGGTCACTGTGTTTGCTG |
| | Deep-seq | CLOP372-E6_rev1 | GTCTCGTGGGCTCGGAGATGTGTATAAGAGACAGNAGAACGAACGACCAATGGAG |
| | Deep-seq | CLOP372-F6_rev2 | GTCTCGTGGGCTCGGAGATGTGTATAAGAGACAGNNAGAACGAACGACCAATGGAG |
| *FKS1* HS2 | Deep-seq | CLOP372-G6_fw1 | TCGTCGGCAGCGTCAGATGTGTATAAGAGACAGNTGTGTGACTACGACAGAAAC |
| | Deep-seq | CLOP372-H6_fw2 | TCGTCGGCAGCGTCAGATGTGTATAAGAGACAGNNTGTGTGACTACGACAGAAAC |
| | Deep-seq | CLOP358-H11_rev1 | GTCTCGTGGGCTCGGAGATGTGTATAAGAGACAGNGTAGCTTTCCAAAGACCTCT |
| | Deep-seq | CLOP358-A12__rev2 | GTCTCGTGGGCTCGGAGATGTGTATAAGAGACAGNNGTAGCTTTCCAAAGACCTCT |

**Supplementary Table 7: Acquisition and processing parameters for two-dimensional solid-state NMR experiments.** Parameters are reported for wt and *hog1Δ C. auris* cells grown under drug-free conditions or with 2 μg $mL^{-1}$ CAS. DARR experiments characterised comparatively rigid carbohydrate components, whereas DP-INADEQUATE experiments characterised comparatively mobile components. HC-CP denotes $^{1}H$–$^{13}C$ cross-polarisation, DP denotes direct polarisation and DARR, dipolar-assisted rotational resonance. SPINAL64 denotes the $^{1}H$ decoupling sequence. Radiofrequency field amplitudes are expressed in kHz and $^{13}C$ carrier positions in ppm. The indirect and direct acquisition dimensions are designated $t_1$ and $t_2$, respectively. Apodisation settings, interscan delays, scan numbers and total measurement times are indicated.

| Sample | wt | wt + cas | hog1Δ | hog1Δ + cas | wt | wt + cas | hog1Δ | hog1Δ + cas |
|---|---|---|---|---|---|---|---|---|
| Experiment | 2D 20 ms DARR (rigid) | | | | 2D DP INADEQUATE (flexible) | | | |
| Transfer I | HC-CP | HC-CP | HC-CP | HC-CP | $^{13}C$ DP | $^{13}C$ DP | $^{13}C$ DP | $^{13}C$ DP |
| $^{1}H$ field/kHz | 59 | 59 | 59 | 59 | | | | |
| $^{15}N$ field/kHz | | | | | | | | |
| $^{13}C$ field/kHz | 49 | 49 | 49 | 37 | | | | |
| Shape | ramp $^{1}H$ | ramp $^{1}H$ | ramp $^{1}H$ | ramp $^{1}H$ | | | | |
| Time/ms | 0.5 | 0.5 | 0.5 | 0.5 | 2.5 | 2.5 | 2.5 | 2.5 |
| $^{13}C$ carrier/ppm | 104 | 104 | 104 | 104 | 104 | 104 | 104 | 104 |
| Transfer II | DARR | DARR | DARR | DARR | INADEQUATE | INADEQUATE | INADEQUATE | INADEQUATE |
| $^{1}H$ field/kHz | 12 | 12 | 12 | 12 | | | | |
| $^{13}C$ field/kHz | | | | | | | | |
| Shape | | | | | | | | |
| Time/ms | 20 | 20 | 20 | 20 | | | | |
| $^{1}H$ decoupling | SPINAL64 | SPINAL64 | SPINAL64 | SPINAL64 | SPINAL64 | SPINAL64 | SPINAL64 | SPINAL64 |
| $^{1}H$ decoupling field/ kHz | 79 | 79 | 79 | 79 | 68 | 68 | 68 | 76 |
| t1 increments | 500 | 500 | 500 | 500 | 480 | 480 | 480 | 360 |
| Windows function | QSine 3 | QSine 3 | QSine 3 | QSine 3 | QSine 3 | QSine 3 | QSine 3 | QSine 3 |
| Sweep width (t1)/kHz | 40 | 40 | 40 | 40 | 72 | 72 | 72 | 68 |
| Acquisition time (t1)/ms | 6.2 | 6.2 | 6.2 | 6.2 | 6.6 | 6.6 | 6.6 | 5.2 |
| t2 increments | 1536 | 1536 | 1536 | 1536 | 1434 | 1434 | 1434 | 1434 |
| Windows function | QSine 3 | QSine 3 | QSine 3 | QSine 3 | QSine 3 | QSine 3 | QSine 3 | QSine 3 |
| Sweep width (t2)/kHz | 60 | 60 | 60 | 60 | 60 | 60 | 60 | 60 |
| Acquisition time (t2)/ms | 12.9 | 12.9 | 12.9 | 12.9 | 12 | 12 | 12 | 12 |
| Interscan delay/s | 3 | 3 | 3 | 3 | 3 | 3 | 3 | 3 |
| Number of scans | 64 | 64 | 64 | 64 | 64 | 64 | 64 | 64 |
| Measurement time/h | 27 | 27 | 27 | 27 | 25 | 25 | 25 | 24 |

**Supplementary Table 8: Population-model parameters, units and sources.** Shared settings, strain-specific concentration-growth fits and empirical colony-forming unit (CFU) inputs used in the stochastic population model. Simulations followed separate wt, *hog1Δ* and *Nik1*$^{E472K}$ parental cohorts, each with the possibility of an emerging Fks1$^{mut}$ lineage, in one well-mixed compartment. Fks1$^{mut}$ parameters are represented by measurements of the Fks1$^{S639Y}$ resistant phenotype. Parent and resistant populations share a carrying capacity with turnover. MCF concentrations declined mono-exponentially between daily doses and were reset to the specified peak at each dose. Drug-exposure and immune-response settings were based on the literature[7,8]. The remaining settings and empirical inputs originate from this study. Growth fits were obtained from measurements in RPMI containing 2% glucose and MCF. Empirical CFU inputs governed the first 24 h, comprising the initial decline from 0 to 0.5 h and recovery from 0.5 to 24 h, after which the concentration–growth fits governed growth. *Nik1*$^{E472K}$ retained its own growth fit but used the *hog1*Δ CFU profile as a proxy. The CFU values listed here correspond to the initial MCF peak of 16 μg ml$^{-1}$. Positive additional-decline values denote extra decline relative to the drug-free control, but recovery differences retain their sign relative to that control. Immune-mediated killing had equal per-capita effects across genotypes. Each parental background was evaluated in 500 simulations. Main endpoint summaries show the Wilson 95% confidence intervals.

| Parameter / setting | Symbol / code | Value / setting | Origin |
|---|---|---|---|
| **A. Initial conditions and population representation** | | | |
| Initial parental population | $N_p(0)$ | 200000 cells | This study |
| Initial resistant population | $N_r(0)$ | 0 cells | |
| Blood-equivalent volume | V | 5000 mL | |
| Initial blood-equivalent burden | $N_p(0)/V$ | 40 CFU ml$^{-1}$ | |
| Shared carrying capacity | K | 200000000 cells | |
| Resistant phenotype proxy | $FKS1_{mut}$ | Fks1$^{S639Y}$ | |
| **B. Micafungin exposure** | | | |
| Concentration immediately after each dose | $C_{peak}$ | 16 μg ml$^{-1}$ | 9 |
| Dose interval | T | 24 h | |
| Micafungin elimination half-life | $t_{½}$ | 15 h | |
| **C. Empirical growth and first-day CFU inputs** | | | |
| Growth-rate measurement window | - | 24 h | This study |
| Concentration–growth response | - | Base four-parameter Emax/Hill | |
| Empirical CFU layer active until | $t_{CFU}$ | 24 h after initial exposure | |
| Initial CFU decline window | - | 0–0.5 h | |
| Initial CFU recovery window | - | 0.5–24 h | |
| **D. Resistance emergence and density regulation** | | | |
| Mutation probability per parental division opportunity | $p_{mut}$ | 5 × 10$^{-10}$ per cell division | This study |
| **E. Dynamic immune-mediated clearance** | | | |
| Initial immune-effector density | E(0) | 100 effector cells ml$^{-1}$ | 7,8 |
| Immune-effector reservoir parameter | $E_{max}$ | 1.0 × 10$^{6}$ effector cells ml$^{-1}$ | |
| Immune-effector recruitment rate coefficient | q | 0.001 h$^{-1}$ | |
| Immune-effector decay coefficient | l | 0.001 h$^{-1}$ | |
| Immune–fungal interaction coefficient | j | 1.2 × 10$^{-6}$ ml effector-cell$^{-1}$ h$^{-1}$ | |
| **F. Numerical and reporting settings** | | | |
| Simulation duration | t(end) | 14 days | This study |
| Maximum simulation step | $\tau$(max) | 0.1 h | |
| Independently seeded model runs | n(runs) | 5 runs | |
| Simulations per run and parental background | n(sim) | 100 simulations | |

| Parameter/input | Symbol | wt | *hog1Δ* | Nik1$^{E472K}$ | Fks1$^{mut}$ | Origin |
|---|---|---|---|---|---|---|
| **G. Strain-specific empirical input values** | | | | | | |
| Drug-free apparent growth-rate parameter | $k_g$ (h$^{-1}$) | 0.490 | 0.514 | 0.538 | 0.532 | This study |
| Maximum fractional growth inhibition | $E_{max}$ | 0.922 | 0.514 | 0.490 | 0.253 | |
| Half-maximal inhibitory concentration | $EC_{50}$ (μg ml$^{-1}$) | 0.160 | 0.027 | 0.143 | 3.418 | |
| Hill coefficient | h | 1.843 | 0.832 | 0.492 | 8.383 | |

| | | | | | | |
|---|---|---|---|---|---|---|
| **H. CFU inputs at the specified initial micafungin peak (16 μg ml$^{-1}$)** | | | | | | |
| Additional early decline relative to drug-free control | L ($\log_{10}$ fold change) | 0.960 | 1.374 | 1.374 | 0.375 | This study |
| Recovery change relative to drug-free control | $\Delta$rec ($\log_{10}$ fold change) | -0.330 | 1.141 | 1.141 | 0.375 | |
| Drug-free control early change | $\Delta$ctrl,early ($\log_{10}$ fold change) | -0.907 | -1.276 | -1.276 | -0.827 | |
| Drug-free control recovery change | $\Delta$ctrl,rec ($\log_{10}$ fold change) | 2.611 | 2.975 | 2.975 | 2.571 | |
